# Breastmilk-promoted bifidobacteria produce aromatic amino acids in the infant gut

**DOI:** 10.1101/2020.01.22.914994

**Authors:** Martin F. Laursen, Mikiyasu Sakanaka, Nicole von Burg, Urs Mörbe, Daniel Andersen, Janne Marie Moll, Ceyda T. Pekmez, Aymeric Rivollier, Kim F. Michaelsen, Christian Mølgaard, Mads Vendelbo Lind, Lars O. Dragsted, Takane Katayama, Henrik L. Frandsen, Anne Marie Vinggaard, Martin I. Bahl, Susanne Brix, William Agace, Tine R. Licht, Henrik M. Roager

**Author notes:** These authors contributed equally (Martin F. Laursen, Henrik M. Roager). **Correspondence** Henrik M. Roager, Rolighedsvej 30, 1958 Frederiksberg C, Denmark, Tine R. Licht, Kemitorvet, 2800 Kongens Lyngby, Denmark.

## Abstract

Breastfeeding profoundly shapes the infant gut microbiota, which is critical for early life immune development. However, few breastmilk-dependent microbial metabolites mediating host-microbiota interactions are currently known. We here demonstrate that breastmilk-promoted *Bifidobacterium* species convert aromatic amino acids (tryptophan, phenylalanine and tyrosine) into their respective aromatic lactic acids (indolelactate, phenyllactate and 4-hydroxyphenyllactate) via a previously unrecognised aromatic lactate dehydrogenase. By longitudinal profiling of the gut microbiota composition and metabolome of stool samples of infants obtained from birth until 6 months of age, we show that stool concentrations of aromatic lactic acids are determined by the abundance of human milk oligosaccharide degrading *Bifidobacterium* species containing the aromatic lactate dehydrogenase. We demonstrate that stool concentrations of *Bifidobacterium*-derived indolelactate, the most abundant aromatic lactic acid *in vivo*, are associated with the capacity of infant stool samples to activate the aryl hydrocarbon receptor (AhR), a receptor important for controlling intestinal homeostasis and immune responses. Finally, we show that indolelactate modulates *ex vivo* immune responses of human CD4+ T-cells and monocytes in a dose-dependent manner by acting as an agonist of both, the AhR and hydroxycarboxylic acid receptor 3 (HCAR3). Our findings reveal that breastmilk-promoted *Bifidobacterium* produce aromatic lactic acids in the gut of infants and suggest that these microbial metabolites may impact immune function in early life.

## INTRODUCTION

Human breastmilk is a perfectly adapted nutritional supply for the infant^1^. Breastfeeding provides children with important short-term protection against infections, and may also provide long-term metabolic benefits^1,2^. These benefits may partly be mediated through the gut microbiota, since breastfeeding is the strongest determinant of gut microbiota composition and function during infancy^3–5^. Human breastmilk contains human milk oligosaccharides (HMOs), which are complex, highly abundant sugars serving as substrates for specific microbes including certain species of *Bifidobacterium*^6^. This co-evolution between bifidobacteria and the host, mediated by HMOs, to a large extent directs the colonisation of the gut in early life, which has critical impact on the immune system^7^. Depletion of specific microbes, including *Bifidobacterium,* in early life has been associated with increased risk of allergy and asthma development in childhood^8,9^, and is suggested to compromise immune function and lead to increased susceptibility to infectious disease^10,11^. Despite *Bifidobacterium* dominating the gut of breastfed infants and being widely acknowledged as beneficial, mechanistic insights on the contribution of these bacteria and their metabolites to immune development during infancy remain limited. Recent studies show that microbial aromatic amino acid metabolites including tryptophan-derived indoles^12^ via activation of the aryl hydrocarbon receptor (AhR)^13,14^ can fortify the intestinal barrier^15,16^, protect against pathogenic infections^13,17^ and influence host metabolism^15,18,19^, making this group of microbial metabolites of particular interest in the context of early life.

Here, we show that breastmilk-promoted *Bifidobacterium* species, via a previously unrecognised aromatic lactate dehydrogenase, produce aromatic lactic acids including indolelactate (ILA) in substantial amounts in the infant gut, and that stool concentrations of this metabolite are correlated with the capacity of infant stool to activate AhR. We furthermore demonstrate that ILA via AhR and HCAR3-dependent pathways impact immune functions *ex vivo*, suggesting that breastmilk-promoted *Bifidobacterium* via production of aromatic lactic acids impact the immune system in early life.

## RESULTS

### *Bifidobacterium* species associate with aromatic amino acid catabolites during late infancy

To explore interactions between breastfeeding status, gut microbial composition and metabolism of aromatic amino acid in early life, we employed 16S rRNA amplicon sequencing to infer gut microbiota composition and a targeted ultra-performance liquid chromatography mass spectrometry (UPLC-MS) metabolomics approach to quantify 19 aromatic amino acids and derivatives thereof (**Supplementary Table 1**) in stool samples from 59 healthy Danish infants from the SKOT I cohort^20^. The SKOT I infants included were born full term, 9.1 ± 0.3 (mean ± SD) months of age at sampling, and 40.7% were still partially breastfed (**Supplementary Data 1a,b**). After stratification of the 9 months old infants based on breastfeeding status (partially breastfed versus weaned), Principal Coordinates Analysis (PCoA) of weighted UniFrac distances showed a significant separation across the first PC-axis (r^2^ = 0.093, p < 0.001, Adonis test; **Fig. 1a**), which was due to an increasing gradient in relative abundance of *Bifidobacterium* in breastfed infants (r^2^ = 0.397, p < 0.001, Adonis test; **Fig. 1b**). Other metadata (age, gender, mode of delivery, current formula intake and age of introduction to solid foods) did not explain gut microbiota variation to the same degree as breastfeeding status (**Supplementary Data 1c,d**). Principal Component Analysis (PCA) of faecal aromatic amino acid metabolite concentrations also revealed separation according to breastfeeding status, which was largely driven by three aromatic lactic acids, 4-hydroxyphenyllactic acid (4-OH-PLA), phenyllactic acid (PLA) and indolelactic acid (ILA) (**Fig. 1c**). Correlation analysis revealed that *Bifidobacterium*, but no other bacterial genera, were significantly associated with faecal concentrations of all three aromatic lactic acids (4-OH-PLA, PLA and ILA), in addition to indolealdehyde (IAld) (**Fig. 1d** and **Supplementary Data 1e**). *Bifidobacterium* species (**Supplementary Fig. 1a** and **Supplementary Data 1f**) significantly enriched in the breastfed infants; *B. longum, B. bifidum*, and *B. breve*, were positively associated with the faecal concentrations of aromatic lactic acids (4-OH-PLA, PLA and ILA) and IAld (cluster 1 in **Fig. 1e**), but negatively associated with the faecal concentrations of aromatic propionic acids, aromatic amino acids and to a lesser degree with aromatic acetic acids (cluster 2 in **Fig. 1e**). In contrast, post weaning type *Bifidobacterium* species, including *B. adolescentis*, *B. animalis/pseudolongum* and *B. catenulatum* group^21,22^, were not significantly associated with aromatic lactic acids nor breastfeeding status (**Fig. 1e**). These associations were in agreement with the observation that the concentrations of the three aromatic lactic acids were higher in the faeces of breastfed than in weaned infants (**Supplementary Fig. 1b**). Furthermore, the abundances of the three aromatic lactic acids in infant urine (**Supplementary Fig. 2-4**) showed similar positive associations with relative abundances of breastmilk-promoted *Bifidobacterium* species (**Supplementary Fig. 1c**). In addition, faecal and urinary levels of ILA were positively correlated (rho = 0.68, p < 0.0001), showing that faecal levels of this metabolite are reflected systemically. Consistently, urine abundance of ILA, but not of PLA and 4-OH-PLA, were significantly higher in breastfed compared to weaned infants (**Supplementary Fig. 1b**). Together, this suggests that specific breastmilk-promoted *Bifidobacterium* species in the gut of infants convert aromatic amino acids to the corresponding aromatic lactic acids (**Fig. 1f**).

**Figure 1.**
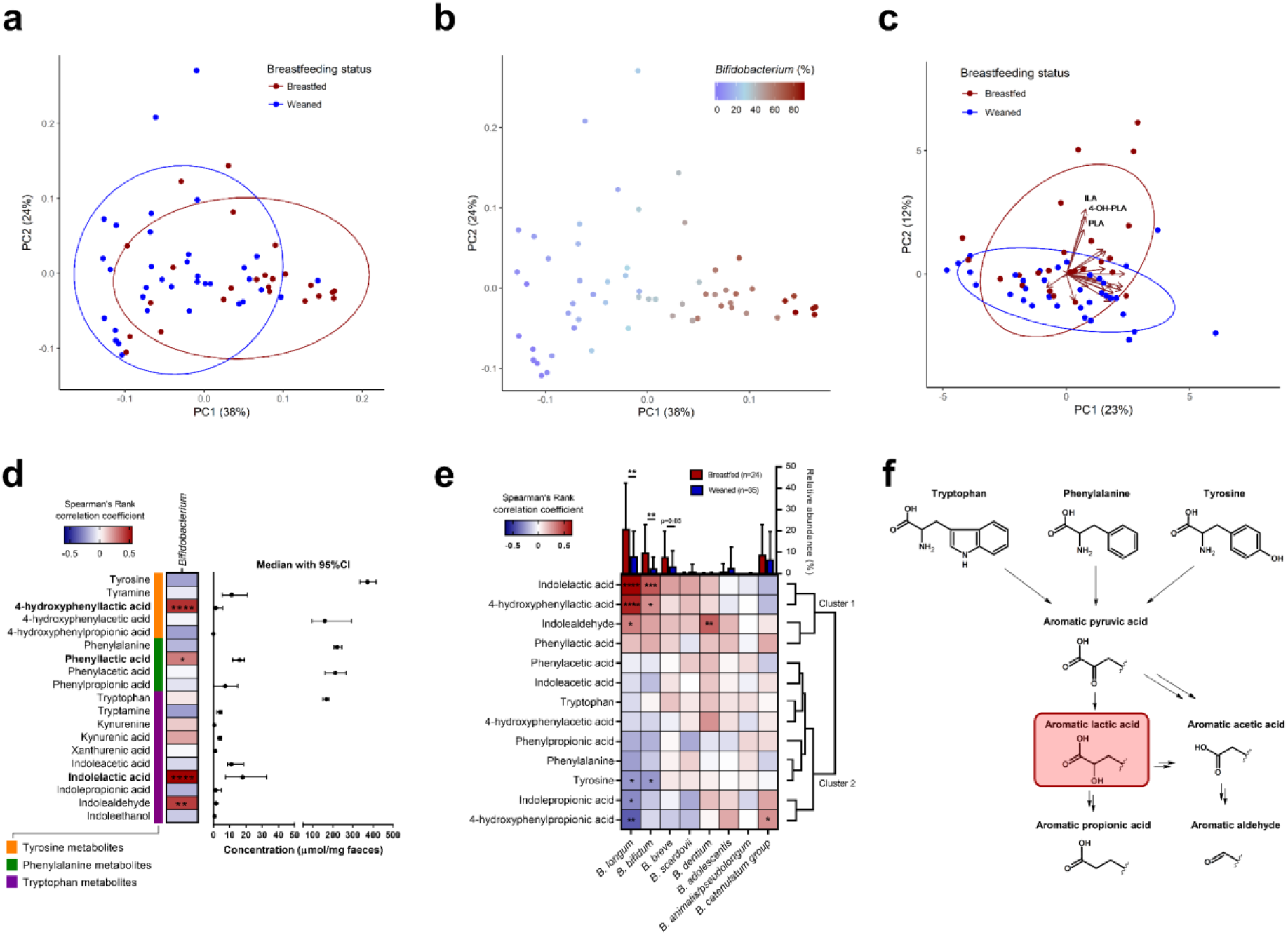
Breastfeeding associates with gut microbiota composition and aromatic amino acid catabolism in 9 months old infants. **a-b**, Principal coordinate analysis plots of weighted UniFrac distances based on OTUs from faecal samples of 9 months old infants participating in the SKOT cohort (n = 59). Samples are coloured according to (a) breastfeeding status with ellipses indicating 80%CI of data points for partially breastfed (red, n = 24) and weaned (blue, n = 35) infants or (b) relative abundance of the genus *Bifidobacterium.* **c**, Principal component analysis plot of concentrations (μmol/mg faeces) of aromatic amino acids and their catabolites in SKOT faecal samples, coloured according to breastfeeding status with ellipses indicating 80%CI of data points for breastfed (red, n = 24) and weaned (blue, n = 35) infants. Loadings are shown with arrows, with annotations of the aromatic amino acids Indolelactic acid (ILA), 4-hydroxyphenyllactic acid (4-OH-PLA) and Phenyllactic acid (PLA) shown. **d**, Heatmap illustrating Spearman’s Rank correlation coefficients between the relative abundance of *Bifidobacterium* and concentrations of aromatic amino acids and their catabolites in SKOT faecal samples (n=59). The concentration of each metabolite (μmol/mg faeces) is presented as the median with 95%CI. **e**, Heatmap illustrating hierarchical clustering of Spearman’s Rank correlation coefficients between the relative abundance of the different *Bifidobacterium* species and selected microbial-derived aromatic amino acid catabolites. Bar plots are showing relative abundance (mean+SD) of the *Bifidobacterium* spp., stratified according to breastfeeding status, with statistical significance evaluated by Mann-Whitney *U* test. **f**, The pathway of aromatic amino acid catabolism by gut microbes (modified from Smith and Macfarlane 1996^107^, Smith and Macfarlane 1997^108^, and Zelante *et al*. 2013^13^). Asterisks indicate statistical significance: * p<0.05, ** p<0.01, *** p<0.001, **** p<0.0001. See also **Supplementary Figure 1-4**.

### *Bifidobacterium* species produce aromatic lactic acids *in vitro*

To confirm the ability of *Bifidobacterium* species detected in infants to produce aromatic lactic acids, *Bifidobacterium* type strains were grown anaerobically in a medium containing all three aromatic amino acids with either glucose or HMOs as sole carbohydrate sources. Analyses of culture supernatants revealed that ILA, PLA and 4-OH-PLA were produced mainly by *B. bifidum, B. breve, B. longum* subsp*. longum, B. longum* subsp*. infantis* and *B. scardovii* (**Fig. 2a**), in accordance with the associations observed in the 9 months old infants (**Fig. 1e**). Other *Bifidobacterium* species, namely *B. adolescentis, B. animalis* subsp*. lactis, B. animalis* subsp*. animalis, B. dentium, B. catenulatum, B. pseudocatenulatum, B. pseudolongum* subsp*. pseudolongum* produced only low amounts of these metabolites **(Fig. 2a**). The ability of *Bifidobacterium* species to produce high levels of the aromatic lactic acids was generally convergent with the ability to utilise HMOs as a carbohydrate source (**Fig. 2a**), supporting the link between breastmilk-promoted bifidobacteria and production of aromatic lactic acids. None of the downstream products of the aromatic lactic acids (**Fig. 1f**) were detected in any of the culture supernatants.

**Figure 2.**
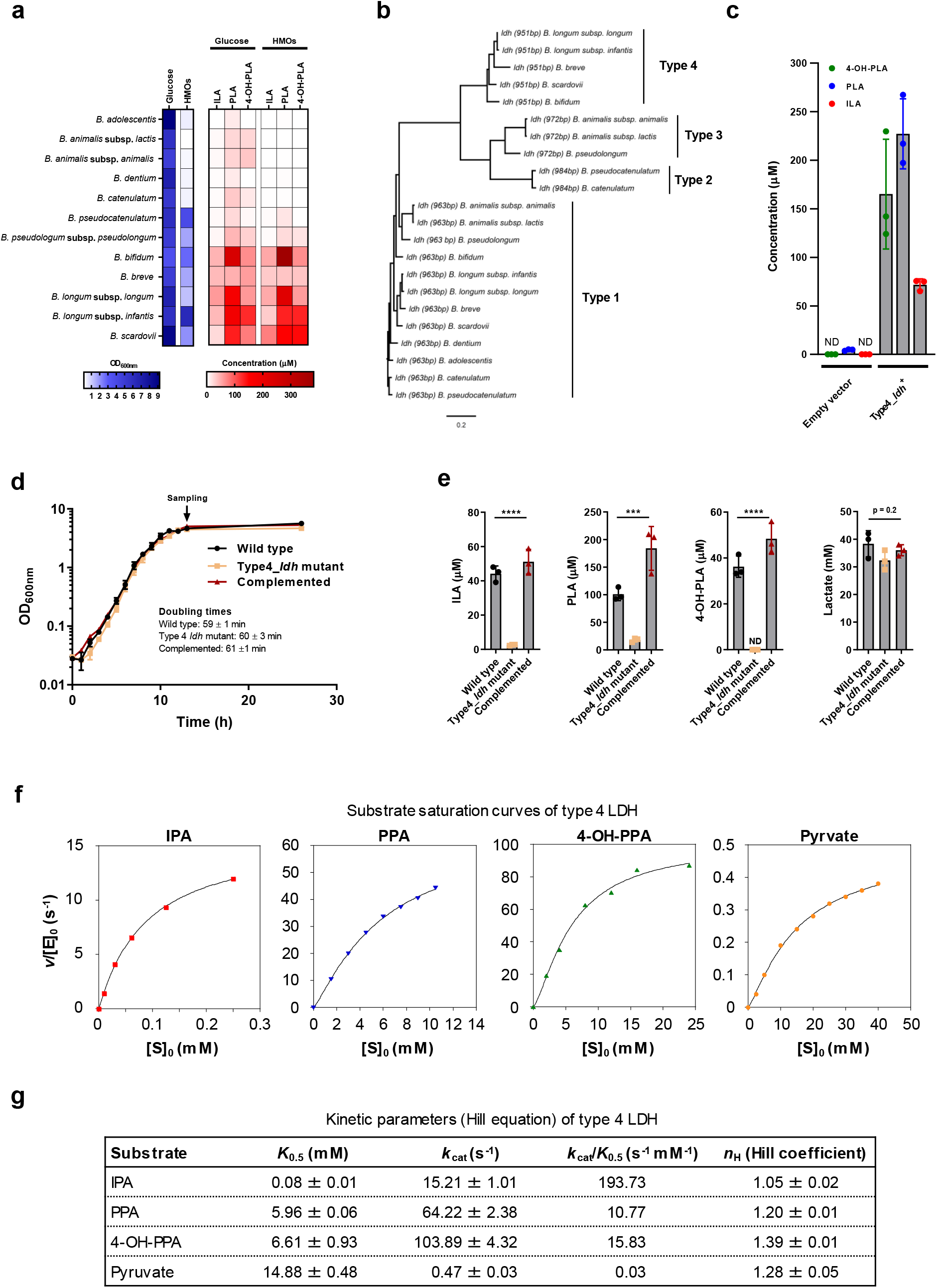
*Bifidobacterium* species produce aromatic lactic acids via an aromatic lactate dehydrogenase. **a**, *In vitro* production of indolelactic acid (ILA), phenyllactic acid (PLA) and 4-hydroxyphenyllactic acid (4-OH-PLA) by *Bifidobacterium* spp. type strains in modified MRS medium (MRSc) with 2% (w/v) glucose or a mix human milk oligosaccharides as sole carbohydrate source. For the type strains of *B. adolescentis*, *B. animalis* subsp. *animalis*, *B. animalis* subsp. *lactis, B. dentium* and *B. catenulatum* no or very poor growth (OD_600nm_ < 0.4) was observed with HMOs as carbohydrate source. Mean of three biological replicates is shown. **b**, Neighbor-Joining phylogenetic tree of all genes in the *Bifidobacterium* spp. type strains annotated as L-lactate dehydrogenases (*ldh*). The four clusters are designated type 1-4. **c**, Production of ILA, PLA and 4-OH-PLA by *E. coli* LMG194 cells transformed with an inducible vector lacking (empty vector) or containing the type 4 *ldh* (Type4_*ldh*+) from *B. longum* subsp. *infantis* DSM 20088T in LB-medium 5 h post-induction of gene expression by addition of L-arabinose and supplementation with the aromatic pyruvic acids (indolepyruvate, phenylpyruvate and 4-hydroxyphenylpyruvate). Bars show mean ± SD of three biological replicates. **d**, Growth curves of *Bifidobacterium longum* subsp. *longum* 105-A (wild type), its isogenic insertional type 4 *ldh* mutant (Type4_*ldh* mutant) and the type 4 *ldh* mutant strain complemented with the type 4 *ldh* gene (Complemented). Curves show mean ± SD of three biological replicates and doubling times reported as mean ± SD. **e**, Production of ILA, PLA, 4-OH-PLA and lactate by wild type, Type4_*ldh* mutant and the complemented strain in early stationary phase cultures. Bars show mean ± SD of three biological replicates. Statistical significance was evaluated by one-way ANOVA, with * p<0.05, ** p<0.01, *** p<0.001, **** p<0.0001. **f-g**, Enzyme kinetics of the type 4 lactate dehydrogenase. **f**, substrate saturation curves of type 4 LDH with indole pyruvate (IPA), phenylpyruvate (PPA), 4-hydroxyphenylpyruvate (4-OH-PPA) or pyruvate as substrates. Data is one representative of two independent assays. **g**, kinetic parameters of type 4 LDH with the different substrates. Data are mean ± SD of two independent assays. ND, not detected See also **Supplementary Figure 5-11**.

### An aromatic lactate dehydrogenase is responsible for the aromatic lactic acid production

Since it has been reported that an L-lactate dehydrogenase (LDH) in *Lactobacillus* spp. can convert phenylpyruvic acid to PLA^23^, we hypothesised that a corresponding enzyme was present in *Bifidobacterium* species. Alignment and phylogenetic analysis of all genes annotated as *ldh* in the *Bifidobacterium* type strains included in this study, revealed four clusters (**Fig. 2b**). Whereas all *Bifidobacterium* genomes contain an *ldh* responsible for conversion of pyruvate to lactate (here designated as type 1 *ldh*) in the bifidobacterial fructose-6-phosphate shunt^24,25^, some species have an extra *ldh*, here designated as type 2, type 3 and type 4, respectively. In agreement with the *in vitro* fermentations (**Fig. 2a**), all prominent aromatic lactic acid-producing *Bifidobacterium* species contain the type 4 *ldh*, suggesting that this could encode a previously unrecognised aromatic lactate dehydrogenase (ALDH). A further analysis of all available whole genome sequenced *Bifidobacterium* strains showed that the type 4 *ldh* is universally present in *B. longum*, *B. bifidum*, *B. breve* and *B. scardovii* strains (**Supplementary Table 2**). Interestingly, genomic analysis of the *Bifidobacterium* type strains revealed that the type 4 *ldh* gene is part of a genetic element containing an amino acid transaminase gene (suspected to be responsible for converting the aromatic amino acids into aromatic pyruvic acids) and a haloacid dehydrogenase gene (of unknown importance) (**Supplementary Fig. 5**), which has been indicated to constitute an operon in *B. breve*^26^. Cloning of the type 4 *ldh* gene from *B. longum* subsp*. infantis*^T^ into a vector transformed into *E. coli* revealed that the expression of the type 4 *ldh* gene indeed resulted in the appearance of PLA, 4-OH-PLA and ILA in the culture supernatant (**Fig. 2c**). To verify the type 4 *ldh* dependent production of aromatic lactic acids in *Bifidobacterium* species, we generated a type 4 *ldh* insertional mutant strain by homologous recombination in *B. longum* subsp. *longum* 105-A (**Supplementary Fig. 6**), a genetically tractable strain containing the type 4 LDH (**Supplementary Fig. 7**)^27,28^. Cultivation of the wild-type (WT), the type 4 *ldh* mutant strain and a complemented type 4 *ldh* mutant strain in a medium containing the three aromatic amino acids (**Supplementary Fig. 6**) confirmed that type 4 *ldh* disruption did not impair growth (**Fig. 2d**). ILA, PLA and 4-OH-PLA accumulated in the supernatant of the WT and of the complemented type 4 *ldh* mutant strains, but not in the type 4 *ldh* mutant (**Fig. 2e**). Importantly, the type 4 *ldh* mutant was not significantly compromised in its ability to convert pyruvate to lactate (**Fig. 2e**), supporting the distinct role of type 4 *ldh* in converting aromatic pyruvic acids. Further, to demonstrate *in vivo* production of the indicated aromatic lactic acids, we mono-colonised germ free mice with either the WT or the type 4 *ldh* mutant strain and found a 20-60 fold increase in their concentrations in WT versus type 4 *ldh* mutant mono-colonised mice (**Supplementary Fig. 8**). Purification and characterisation of the recombinant type 4 LDH enzyme revealed that it had a mass of 33.9 kDa (**Supplementary Fig. 9a**), while the native molecular mass was estimated to be 72.9 kDa by size exclusion chromatography, indicating dimer formation in solution **(Supplementary Fig. 9b**). No metal requirement was observed, the optimal pH was 8.0-8.5 and the enzyme was most stable at 37°C (**Supplementary Fig. 9c-e**). Heterotrophic effects were neither observed for fructose-1,6-bisphosphate (an allosteric effector for type 1 LDH) nor for several intermediates for aromatic amino acid synthesis^24,25^ (**Supplementary Fig. 10**). However, we found that phosphate served as a positive effector suggesting that type 4 LDH is an intracellular enzyme (**Supplementary Fig. 11a-c**). Assay at the different phosphate concentrations revealed the type 4 LDH is a *K*-type allosteric enzyme (**Supplementary Fig. 11b,c**). The catalytic rate (*k*_cat_) was moderate to high for the aromatic pyruvic acid substrates, but very low for pyruvate (**Fig. 2f,g**), in accordance with the non-impaired lactate production observed for the type 4 *ldh* mutant (**Fig. 2e**). Production of ILA, PLA and 4-OH-PLA from the respective aromatic pyruvic acid substrates was verified by high-performance liquid chromatography (HPLC) (**Supplementary Fig. 11d**). The enzyme showed highest affinity (lowest *K*_0.5_) for indole pyruvic acid, but highest catalytic rate for 4-OH-phenyl pyruvic acid in the presence of 100 mM phosphate (**Fig. 2f,g**). However, the catalytic efficiency (*k*_cat_/*K*_0.5_) was highest for indole pyruvic acid (194 s^−1^ mM^−1^), followed by 4-OH-phenyl pyruvic acid (16 s^−1^ mM^−1^) and phenyl pyruvic acid (11 s^−1^ mM^−1^), suggesting preference for indole pyruvic acid. The observed Hill coefficient (*n*_H_ = 1-1.4) for all substrates indicate weak positive cooperativity under the conditions tested. Collectively, these results show that the type 4 *ldh* gene encodes an aromatic lactate dehydrogenase responsible for the production of ILA, PLA and 4-OH-PLA in *Bifidobacterium* species associated with breastfeeding. We therefore suggest that the type 4 *ldh* gene should be re-classified as an aromatic lactate dehydrogenase gene (from now on denoted *aldh*).

### *Bifidobacterium* species govern aromatic lactic acid profiles during early infancy

To study the dynamics of *Bifidobacterium* species establishment and aromatic lactic acids in infants, we established the Copenhagen Infant Gut (CIG) cohort including 25 healthy breast- or mixed fed infants, which were sampled every 2-4 weeks from birth until the age of 6 months (**Supplementary Data 2a**) for microbiome profiling and targeted metabolite quantification including aromatic lactic acids (**Supplementary Fig. 12** and **Supplementary Table 3**). A total of 145 operational taxonomic units (OTUs) were detected by 16S rRNA amplicon sequencing, however, collapsing of OTUs with identical taxonomic classifications and using a cut-off of average relative abundance of 0.1%, resulted in the identification of 39 bacterial species/taxa, representing 97.5% of the total community (**Supplementary Fig. 12a** and **Supplementary Data 2b**). As expected, the gut microbiota was highly dominated by *Bifidobacterium* (average of 64.2 %) and among the top 10 dominating taxa, *B. longum* (38.5 %), *B. breve* (9.1 %), *B. bifidum* (7.9 %), *B. catenulatum* group (6.4 %) and *B. dentium* (1.7 %) were found (**Supplementary Fig. 12a**), with the remaining *Bifidobacterium* spp. being assigned to *B. scardovii* (0.24 %), *B. adolescentis* (0.15 %) and *B. animalis/pseudolongum* (0.10 %) (**Supplementary Data 2b,c**). Although the relative abundance of *Bifidobacterium* increased with time, on average the microbial composition and Shannon diversity did not change dramatically during the six months (**Supplementary Fig. 12b**). However, the subject specific gut microbiota profiles revealed a highly individual species composition (**Supplementary Fig. 12c**) and 48% of the variation in community structure was explained by subject (weighted UniFrac, r^2^ = 0.48, p < 0.001, Adonis, **Supplementary Fig. 13a,b**). Indeed, the difference in microbial community structure between faecal samples were primarily driven by the abundance of *Bifidobacterium* as assessed by PCoA of weighted UniFrac distances (r^2^ = 0.48, p < 0.001, Adonis, **Fig. 3a**). The non-phylogenetic Bray-Curtis dissimilarity analysis revealed a separation of the communities based on abundance of the five dominating *Bifidobacterium* species (*B. longum, B. bifidum, B. breve, B. catenulatum* group and *B. dentium*) (**Fig. 3b** and **Supplementary Fig. 13c-h**). Community abundance of *B. longum, B. bifidum* and *B. breve,* but not *B. catenulatum* group and *B. dentium* (**Fig. 3b**) matched the measured faecal concentrations of aromatic lactic acids (**Fig. 3c**). Consistently, faecal concentrations of ILA, PLA and 4-OH-PLA increased concurrently with an increase in absolute abundance of infant type *Bifidobacterium* species (defined as the summarised abundance of *B. longum, B. bifidum, B. breve* and *B. scardovii*) from birth to around 6 months of age (**Fig. 3d, upper panels**). The gut microbiota of individuals dominated by infant type *Bifidobacterium* spp. was more stable over time than in individuals with a gut microbiota not dominated by infant type *Bifidobacterium* spp. (p < 0.0001, Mann-Whitney *U* test) (**Supplementary Fig. 13i-j**). Using solely samples from breastfed infants, faecal abundances of HMO residuals showed a progressive decline with age, concurrent with the progressive increase in infant type *Bifidobacterium* species (**Fig. 3d, lower panels**). By repeated measure correlation analyses^29^ (**Supplementary Fig. 14**) and partial Spearman’s Rank correlation analyses^30^ adjusted for age, we confirmed that the abundance of infant type *Bifidobacterium* species were positively associated with faecal levels of ILA, PLA and 4-OH-PLA and negatively associated with abundances of HMOs in faeces (**Fig. 3e**). Both subspecies of *B. longum* were associated with the aromatic lactic acids, but mainly *B. longum* subsp. *infantis* was associated with the HMO residuals in faeces (**Fig. 3e**). We have thus established a link between breastfeeding, degradation of HMOs, abundance of specific *Bifidobacterium* spp. and concentrations of aromatic lactic acids in early infancy.

**Fig. 3.**
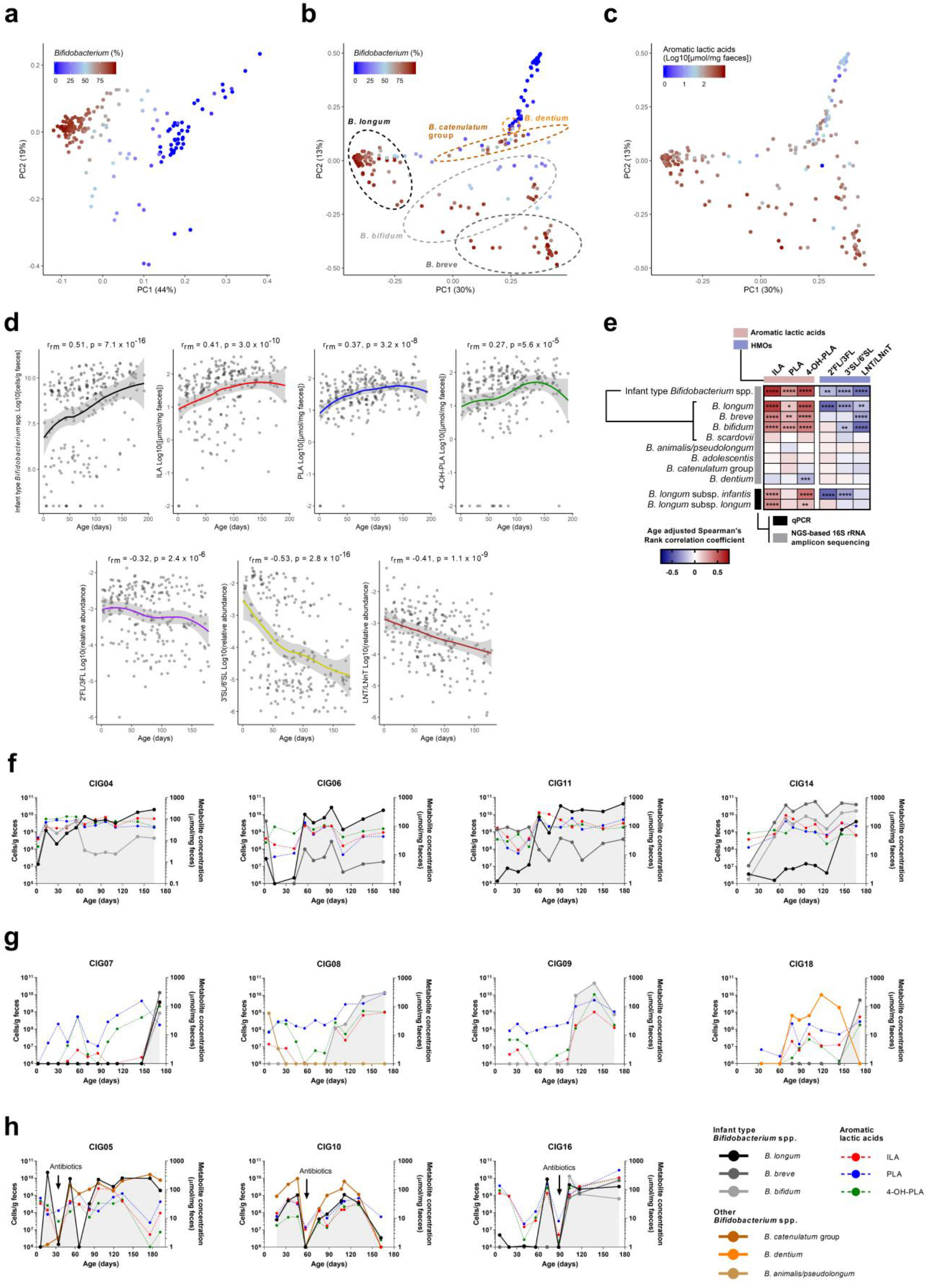
Infant type *Bifidobacterium* species determine aromatic lactic acid concentrations during early infancy. **a-c**, Principal coordinate analysis plots of weighted UniFrac (a) or Bray-Curtis (b-c) distances/dissimilarities (n=234^#^), coloured according to relative abundance of *Bifidobacterium* (a-b) or log10 transformed absolute concentration [μmol/mg faeces] of aromatic lactic acids (sum of indolelactic acid (ILA), phenyllactic acid (PLA) and 4-hydroxyphenyllactic acid (4-OH-PLA)) in the Copenhagen Infant Gut (CIG) cohort. Dashed lined circles indicate communities dominated (relative abundance > 50%) either by *B. longum*, *B. bifidum*, *B. breve*, *B. catenulatum* group or *B. dentium* (*B. adolescentis, B. scardovii* and *B. animalis/pseudolongum* never dominated any of the communities, see **Supplementary Fig 12-13)**. ^#^6 samples were omitted from the analyses due to low read counts (<8000). **d**, Temporal development in absolute abundance of infant type *Bifidobacterium* spp. (defined as the sum of absolute abundances of *B. longum*, *B. breve*, *B. bifidum* and *B. scardovii*), faecal concentrations of aromatic lactic acids (ILA, PLA and 4-OH-PLA, n=240) and relative faecal abundance of HMOs (2’FL/3FL, 2’/3-*O*-fucosyllactose; 3’SL/6’SL, 3’/6’-*O*-sialyllactose; LNT/LNnT, lacto-N-tetraose/ lacto-N-*neo*tetraose, n=228) during the first 6 months of life in the CIG cohort. A local polynomial regression (LOESS) fit is shown with 95% CI shaded in grey. Statistical significance is evaluated by repeated measures correlations (r_rm_). **e**, Heatmap illustrating partial Spearman’s Rank correlation coefficients (adjusted for age) between the absolute abundance of *Bifidobacterium* species/subspecies and absolute faecal concentrations of aromatic lactic acids (n=240) or relative abundances of HMOs (n=228) in the CIG cohort. Infant type *Bifidobacterium* spp. is the sum of absolute abundances of *B. longum*, *B. breve*, *B. bifidum* and *B. scardovii*. Statistical significance is indicated by asterisks with * p<0.05, **p<0.01, ***p<0.001 and ****p<0.0001. **f-h,** Absolute abundance of *Bifidobacterium* spp. (average relative abundance >1% of total community) and concentrations of ILA, PLA and 4-OH-PLA in selected individuals from the CIG cohort. Summarized absolute abundance of infant type *Bifidobactrium* spp. is indicated with grey background shading. **f**, Fully breastfed infants early colonised with infant type *Bifidobacterium* spp. (colonised within first month reaching average relative abundance >40% during first 6 months) and concurrent high absolute concentrations of ILA, PLA and 4-OH-PLA through the first 6 months of life. **g**, Infants with late colonised with infant type *Bifidobacterium* spp. (not detectable or on average <0.5% of total community within the first 3 months of life) and concurrent low concentrations of ILA, PLA and 4-OH-PLA. **h**, Infants with recorded oral antibiotics intake during the first 6 months of life. Similar dynamics of the remaining infants can be seen in **Supplementary Fig. 16**. **See Supplementary Fig 12-16**

Examination of the *Bifidobacterium* and aromatic lactic acid dynamics in each of the 25 infants during the first 6 months of life (**Fig. 3f-h** and **Supplementary Fig. 15-16**) revealed that breastfed infants early and consistently colonised by infant type *Bifidobacterium* species consistently showed high concentrations of aromatic lactic acids in faeces (**Fig. 3f** and **Supplementary Fig. 16a**). In contrast, infants with delayed infant type *Bifidobacterium* species colonisation showed considerably lower concentrations of the aromatic lactic acids, in particular of ILA, despite breastfeeding (**Fig. 3g** and **Supplementary Fig. 16b**). Among the latter, CIG08 and CIG09 were twins, born late preterm, and dominated by an OTU assigned to *Clostridium neonatale* (**Supplementary Fig. 12c** and **Supplementary Data 2b**) in accordance with previous reports on *C. neonatale* overgrowth^31^ and delayed *Bifidobacterium* colonisation^32–35^ in preterm infants. CIG07 who also showed delayed colonisation with infant type *Bifidobacterium*, was mixed fed throughout the whole period and predominantly colonised with *E. coli* and *Clostridium* spp. (**Supplementary Fig. 12c**). CIG18 had relatively low faecal concentrations of aromatic lactic acids until age 172 days, when *B. breve* replaced *B. dentium* (**Fig. 3g**), consistent with the fact that *B. dentium* lacks the *aldh* gene while *B. breve* contains it (**Fig. 2a,b**). Finally, in the three infants treated with antibiotics during our study, *Bifidobacterium* species abundance were temporarily decreased simultaneously with reduced concentrations of the aromatic lactic acids (**Fig. 3h**). This indicates that bifidobacterial aromatic lactic acid production is compromised by pre-term delivery, exposure to antibiotics and formula supplementation. To further corroborate our findings we mined a published metagenomic dataset from faecal samples of a cohort of 98 Swedish mother-infants pairs^4^ for bifidobacterial metagenome-assembled genomes (MAGs) containing the *aldh* gene. This analysis revealed a significantly higher abundance of *aldh* containing MAGs in exclusively breastfed (compared to mixed or formula fed) infants at 4 months and in partially breastfed (compared to weaned) infants at 12 months of age (**Supplementary Fig. 17**). In addition, we found very low abundance of *aldh* containing MAGs in the mothers and a significant decline of these MAGs in infants after introduction to solid foods (4 versus 12 months of age). These results underline that the bifidobacterial potential for production of aromatic lactic acids in the gut is indeed related to breastfeeding.

### Indolelactate modulates immune responses via AhR and HCAR3

The tryptophan-derived metabolite ILA was the most abundant aromatic lactic acid, as well as the most abundant tryptophan catabolite measured in the faeces of infants at 0-6 months (**Supplementary Table 3**) and 9 months of age (**Fig. 1d**). Microbial tryptophan catabolites have been found to contribute to intestinal and systemic homeostasis, in particular by their ability to bind the AhR^12^. Furthermore, aromatic lactic acids have been found to activate HCAR3^36^, which is involved in the regulation of immune function and energy homeostasis^37,38^. In accordance with previous reports^13,14^, we observed modest but significant dose-dependent increases in agonistic activity of ILA in both rat and human AhR reporter gene cell lines (**Supplementary Fig. 18**). Furthermore, all three aromatic amino lactic acids, and especially ILA, showed very potent and dose dependent agonistic activity towards the HCAR3 in a reporter cell line assay (**Supplementary Fig. 19**), in agreement with previous reports^36,38^. To investigate the relationship between gut microbiota, aromatic amino acid metabolites and AhR signalling, the AhR activity induced by sterile-filtered faecal water from selected CIG infants (**Fig. 3f-h**) was associated to the most abundant bacterial taxa and all quantified aromatic amino acid metabolites (n=20) in the same samples (**Fig. 4a**).This revealed that particularly the infant type *Bifidobacterium* spp. were positively associated with AhR activity (**Fig. 4b** & **Supplementary Fig. 20a**). *B. dentium* also correlated positively with AhR activity despite the absence of an *aldh* gene in this species (**Fig. 4a**), which might be due to its association to the AhR agonist IAld (**Fig. 1e**), showing similar AhR reporter activity as ILA (**Supplementary Fig. 17a**). Of all the aromatic amino acid metabolites measured, only faecal concentrations of three known AhR agonists indoleacetic acid (IAA), IAld and in particular ILA were significantly associated with AhR activity (**Fig. 4a, c** & **Supplementary Fig. 20b**). Temporal development in AhR activity was assessed in the CIG samples, and stratification of infants into those colonised early (**Fig. 3f**) and late (**Fig. 3g**) with infant type *Bifidobacterium* spp. revealed a higher AhR activity of the faecal water of early colonised infants (**Fig. 4d-e**).

**Fig. 4.**
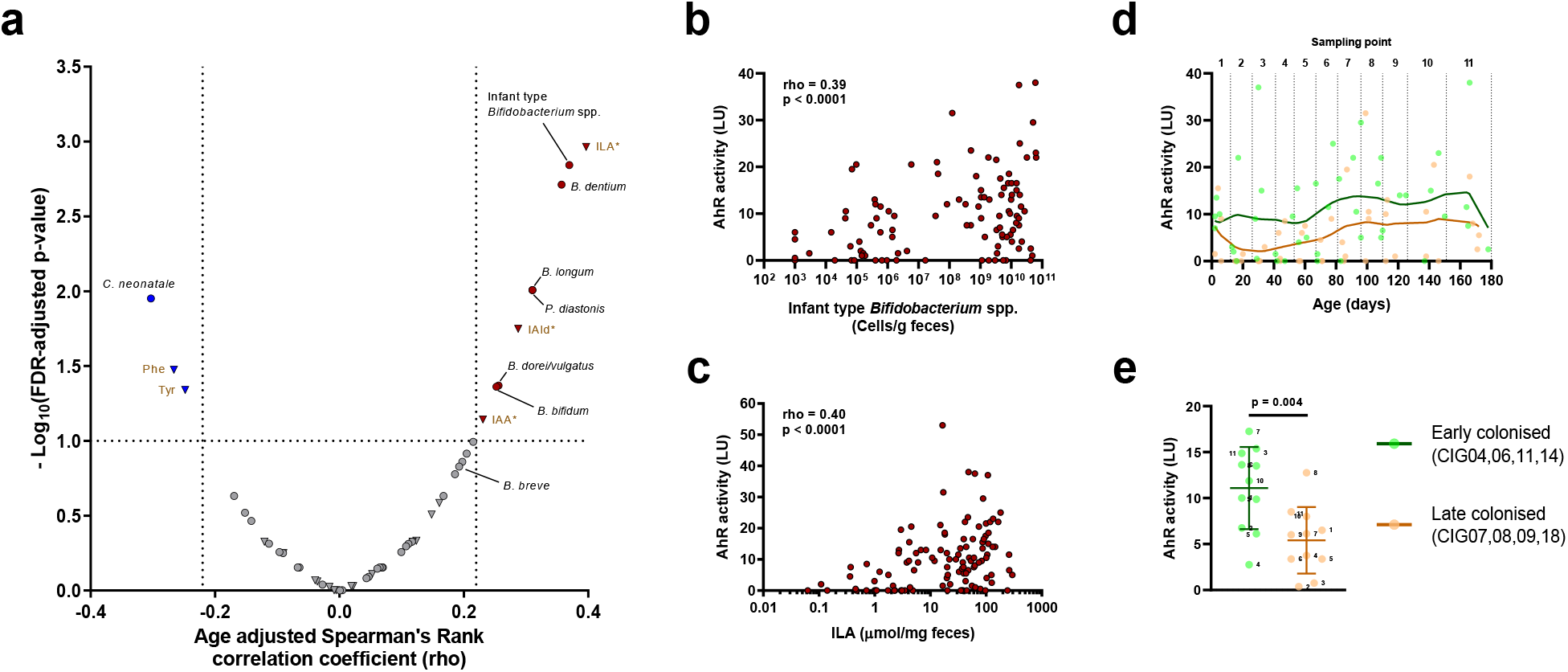
Faecal concentrations of infant type *Bifidobacterium* spp. and indolelactic acid associate with AhR activity. **a,** Scatterplot of age adjusted Spearman’s Rank correlation coefficients (versus associated FDR-adjusted p-values) between AhR activity (Luminescence Units, LU) of faecal water (n=119) from selected Copenhagen Infant Gut (CIG) infants (**Fig. 3f-h,** n=11) in a reporter cell line assay and absolute abundance of gut bacterial taxa (relative abundance>0.1%, n=40, circles) or quantities of aromatic amino acid catabolites (n=20, triangles) measured in the same samples. Coloured circles/triangles mark taxa/metabolites measures that are significantly positively (red) or negatively (blue) associated with AhR activity, within an FDR-adjusted p-value of 0.1. Labels are coloured brown for metabolites and black for bacterial taxa. Asterisks indicate known AhR agonists^13^. **b-c**, Scatterplots showing the associations from (a) between infant type *Bifidobacterium* spp. (b) or ILA (c) and AhR activity in faecal water from selected CIG infants, assessed by age adjusted Spearman’s Rank correlation analysis. **d**, Temporal development (11 samplings points) of *in vitro* AhR activity in CIG infants early (green; CIG04, CIG06, CIG11 and CIG14) or late (orange; CIG07, CIG08, CIG09, CIG18) colonised with infant type *Bifidobacterium* spp. (See **Fig. 3f-g**). Locally weighted regression scatterplot smoothing (LOWESS) curves were fitted to the data points. **e**, Scatterplots (mean ± SD) of AhR activity for each sampling point, stratifying samples into individuals early and late colonised with infant type *Bifidobacterium*. Statistical significance was evaluated by unpaired *t*-test. **See Supplementary Fig 18-19**

Next, we asked whether ILA affects immune function via AhR and HCAR3. Since the human AhR has adapted to sense microbial tryptophan catabolites^39^ and only humans and other hominids contain HCAR3^36^, we isolated immune cells from human blood and assessed the impact of ILA on their function. Specifically, we cultured isolated human CD4^+^ T cells under Th17-polarising conditions and assessed IL-22 production upon exposure to ILA. Interestingly, ILA induced the production of IL-22, an effector cytokine produced by Th17 cells after AhR stimulation^40–42^, in a dose dependent manner (**Fig. 5a**). Conversely, the addition of AhR antagonist CH-223191 inhibited IL-22 production further corroborating that ILA acts through AhR to induce IL-22 production (**Fig. 5b**). We also isolated monocytes from human blood, where both AhR^43^ and HCAR3 are expressed^44^, stimulated the cells with *E. coli* lipopolysaccharide (LPS) and interferon-gamma (IFN-γ) to induce pro-inflammatory conditions, and assessed IL-12p70 production upon ILA exposure. ILA reduced pro-inflammatory IL-12p70 production in a dose-dependent manner (**Fig. 5c**). Addition of CH-223191 blocked the ILA-induced inhibition of IL-12p70 production, confirming that ILA also acts through AhR in human monocytes (**Fig. 5d**). Furthermore, ILA-induced inhibition of IL-12p70 was prevented, when using knockdown of HCAR3 by small interfering RNA (siRNA), supporting that ILA also acts as an anti-inflammatory agent via HCAR3 in human monocytes (**Fig. 5e**). Thus, ILA affects human immune responses via AhR and HCAR3-dependent pathways, suggesting that *Bifidobacterium*-derived ILA is a highly relevant AhR and HCAR3 agonist that may impact immune responses in early life.

**Figure 5.**
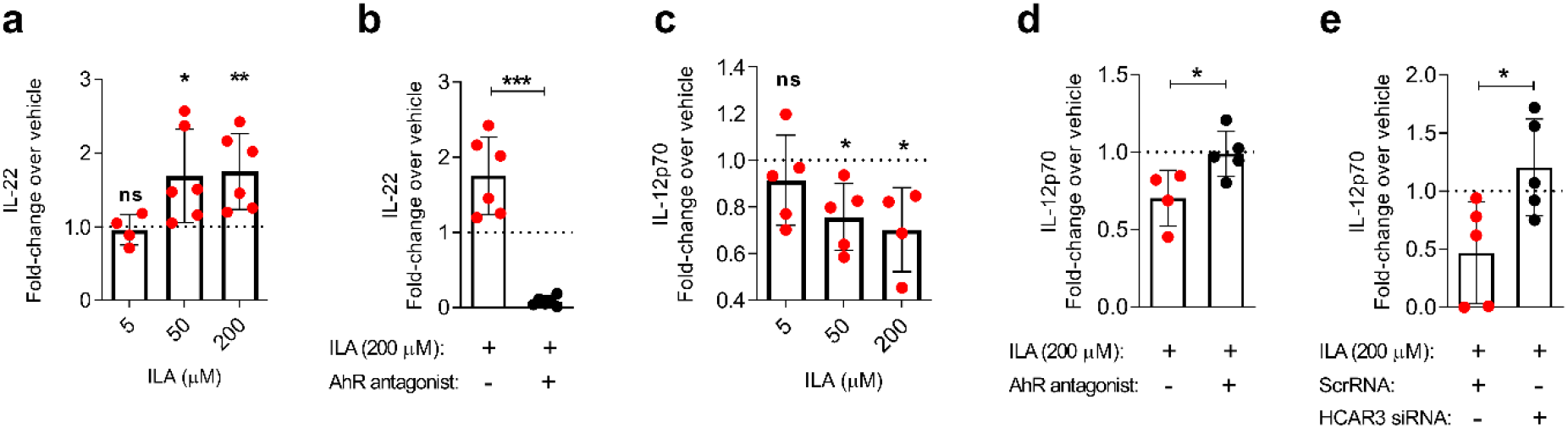
ILA affects human immune responses via AhR and HCAR3. **a,** Fold change in IL-22 production measured by ELISA in purified human CD4+ T cells (4-6 donors, dots represent average per concentration), cultured for 3 days under Th17-polarising conditions in the presence of ILA at 5, 50 and 200 μM compared to vehicle (DMSO control). **b,** Comparison of fold changes over vehicle (DMSO control) in IL-22 production in human purified CD4+ T cells (6 donors) under Th17-polarising conditions in the presence of 200 μM ILA with and without the AhR-antagonist CH-223191. Each dot (·) represents the average of an ELISA test duplicate from an individual donor. Measurements are derived from 2-3 independent experiments. **c,** Fold change in IL-12p70 production in purified human monocytes stimulated with LPS and IFN-γ in the presence of ILA at 5, 50 and 200 μM compared to absence (DMSO control). Dots represent average per concentration. **d,** Comparison of fold changes over vehicle (DMSO control) in IL-12p70 production in purified human monocytes stimulated with LPS and IFN-γ in the presence of 200 μM ILA with and without the AhR-antagonist CH-223191. **e,** Comparison of fold changes over vehicle (DMSO control) in IL-12p70 production in purified human monocytes stimulated with LPS and IFN-γ in the presence of 200 μM ILA and scrambled small interfering RNA (ScrRNA) or HCAR3 small interfering RNA (siRNA). Bars show mean ± SD. Statistical significance was evaluated by paired (panel a-c, e) or unpaired (panel d) *t*-tests and asterisks indicating * p<0.05, **p<0.01, ***p<0.001. For panel a and c, the absolute cytokine values were compared between ILA versus vehicle treated cells. For panel b, d and e the fold changes over vehicle (ratios) were compared between ILA and ILA+ antagonist/siRNA treated cells.

## DISCUSSION

The importance of intestinal commensal bacteria in regulation of the intestinal barrier function and immune development during infancy is well established^45,46^. Yet, specifically the symbiotic role of the breastmilk-promoted *Bifidobacterium* species, which are highly abundant in breastfed infants, remains largely unknown. Here we identified an aromatic lactate dehydrogenase, which catalyses the last step of the conversion of aromatic amino acids into their respective aromatic lactic acids in the infant gut. Although an aromatic lactate dehydrogenase has previously been identified in *Clostridiales* species (*C. sporogenes*, *C. cadaveris* and *P. anaerobius*)^16^, these species are not prevalent nor abundant in the infant gut and have previously been found to convert the aromatic lactic acids into aromatic propionic acids as end products^16^. We show that only the infant type *Bifidobacterium* species, adapted to breastfeeding by their HMO-transport and degradation genes providing them with a colonisation advantage in infant gut^47–51^, contain the aromatic lactate dehydrogenase gene. This fits the observation that *Bifidobacterium* species commonly isolated from the infant gut *in vitro* produce relative higher levels of ILA compared with adult or animal associated *Bifidobacterium* species^52^. Our enzymatic assays showed strong adaptation of ALDH towards indolepyruvate, resulting preferentially in the formation of ILA, the most abundant of the aromatic lactic acids and microbial tryptophan catabolites detected in our two cohorts. Importantly, our data suggest that the production of the AhR agonist ILA by breastmilk-promoted *Bifidobacterium* is a key determinant of AhR-dependent signalling in the gut during infancy. Previous studies have found that ILA *in vitro* decreases inflammation in intestinal cells through activation of AhR^53,54^. Here, we show *ex vivo* that ILA impacts human primary immune cells via AhR and HCAR3-dependent pathways. The observed dose- and AhR-dependent stimulation of IL-22 secretion by ILA may reflect a mechanism by which infant type *Bifidobacterium* species impact intestinal homeostasis in early life, as IL-22 for example provides protection against gastrointestinal pathogens^55–57^, and promotes mucus production^56,58^ and epithelial function^59^. Further, the AhR and HCAR3-dependent inhibitory effect of ILA on IL-12p70 secretion by monocytes may constitute a means by which infant type *Bifidobacterium* species contribute to the regulation of the pro-inflammatory responses to LPS derived from *Enterobacteriaceae* species that often co-inhabit the neonatal/infant gut. While the functional implications of aromatic lactic acids remain to be fully characterised, the phenomenon observed here is likely of fundamental importance, since HCAR3, which is only expressed in humans and other hominids^36^, is involved in the regulation of immune functions and energy homeostasis^37,38^. Furthermore, AhR signalling is involved in protection against gastrointestinal pathogens^13^, and plays a key role in enhancement of intestinal barrier function^60,61^, immune development^17,62–64^, attenuation of induced colitis^65^, autoimmunity^66–68^ and metabolic syndrome^60^. In addition, ILA and PLA have been shown *in vitro* to have direct anti-bacterial^69,70^ and anti-fungal properties^71,72^. Collectively our findings suggest a central role for aromatic lactic acids in mediating host-microbiota interactions in early life.

## METHODS

### Human study populations and metadata

#### SKOT cohort

The discovery cohort consisted of a random subset of 59 healthy infants of the observational SKOT I cohort^20^. These infants were originally recruited from Copenhagen and Frederiksberg by random selection from the National Danish Civil Registry^73^. Inclusion criteria were single birth and full term delivery, absence of chronic illness and age of 9 months ± 2 weeks at inclusion. Mode of delivery, gender, age at sampling, use of medication, breast- and formula feeding prevalence as well as exclusive and total breastfeeding duration and age of introduction to solid foods was recorded by parental questionnaires (**Supplementary Data 1a,b**). Anthropometrics, full dietary assessment and other relevant metadata have been published previously^3,74^. Faecal samples were obtained at 9 months ± 2 weeks of age and were stored at −80°C until DNA extraction, as described previously^3^. Urine samples were collected by the use of cotton balls placed in the infants’ disposable nappies from which the urine was squeezed into a sterile tube and stored at −80°C. In case of faeces in the nappy, the urine sample was discarded. The study protocol was approved by the Committees on Biomedical Research Ethics for the Capital Region of Denmark (H-KF-2007-0003) and The Data Protection Agency (2002-54-0938, 2007-54-026) approved the study. Informed consent was obtained from all parents of infants participating in the SKOT I study.

#### CIG cohort

The validation cohort, CIG, consisted of 25 healthy infants, vaginally born (23/25) and full-term (23/25) delivered. Infants in CIG were recruited through social media and limited to the Copenhagen region. Parents collected faecal samples approximately every second week, starting from the first week of life until 6 months of age (i.e. within week 0, 2, 4, 6, 8, 10, 12, 16, 20 and 24). Parents were instructed to collect faecal samples from nappies into sterile faeces collection tubes (Sarstedt, Nümbrecht, Germany) and immediately store them at −18°C in a home freezer until transportation to the Technical University of Denmark where the samples were stored at −80°C until sample preparation. Gender, pre-term vs full-term birth, mode of delivery, infant/maternal antibiotics, feeding patterns (breastmilk vs formula), introduction to solid foods and consumption of probiotics was recorded (**Supplementary Data 2a**). The Data Protection Agency (18/02459) approved the study. The office of the Committees on Biomedical Research Ethics for the Capital Region of Denmark confirmed that the CIG study was not notifiable according to the Act on Research Ethics Review of Health Research Projects (Journal nr.: 16049041), as the study only concerned the faecal microbial composition and activity and not the health of the children. Informed consent was obtained from all parents of infants participating in the CIG study.

### Gut microbiota analysis

#### 16S rRNA gene amplicon sequencing

Sample preparation and sequencing was performed as previously described^3^ using a subset of 59 faecal samples originating from infants participating in the SKOT I cohort and 241 faecal samples from 25 infants participating in the CIG cohort (data from a total of 28 samples were missing due to insufficient DNA extraction, lack of PCR product, very low number of sequencing reads or resemblance of community to negative controls). Briefly, DNA was extracted from 250 mg faeces (PowerLyzer® PowerSoil® DNA isolation kit, MoBio 12855-100) and the V3 region of the 16S rRNA gene was amplified (30s at 98°C, 24-30 cycles of 15s at 98°C and 30s at 72°C, followed by 5 min at 72°C) using non-degenerate universal barcoded primers^75^ and then sequenced with the Ion OneTouch^TM^ and Ion PGM platform with a 318-Chip v2. Sequences from SKOT and CIG were analysed separately. Briefly, they were de-multiplexed according to barcode and trimmed as previously described^75,76^ in CLC Genomic Workbench (v8.5. CLCbio, Qiagen, Aarhus, DK). Quality filtering (-fastq_filter, MAX_EE_[SKOT]_ = 2.0, MAX_EE_[CIG]_ = 1.0), dereplication, OTU clustering (-cluster_otus, minsize 4), chimera filtering (-uchime_ref, RDP_gold database), mapping of reads to OTUs (-usearch_global, id 97%) and generation of OTU tables (python, uc2otutab.py) was done according to the UPARSE pipeline^77^. In QIIME^78^, OTU tables (n_OTUs[SKOT]_ = 545, n_OTUs[CIG]_ = 478) was filtered to include only OTUs with abundance across all samples above 0.005% of the total OTU counts (n_OTUs[SKOT]_= 258, n_OTUs[CIG]_= 145). OTU relative abundances within samples were estimated by total sum scaling. Taxonomy was assigned to the OTUs using the rdp classifier with confidence threshold 0.5^79^ and the GreenGenes database v13.8^80^. Estimating species composition in the CIG cohort, the OTUs detected with identical taxonomy were collapsed and using a cutoff of average relative abundance of 0.1%, only 39 bacterial species/taxa remained, representing 97.5% of total community (**Supplementary Data 2b & Supplementary Fig. 11**). Based on PyNAST alignment of representative OTU sequences from each cohort separately, a phylogenetic tree was created with FastTree, as described previously^76^. Alpha diversity (Shannon index, Observed OTUs, Pielou’s evenness index) and beta diversity (weighted and unweighted UniFrac distances, abundance weighted and binary Bray-Curtis and abundance weighted Jaccard dissimilarities) measures were calculated in QIIME, with the sequencing depth rarefied to 2,000 (SKOT) - 8,000 (CIG) sequences per sample. Jaccard similarity index (1-abundance weighted Jaccard distance) was computed by calculating the median of all Jaccard similarity index values between adjacent time points within each individual of the CIG cohort. In order to investigate *Bifidobacterium* species composition OTUs sequences classified as *Bifidobacterium* according to the GreenGenes database v13.8 were filtered to remove low abundant OTUs (cutoff 0.1% of total *Bifidobacterium*) and the taxonomy of these resulting OTUs (n_OTUs[SKOT]_ = 23, n_OTUs[CIG]_ = 8) was confirmed by BLAST^81^ search against the 16S rRNA gene sequence database at NCBI. The top BLAST hit indicated species annotation (**Supplementary Data 1f and 2c**). OTUs were collapsed into *Bifidobacterium* species (*B. longum, B. bifidum, B. breve, B. catenulatum* group*, B. adolescentis, B. scardovii, B. dentium, B. animalis/pseudolongum)* based on the top BLAST hit (**Supplementary Data 1f and 2c**). Infant type *Bifidobacterium* spp. were defined as the summarised abundance of *B. longum, B. bifidum, B. breve* and *B. scardovii*. CIG individuals were grouped based on colonisation with infant type *Bifidobacterium* spp., into those with early (colonised within first month reaching average relative abundance >40% during first 6 months) and late colonisation (not detectable or on average <0.5% of total community within the first 3 months of life).

#### Quantitative PCR

Total bacterial load (universal primers) and absolute abundances of *B. longum* subsp. *longum* and *B. longum* subsp. *infantis* (subspecies specific primers) were estimated by quantitative PCR (qPCR), using the primers listed in **Supplementary Table 4**. Each reaction was performed (in triplicates) with 5 μl PCR-grade water, 1.5 μl forward and reverse primer, 10 μl SYBR Green I Master 2X (LightCycler^®^ 480 SYBR Green I Master, Roche) and 2 μl template DNA, in a total volume of 20 μl. Standard curves were generated from 10-fold serial dilutions of linearized plasmid (containing 10^8^ - 10^0^ gene copies/μl), constructed by cloning a PCR amplified 199bp fragment of the 16S rRNA gene (V3-region) of *E. coli* (ATCC 25922) or a 307bp fragment of the Blon0915 gene^82^ of *B. longum* subsp. *infantis* (DSM 20088) or a 301bp fragment of the BL0274 gene^83^ of *B. longum* subsp. *longum* (DSM 20219) into a pCR4-Blunt-TOPO (Invitrogen) or pCRII-Blunt-TOPO vector (Invitrogen). Plates were run on the LightCycler^®^ 480 Instrument II (Roche) with the program including 5 min pre-incubation at 95°C, followed by 45 cycles with 10 sec at 95°C, 15 sec at 50-60°C and 15 sec at 72°C and a subsequent melting curve analysis including 5 min at 95°C, 1 min at 65°C and continuous temperature increase (ramp rate 0.11 °C/s) until 98°C. Data were analysed with the LightCycler^®^ 480 Software (v1.5) (Roche). Bacterial load data (using the universal primers) were used to estimate absolute abundances of each microbial taxa by multiplying with relative abundances (derived from 16S rRNA gene amplicon sequencing).

### *Bifidobacterium* strains and growth experiments

#### Aromatic lactic acid production by Bifidobacterium type strains

*Bifidobacterium* type strains (**Supplementary Table 5**) were cultivated on MRSc (MRS containing 2% (w/v) glucose and supplemented with 0.05% (w/v) L-cysteine) agar plates for 48h at 37°C anaerobically. Single colonies were dissolved in 5.0 mL pre-reduced MRSc broth and incubated for 24h at 37°C anaerobically with shake. The overnight (ON) cultures were washed (10000xg, room temperature, 5 min) and resuspended in sterile 0.9% NaCl water, diluted 1:20 (in triplicates) in pre-reduced MRSc or MRSc+HMOs (MRS broth without glucose, but supplemented with 2.0% (w/v) HMO mixture and 0.05% (w/v) L-cysteine) and re-incubated at 37°C anaerobically for 72h, after which OD_600nm_ was measured and the culture supernatants (16000xg, 5 min, 4°C) were analysed by UPLC-MS. The individual HMOs were kindly donated by Glycom A/S (Hørsholm, Denmark); 2’-*O*-fucosyllactose (2’FL), 3-*O*-fucosyllactose (3FL), lacto-*N*-tetraose (LNT), lacto-*N*-*neo*tetraose (LN*n*T), 6’-*O*-sialyllactose (6’SL), 3’-*O*-sialyllactose (3’SL), together representing the three structures found in human breastmilk (fucosylated, sialylated, and neutral core). Based on the HMO composition in breastmilk^84,85^, these were mixed in a ratio of 53% 2’FL, 18% 3FL, 13% LNT, 5% LN*n*T, 7% 6’SL and 4% 3’SL in sterile water to obtain a representative HMO mix used in the *in vitro* experiments at 2% (w/v).

#### Growth experiment with B. longum subsp. longum 105-A strains

*B. longum* subsp. *longum* 105-A (JCM 31944) was obtained from Japan Collection of Microorganisms (RIKEN BioResource Research Center, Tsukuba, Japan). *B. longum* subsp. *longum* 105-A strains (wild type [WT], insertional mutant [type4 *ldh*::pMSK127] and complemented insertional mutant [type4 *ldh*::pMSK127 / pMSK128 (P*xfp*-type4 *ldh*)]; **Supplementary Table 5**) were cultivated on MRSc or MRSc-Chl (MRSc supplemented with 2.5 μg/mL chloramphenicol) agar plates for 48h at 37°C anaerobically. Single colonies were dissolved in 5.4 mL MRSc or MRSc-Chl broth, 10-fold serially diluted and incubated for 15h at 37°C anaerobically with shake. The most diluted culture (exponential phase) was washed in same medium (10000xg, room temperature, 5 min) and resuspended in MRSc or MRSc-Chl broth to yield OD_600nm_ = 1 and subsequently diluted 1:40 in prewarmed and reduced MRSc or MRSc-Chl broth (in triplicates), before incubation at 37°C, anaerobically with shake. The cultures were sampled (500 μL) every hour for OD_600nm_ measurements and the culture supernatants (16000xg, 4°C, 5 min) from early (13h) stationary phase was analysed by UPLC-MS for aromatic amino metabolites and by GC-MS for lactate.

### Alignments and construction of phylogenetic trees

From the full genome sequences (available at https://www.ncbi.nlm.nih.gov/genome/) of *Bifidobacterium* type strains included in this study (**Supplementary Table 5**) all genes annotated as L-lactate dehydrogenases were aligned (gap cost 10, gap extension cost 1) and subsequently a phylogenetic tree (Algorithm = Neighbor-Joining, Distance measure = Jukes-Cantor, 100 bootstrap replications) was constructed in CLC Main Workbench (v7.6.3, CLCbio, Qiagen, Aarhus, DK). The tree was visualised by use of the FigTree software v1.4.3 (http://tree.bio.ed.ac.uk/software/figtree/). For identification of *aldh* (type 4 *ldh*) in *Bifidobacterium* strains, all complete human gut associated *Bifidobacterium* genomes (n = 127) including plasmids were retrieved from NCBI Genome (14 February 2020) and *aldh* genes were identified using NCBI BLAST+ v. 2.10.0+ tBLASTn with default settings and a cutoff of 70% identity and 70% query coverage. Aligned genomic nucleotide sequences were translated and verified to match lactate dehydrogenases using reciprocal BLASTx against NCBI’s nr database. In addition, the ALDH amino acid sequence (translated from the *aldh* nucleotide sequence) of *B. longum* subsp. *longum* 105-A was aligned (gap cost 10, gap extension cost 1) with the ALDH amino acid sequences of the *B. longum* subsp. *longum*, *B. longum* subsp. *infantis*, *B. bifidum*, *B. breve* and *B. scardovii* type strains. Further, comparison of *aldh* gene cluster/operon in 12 *Bifidobacterium* type strains (**Supplementary Fig. 5**) was basically conducted by pairwise alignments in MBGD (Microbial Genome Database for Comparative Analysis; http://mbgd.genome.ad.jp/). The amino acid sequences of the gene cluster from *B. pseudolongum* subsp. *pseudolongum* type strain was collected from NCBI database (https://www.ncbi.nlm.nih.gov/genome/) and was used for comparison with that from *B. animalis* subsp. *animalis* type strain.

### Identification of *aldh* in metagenomic data

Using 193 infant samples collected at 4 and 12 months of age with data on feeding practice available and data from 98 mothers^4^, we used IGGsearch and IGGdb v.1.0.0^86^ to identify *Bifidobacterium* metagenome assembled genomes (MAGs). MAGs were included in the analysis if they passed the following criteria: --min-reads-gene=2 --min-perc-genes=40 --min-sp-quality=75. For each *Bifidobacterium* MAG identified, we used the representative genome to search for *aldh* genes. *aldh* genes were identified using NCBI BLAST+ v. 2.10.0+ tBLASTn with default settings and a cutoff of 70% identity and 70% query coverage.

### Recombinant expression of *aldh* (type 4*ldh*)

#### Chemically competent cells for recombinant expression

*E. coli* LMG 194 ON culture (200μL) was inoculated into 5 ml Luria-Bertani (LB) medium and incubated at 37°C, 250 rpm until OD_600nm_ = 0.5, at which the culture was centrifuged 5 min at 10000xg at 4°C and supernatant discarded. Cell pellet was resuspended in ice-cold 1.8 ml 10mM MgSO_4_ (Sigma, M2643) and centrifuged for 2 min at 5000xg at 0°C. Supernatant was discarded and cell pellet resuspended in 1.8 ml ice-cold 50 mM CaCL_2_ (Merck, 1.02083.0250), incubated on ice for 20 min and centrifuged for 2 min at 5000xg at 0°C. Cell pellet was resuspended in 0.2 ml ice-cold 100mM CaCl_2_, 10mM MgSO_4_ and placed on ice until transformation.

#### Cloning and recombinant expression

Genomic DNA was extracted (PowerLyzer® PowerSoil® DNA isolation kit, MoBio 12855-100) from colony material of *B. longum* subsp. *infantis* DSM 20088^T^. In order to amplify the type 4 *ldh* gene, 50 ng template DNA was mixed with 5 μL 10X PCR buffer, 0.5 μL (50 mM) dNTP mix, 1 μL (10μM) forward primer (ldh4_F, 5’-ACCATGGTCACTATGAACCG-3’), 1 μL (10μM) reverse primer (ldh4_R, 5’-AATCACAGCAGCCCCTTG-3’) and 1 μL (1 U/μL) Platinum Taq DNA polymerase (Invitrogen, 10966-018) in a 50 μL total reaction volume. The PCR program included 2 min at 94°C, 35 cycles of 30 sec at 94°C, 30 sec at 55°C, 60 sec at 72°C, followed by a final extension 10 min at 72°C. The PCR product was purified (MinElute PCR purification kit, Qiagen, 28004) and 4 μL was mixed with 1 μL Salt solution (1.2M NaCl, 0.06M MgCl_2_) and 1 μL pBAD-TOPO® plasmid (Invitrogen, K4300-01) and incubated for 5 minutes at room temperature. 2 μL of the cloning mixture was transformed into 50 μL One Shot® TOP10 Competent Cells (Invitrogen, K4300-01) by gentle mix, incubation 15 min on ice and heat-shock for 30 sec at 42°C. 250μL S.O.C medium (Invitrogen, K4300-01) was added and incubated at 37°C for 1 h at 200 rpm and subsequently spread on LB-AMP (LB supplemented with 20 μg/ml Ampicillin (Sigma, A9518)) agar plates and incubated at 37°C ON. Transformants were picked and clean streaked on LB-AMP agar plates, incubated 37°C, ON and afterwards single colonies of each transformant was inoculated into 5 ml LB-AMP broth and incubated at 37°C for 15 h at 250 rpm. Plasmid DNA was isolated (QIAprep Spin Miniprep Kit, Qiagen, 27104) from each transformant and subsequently 5 μL plasmid DNA (80-100 ng/μL) was mixed with 5μL (5pmol/μL) pBAD forward (5’-ATGCCATAGCATTTTTATCC-3’) or reverse (5’-GATTTAATCTGTATCAGG-3’) sequencing primers (5pmol/μL) and shipped for sequencing at GATC (GATC-biotech, Koln, Germany). In order to remove the leader peptide in pBAD-TOPO, 10 μL plasmid (0.1 μg) with correct insert was cut with FastDigest *NcoI* (Thermo Scientific, FD0563) for 10 min at 37°C and the enzyme inactivated 15 min at 65°C. Plasmid was ligated using 1 μL (1U/μL) T4 DNA Ligase (Invitrogen, 15224-017) for 5 min at room temperature and subsequently 2μl plasmid was transformed into 100μl chemically competent *E. coli* LMG194 cells by incubation on ice for 30 min, followed by heat-shock at 43°C for 3 min and incubation on ice for 2 min. 900μL LB medium was added and cells were incubated at 37°C for 1h at 250 rpm, before plating on LB-AMP agar plates and incubation at 37°C ON. Transformants were picked, clean streaked and plasmid DNA isolated and sequenced as described above. A transformant with correct insert was selected for recombinant expression of the type 4 *ldh* gene; 2 ml LB-AMP broth was inoculated with a single recombinant colony or the non-transformed *E. coli* LMG194 (negative control) and grown at 37°C ON at 250rpm. In 3x triplicates, 100 μL of the ON cultures (2×3x 100 μL transformant culture + 1×3x 100 μL non-transformed *E. coli* LMG194 culture) were diluted 100-fold into 9.9 mL prewarmed LB-AMP/LB broth and grown at 37°C, 250 rpm until OD_600nm_ ≈ 0.5, at which 9 mL culture was added 1 mL mix of indolepyruvic acid, phenylpyruvic acid and 4-hydroxyphenylpyruvic acid (1mg/mL each). The cultures were sampled (time zero) and subsequently 100 μL 20% L-arabinose (or 100 uL sterile water; control for induction) was added to induce gene expression and the cultures were re-incubated at 37°C, 250 rpm, before sampling at 1 h and 5 h post-induction for OD_600nm_ measurements and assessment of production of aromatic lactic acids. For the latter, samples were centrifuged at 16000xg, 5 min, 4°C and supernatants were stored at −20°C for UPLC-MS analyses.

### Construction of *aldh* (type 4 *ldh*) insertional mutant

#### Transformation of B. longum subsp. longum 105-A

*B. longum* subsp. *longum* 105-A cells were grown to exponential phase at 37°C in Gifu anaerobic liquid medium (Nissui Pharmaceutical Co., Ltd., Tokyo, catalog no. 05422), harvested by centrifugation, and washed twice with ice-cold 1 mM ammonium citrate buffer containing 50 mM sucrose (pH = 6.0). The cells were concentrated 200 times with the same buffer and used for electroporation with settings of 10 kV/cm, 25 μF, and 200Ω. After recovery culturing in Gifu anaerobic liquid medium at 37°C for 3 h, the cells were spread onto Gifu anaerobic agar containing antibiotics (i.e. 30 μg/mL spectinomycin and/or 2.5μg/mL chloramphenicol) for selection.

#### Insertional mutant construction and plasmid complementation

The type 4 *ldh* gene (BL105A_0985) of *B. longum* subsp. *longum* 105-A was disrupted by a plasmid-mediated single crossover event as described previously^87^. The plasmid used for disruption was constructed using the In-Fusion cloning kit (Clontech Laboratories, Inc., Mountain View, CA, USA, catalog no. 639649). *Escherichia coli* DH5α was used as a host. In brief, the internal region of the *ldh* gene (position 142-638 of the nucleotide sequence of BL105A_0985, see **Supplementary Fig. 6**) was amplified by PCR using a primer pair Pr-580/581 (**Supplementary Table 6**) and ligated with the BamHI-digested pBS423 fragment carrying pUC *ori* and a spectinomycin resistance gene^27^. The resulting plasmid pMSK127 was introduced into *B. longum* subsp. *longum* 105-A by electroporation to be integrated into type 4 *ldh* locus by single crossover recombination (type 4 *ldh*::pMSK127). Type 4 *ldh* disruption was confirmed by PCR with a primer pair designed to anneal outside of the gene (**Supplementary Fig. 6** and **Supplementary Table 6**). The amplified fragment was also sequenced to ensure the correct recombination event.

Complementation plasmid pMSK128 was constructed by ligating PCR-amplified *xfp* (xylulose 5-phosphate/fructose 6-phosphate phosphoketolase) promoter region (P*xfp*) and the type 4 *ldh* coding region with PstI- and SalI-digested pBFS38^88^ using the In-Fusion cloning kit, by which type 4 *ldh* was placed under the control of P*xfp*. Primer pairs of Pr-598/Pr-599 and Pr-600/Pr-601 were used for amplifying P*xfp* from pBFS48^88^ and the type 4 *ldh* gene from the *B. longum* subsp. *longum* 105-A genome, respectively (**Supplementary Table 6**). The resulting plasmid was electroporated into type 4 *ldh*::pMSK127 to give type 4 *ldh*::pMSK127 / pMSK128 (P*xfp*-type4_*ldh*) (**Supplementary Fig. 6**).

### Biochemical characterisation of ALDH (type 4 LDH)

#### Recombinant expression and purification

Type 4 LDH (BL105A_0985) was recombinantly expressed as a non-tagged form. The gene was amplified by PCR using the genomic DNA of *B. longum* subsp. *longum* 105-A as a template and a primer pair of Pr-617 (5’-GGTGGTGGTGCTCGAGTCACAGCAGCCCCTCGCAG-3’) and Pr-635 (5’-AAGGAGATATACATATGGTCACTATGAACCGC-3’). Underlined bases indicate 15-bp for In-Fusion cloning (Clontech). The amplified DNA fragment was inserted into the NdeI and XhoI site of pET23b(+) (Novagen) using an In-Fusion HD cloning kit (Clontech). The resulting plasmid was introduced into *E. coli* BL21 (DE3) Δ*lacZ* carrying pRARE2^87^, and the transformant was cultured in LB medium supplemented with ampicillin (100 μg ml^−1^) and chloramphenicol (7.5 μg ml^−1^). When OD_600nm_ reached 0.5, isopropyl β-D-thiogalactopyranoside was added at a final concentration of 0.02 mM to induce the protein expression. The culture was incubated for four days at 18°C, harvested by centrifugation, and resuspended in 50 mM potassium phosphate buffer (KPB; pH 7.0) supplemented with 1 mM 2-mercaptoethanol (2-ME) and 200 μM phenylmethane sulfonyl fluoride. Following cell disruption by sonication, the cleared lysate was saturated with ammonium sulfate (40–60%). The resulting precipitate was dissolved, dialysed against 20 mM KPB (pH 7.0) containing 1 mM 2-ME, and concentrated by Amicon Ultra 10K centrifugal device (Merck Millipore). The sample was then loaded onto an Affigel blue column (Bio-Rad) preequilibrated with 20 mM KPB (pH 7.0) containing 1 mM 2-ME, and eluted by the same buffer containing 1 M NaCl. The protein was further purified by a Mono Q 5/50 (GE Healthcare; a linear gradient of 0–1 M NaCl in 20 mM Tris-HCl (pH 8.0) containing 1 mM 2-ME) and Superdex 200 Increase 10/300 GL column (GE Healthcare; 10 mM KPB [pH 7.0] containing 50 mM NaCl and 1 mM 2-ME). Protein concentration was determined by measuring the absorbance at 280 nm based on a theoretical extinction coefficient of 26,470 M^−1^ cm^−1^.

#### Enzyme assay

The standard reaction mixture contained 100 mM KPB (pH 8.0), 1 mM 2-ME, 0.1 mM β-NADH, and the substrate. The reaction was initiated by adding the enzyme, and the mixture was incubated at 37°C for an appropriate time, in which the linearity of the reaction rate was observed. The substrate concentrations were varied between 0.01 and 0.25 mM for IPA, 1.5 and 12.75 mM for PPA, 2 and 24 mM for 4-OH-PPA, and 2.5 and 40 mM for pyruvic acid. The enzyme was used at the concentrations of 0.22 nM for IPA, 1.47 nM for PPA, 0.12 nM for 4-OH-PPA, and 88.50 nM for pyruvic acid. The reducing reactions of PPA and pyruvic acid was continuously monitored by measuring the decrease of the absorbance at 340 nm (NADH consumption). When 4-OH-PPA and IPA were used as the substrates, the reaction products 4-OH-PLA and ILA were quantified by HPLC after the termination of the reactions by adding 5 % (w/v) trichloroacetic acid. HPLC analysis was performed using a Waters e2695 separation module (Waters) equipped with a LiChrospher 100 RP-18 column (250 × 4 mm, φ = 5 μm; Merck Millipore) at 50°C. Following equilibration with a mixture of 10% solvent A (50% methanol, 0.05% trifluoroacetic acid) and 90% solvent B (0.05% trifluoroacetic acid) at a flow rate of 1 mL/min, the concentration of solvent A was linearly increased to 100 % for 25 min and maintained at 100 % for additional 15 min. 4-OH-PLA and ILA were detected by a Waters 2475 Fluorescence Detector with λ_ex_ 277 nm and λ_em_ 301 nm and λ_ex_ 282 nm and λ_em_ 349 nm, respectively. The standard curves were created using the known concentrations of both compounds. The kinetic parameters (*k*_cat_, *K*_0.5_, and Hill coefficient *n*H) were calculated by curve-fitting the experimental data to the Hill equation, using KaleidaGraph version 4.1 (Synergy Software). Experiments were performed at least in duplicate. Physicochemical property of the enzyme was examined by using 1 mM PPA as a substrate. The effects of metal ions (0.1 mM each) on the enzyme activity was examined using 50 mM MES (2-(*N*-morpholino)ethanesulfonic acid) buffer (pH 7.0). EDTA (ethylenediaminetetraacetic acid) was added at the final concentration of 0.1, 0.5, or 1 mM. The optimal pH was determined using 50 mM KPB (pH 6.0–8.5) and TAPS (*N*-Tris(hydroxymethyl)methyl-3-aminopropanesulfonic acid) buffer (pH 8.0–9.0). The thermostability was evaluated by the residual activities after incubating the enzyme (1.0 mg/ml in 10 mM KPB [pH 7.0] containing 50 mM NaCl and 1 mM 2-ME) at the indicated temperatures for 30 min prior to the assay. Fructose-1,6-bisphosphate, shikimate-3-phosphate, D-erythrose-4-phosphate, and phosphoenolpyruvate were added to the reaction mixtures at the concentrations of 0.1 and 1 mM to examine their heterotropic effects. KPB, TAPS buffer, or HEPES (4-(2-hydroxyethyl)-1-piperazineethanesulfonic acid) buffer (pH 8.0 each) containing 1 and 4 mM PPA as a substrate was used. The effect of phosphate ion was analysed by adding various concentration of KPB (pH 8.0) into 10 mM HEPES buffer (pH 8.0). All experiments were conducted at least in duplicate. In the subsequent kinetic analysis, we used phosphate ion at the concentration of 100 mM because (i) no saturation was obtained for phosphate under the tested conditions (**Supplementary Fig. 10a**), (ii) the intracellular phosphate concentration in Gram-positive bacteria is known to be 130 mM at maximum^89^, and (iii) the strong homotrophic effect of the substrate PPA was observed only in the presence of 10 mM phosphate ion.

### *In vivo* mono-colonisation experiments

Germ-free (GF) Swiss Webster mice (Tac:SW, originally obtained from Taconic Biosciences, NY, USA) were bred and housed within GF isolators (Scanbur, Karlslunde, Denmark) in type II makrolon cages (Techniplast, Varese, Italy) with bedding, nesting material, hiding place and a wooden block at the National Food Institute, Technical University of Denmark. The mice were fed an irradiated standard Altromin 1314 chow (Brogaarden, Gentofte, Denmark) and the environment was maintained on a 12h light/12h dark cycle at a constant temperature of 22 ± 1 °C, with air humidity of 55 ± 5% relative humidity and change of air 50 times per hour. The GF condition of the mice prior to inoculation of bacteria was confirmed by plating of faecal sample suspensions on blood agar plates (Statens Serum Institut, Copenhagen, Denmark) incubated both aerobically and anaerobically. In two separate experiments, GF mice were colonised with either *B. longum* 105-A wild-type or *aldh* (type 4 *ldh*) mutant by a single oral gavage (200 μl, ≈5×10^7^ CFU/dosis). The mono-colonised mice were euthanised by cervical dislocation and dissected in order to collect cecal contents. Successful colonisation with *B. longum* and absence of contamination in GF offspring was confirmed by cultivation of cecal content on MRSc and blood agar plates incubated both aerobically and anaerobically. Aromatic lactic acids were quantified from cecal content. All mouse experiments were approved by the Danish Animal Experiments Inspectorate (License number 2015-15-0201-00553) and carried out in accordance with existing Danish guidelines for experimental animal welfare.

## Metabolomics

### Chemicals

Authentic standards of the aromatic amino acids and derivatives (**Supplementary Table 1**) were obtained from Sigma Aldrich (Germany), whereas isotope-labelled aromatic amino acids used as internal standards (L-Phenylalanine (ring-d5, 98%), L-Tyrosine (ring-d4, 98%), L-Tryptophan (indole-d5, 98%) and indoleacetic acid (2,2-d2, 96%)) of the highest purity grade available were obtained from Cambridge Isotope Laboratories Inc. (Andover, MA).

### Extraction of metabolites from faecal samples

Faecal samples (100-500 mg) were diluted 1:2 with sterile MQ water, vortexed for 10 seconds and centrifuged at 16.000xg, 4°C for 5 minutes. Subsequently, the supernatant liquor was transferred to a new tube and centrifuged again at 16.000xg, 4°C for 10 minutes. Finally, an aliquot of 150-300 μL was stored at −20°C. All samples were later thawed at 4°C, centrifuged at 16.000xg, 4°C for 5 minutes, and diluted in a total volume of 80 μL water corresponding to a 1:5 dilution of the faecal sample. To each sample, 20 μL internal standard mix (4 μg/mL) and 240 μL of acetonitrile were added. The tubes were vortexed for 10 seconds and left at −20°C for 10 minutes to precipitate the proteins. The tubes were then centrifuged at 16.000xg, 4°C for 10 minutes and each supernatant (320 μL) was transferred to a new tube, which was dried with nitrogen gas. Subsequently, the residues were reconstituted in 80 μL water (equalling a 1:5 dilution of the faecal sample with internal standards having a concentration of 1 μg/mL), vortexed for 10 seconds, centrifuged at 16.000xg, 4°C for 5 minutes, and transferred to LC vial, which was stored at −20°C until analysis.

### Extraction of metabolites from urine samples

Urine samples (n=49) from the SKOT cohort were thawed in a refrigerator and all procedures during the sample preparation were carried out at 0-4°C using an ice bath. The subjects were randomised between analytical batches by placing all the samples from the each subject in the same 96 well-plate. The run order of the samples was randomised within the analytical batch. Urine samples were centrifuged at 3000xg for 2 minutes at 4°C. 150 μL of each urine sample were added to separate wells and diluted with 150 μL of diluent (MQ water: Formic acid (99.9:0.1, v/v) / Internal standard mixture (100 μg/mL) (90:10, v/v). A blank sample (diluent), standard mixture of external standard containing 44 biologically relevant metabolites (metabolomics standard)^90^ and pooled sample containing equal amount of each sample (20 μL) were added to spare wells as quality control samples. The plates were stored at −80°C until the analysis. Immediately prior to analysis, the plates were thawed and mixed by vortex stirring for 10 minutes.

### Extraction of metabolites from in vitro fermentation samples

Supernatants from *in vitro* fermentations were thawed at 4°C, centrifuged at 16.000xg, 4°C for 10 minutes, before 80 μL was transferred to a new tube. To each sample, 20 μL internal standard (40 μg/mL) and 300 μL of acetonitrile were added. The tubes were vortexed for 10 seconds and left at −20°C for 10 minutes to precipitate the proteins. Following, the tubes were centrifuged at 16.000xg, 4°C for 10 minutes before 50 μL of each sample was diluted with 50 μL of sterile water and transferred to a LC vial (equalling a 1:10 dilution of the sample with internal standards having a concentration of 1 μg/mL).

### Metabolic profiling of faecal and in vitro samples using UPLC-MS

Aromatic amino acids and derivatives (**Supplementary Table 1**) of faecal and *in vitro* samples were quantified by ultra performance liquid chromatography mass spectrometry (UPLC-MS) as previously published^91^.

In brief, samples were analysed in random order. For the analysis of the CIG faecal samples, a pooled quality control (QC) sample was injected for every 10 sample. In all cases, five standard mix solutions (0.1 μg/mL, 0.5 μg/mL, 1 μg/mL, 2 μg/mL and 4 μg/mL) were analysed once for every 10 samples to obtain a standard curve for every 10 samples. For each sample, a volume of 2 μL was injected into a ultra-performance liquid chromatography quadrupole time-of-flight mass spectrometry (UPLC-QTOF-MS) system consisting of Dionex Ultimate 3000 RS liquid chromatograph (Thermo Scientific, CA, USA) coupled to a Bruker maXis time of flight mass spectrometer equipped with an electrospray interphase (Bruker Daltonics, Bremen, Germany) operating in positive mode. The analytes were separated on a Poroshell 120 SB-C18 column with a dimension of 2.1×100 mm and 2.7 μm particle size (Agilent Technologies, CA, USA) as previously published^91^. Aromatic amino acids and derivatives were detected by selected ions and quantified by isotopic internal standards with similar molecular structures as listed in **Supplementary Table 1**. Data were processed using QuantAnalysis version 2.2 (Bruker Daltonics, Bremen, Germany) and bracket calibration curves for every 10 lumen samples were obtained for each metabolite. The calibration curves were established by plotting the peak area ratios of all of the analytes with respect to the internal standard against the concentrations of the calibration standards. The calibration curves were fitted to a quadratic regression.

For untargeted metabolomics, the raw UPLC-MS data, obtained by analysis of the CIG faecal samples in positive ionisation mode, were converted to mzXML files using Bruker Compass DataAnalysis 4.2 software (Bruker Daltonics) and pre-processed as previously reported^92^ using the R packpage XCMS (v.1.38.0)^93^. Noise filtering settings included that features should be detected in minimum 50% of the samples. A data table was generated comprising mass-to charge (*m/z*), retention time and intensity (peak area) for each feature in the every sample. The data were normalised to the total intensity. Subsequently, features with a coefficient of variation above 0.3 in the QC samples and features with a retention time below 0.5 min were excluded from the data. Parent ion masses of compounds of interest (2’FL/3FL, LNT/LN*n*T, 3’SL/6’SL) were searched in the cleaned dataset with 0.02 Da *m/z* and 0.02 min retention time tolerance. Subsequently, the identities of the features of interest were confirmed at level 1^94^ by tandem mass spectrometry and comparison to authentic standards (**Supplementary Table 7**). Of notice, HMO isomers could not be distinguished with the method applied due to identical retention times.

### Metabolic profiling of urine samples using UPLC-MS

The samples were analysed by UPLC coupled with a quadrupole-Time of Flight Mass Spectrometer (q-TOF-MS) equipped with an electrospray ionisation (ESI) (Waters Corporation, Manchester, UK). Reverse phase HSS T3 C_18_ column (2.1×100 mm, 1.8 μm) coupled with a pre-column (VanGuard HSS T3 C18 column (2.1×5 mm, 1.8 μm)) were used for chromatographic separation. Five μl of each well was injected into the mobile phase A (0.1 % formic acid in MQ water), mobile phase B (10% 1M ammonium acetate in methanol), mobile phase C (methanol) and mobile phase D (isopropanol). Mobile phase gradient during the run time of 10 min was as follows: start condition (100 % A), 0.75 min (100 % A), 6 min (100 % C), 6.5 min (70 % B, 30 % D), 8 min (70 % B, 30 % D), 8.1 min (70 % B, 30 % D), 9 min (100 % A), 10 min (100 % A). The flow rate gradient was as follows: start condition (0.4 mL/min), 0.75 min (0.4 mL/min), 6 min (0.5 mL/min), 6.5 min (0.5 mL/min), 8 min (0.6 mL/min), 8.10 min (0.4 mL/min), 9 min (0.4 mL/min), 10 min (0.4 mL/min). ESI was operated in negative mode with 3.0 kV capillary probe voltage. The cone voltage and the collision energy were set at 30 kV and 5 eV, respectively. Ion source and desolvation gas (nitrogen) temperature were 120 and 400 °C while sampling cone and desolvation gas flow rates were 50 and 1000 l/hr. Scan time set as 0.08 s with 0.02 sec interscan time for both modes. Data were acquired in centroid mode with mass range between 50 to 1500 Da. Leucine-enkephalin (500 ng/ml) was infused as the lock-spray agent to calibrate the mass accuracy every 5 sec with 1 sec scan time. Quality control samples were used to evaluate possible contamination, monitoring the changes in mass accuracy, retention time and instrumental sensitivity drifts^90,95^.

The raw data were converted to netCDF format using DataBridge Software (Waters, Manchester, UK) and imported into MZmine version 2.28^96^. A subset of samples was used to optimise the pre-processing parameters for the positive and negative mode data separately. Optimised pre-processing parameters are listed in **Supplementary Table 8**. Data pre-processing was employed with the following steps: mass detection, chromatogram builder, chromatogram deconvolution, deisotoping, peak alignment and gap filling. After the pre-processing, each detected peak was represented by a feature defined with a retention time, *m/z* and peak area.

The data matrix was imported into MATLAB R2015b (The MathWorksInc., Natick, MA). Features that were present in the blanks, were very early and late eluting (rt<0.30 and rt>9.46 min), potential isotopes, duplicates as well as features with masses indicating multiple charges were removed from the dataset using an in-house algorithm. The data were normalised using unit length normalisation to correct the variation in urine concentration. Parent ion masses of the aromatic lactic acids (ILA, PLA and 4-OH-PLA) were searched in the cleaned dataset with 0.02 Da *m/z* and 0.02 s retention time tolerance. A linear regression model were employed feature wise to correct for batch differences and instrumental sensitivity drifts^97^. The aromatic lactic acids were confirmed at level 1 ^94^ by comparison to authentic standards and by tandem mass spectrometry using the same experimental conditions (**Supplementary Fig. 2-4**).

### Lactate production by B. longum subsp. longum 105-A strains using GC-MS

The lactate production of the *B. longum* subsp. *longum* 105-A wild-type, type 4 *ldh* mutant and type 4 *ldh* complemented strains were assessed in supernatants obtained after 13h of growth (early stationary phase) by gas-chromatography mass spectrometry (GC-MS) upon methyl chloroformate (MCF) derivatisation using a slightly modified version of the protocol previously described^98^. All samples were analysed in a randomised order. Analysis was performed using GC (7890B, Agilent Technologies, Inc., Santa Clara, CA) coupled with a quadrupole detector (59977B, Agilent Technologies, Inc., Santa Clara, CA). The system was controlled by ChemStation (Agilent Technologies, Inc., Santa Clara, CA). Raw data was converted to netCDF format using Chemstation, before the data was imported and processed in Matlab R2014b (Mathworks, Inc.) using the PARADISe software^99^.

#### Rat Aryl hydrocarbon receptor (AhR) reporter gene assay

Stably transfected rat hepatoma (H4IIE-CALUX) cells provided by Dr. Michael Denison (University of California, USA) were used. The assay was conducted as previously described^100^, where cells were incubated for ∼22 h in Minimum Essential Medium (MEM)α with 1% foetal bovine serum (FBS) and 1% penicillin/streptomycin/fungizone. Chemical exposure was performed for 24 h, and successively luminescence was measured. Cell viability was analysed by measuring ATP levels with the CellTiter-Glo® Luminescent Assay according to the manufacturer’s instruction (Promega, Denmark). 2,3,7,8-Tetrachlorodibenzo-p-dioxin (TCDD) was used as a positive control. Three experiments in triplicates were conducted with five 2-fold dilutions of ILA and IAld ranging from 12.5 to 200 μM with a constant vehicle concentration in all wells. Further, sterile filtered faecal water (10 mg faeces/ml MQ water) obtained from all samples (n=119) of 11 selected CIG infants (**Fig. 3f-h**) were run in technical triplicates in the assay. Only mild toxicity that did not correlate with AhR-induced luminescence signal was observed for some faecal water samples.

#### Human AhR reporter gene assay

ILA and IAld (positive control)^13^ were tested for activation of the human AhR. AhR Reporter Cells from Indigo Biosciences (PA, USA) that include a luciferase reporter gene functionally linked to an AhR-responsive promoter were used. The assay was run according to the instructions of the manufacturer (technical manual version 6.0) with the reference agonist MeBIO as the positive control. Three experiments in triplicates were conducted with five 2-fold dilutions of ILA and IAld ranging from 12.5 to 200 μM with a constant vehicle concentration in all wells. No cytotoxicity was observed for any of the tests as determined by a resazurin toxicity assay.

#### Human HCAR3 receptor assay

The aromatic lactic acids (ILA, PLA and 4-OH-PLA) were tested for activation of the human hydroxycarboxylic acid-3 (HCAR3) receptor, which is a Gα_i_-coupled receptor (GPCR). The cAMP Hunter™ eXpress GPR109B CHO-K1 GPCR Assay for chemiluminescence detection of cAMP was used. Following ligand stimulation of cells overexpressing the HCAR3 receptor, the functional status of the receptor was monitored by measuring cellular cAMP levels using a homogeneous, competitive immunoassay based on Enzyme Fragment Complementation technology. The assay was run in agonist mode in a 96-well plate format according to the instructions of the manufacturer (DiscoveRx Corporation, Fremont, CA, USA) in the presence of 15 μM forskolin. Eleven 3-fold dilutions of ILA ranging from 0.03 to 1574 μM and of PLA and OH-PLA ranging from 0.02 to 1000 μM were tested twice in duplicates.

#### *Ex vivo* stimulation of human immune cells

Human buffy coats were acquired from the Copenhagen University Hospital (Rigshospitalet, Copenhagen) from healthy anonymous donors under approval by the local Scientific Ethics Committee. Prior written informed consent was obtained according to the Declaration of Helsinki. Blood samples were handled in accordance with guidelines put forward in the “Transfusion Medicine Standards” by the Danish Society for Clinical Immunology (www.dski.dk).

### Isolation, cell culture and stimulation of T cells

Peripheral blood mononuclear cells (PBMCs) were isolated from whole blood by density centrifugation on Lymphoprep and cryopreserved at −150°C in FBS with 10% DMSO until the day of cell culture. For cultivation, PBMCs were thawed and CD4+T cells isolated using EasySep™ Human CD4+ T Cell Isolation Kit (Stemcell, 17952) following the manufacturers protocol. In short, approximately 2.5×10^7^ PBMCs were incubated for 5 min at room temperature in 500 μL IMDM-medium containing 50 μL CD4+T cell isolation cocktail, followed by the addition of 50 μL RapidSpheres™. Subsequently, the volume was topped up to 2.5 mL with IMDM-medium, the cells placed in an EasySep™ magnet (Stemcell) and incubated at room temperature for 3 min. The pure CD4+T cell fraction was obtained by pouring the enriched non-bound cell fraction into a new tube. Enriched CD4+T cells were cultured in Th17-polarising culture medium (IMDM supplemented with 10% FCS, 20 mM Hepes (pH 7.4), 50 μM 2-mercaptoethanol, 2 mM l-glutamine, and penicillin-streptomycin (10,000 U/ ml), 30 ng/mL IL-6, 10 ng/mL IL-1β, 0.5 ng/mL TGFβ-1, 10 ng/mL IL-23, 25 μL/mL ImmunoCult™ Human CD3/CD28 T cell activator for 3 days at 37°C and 5% CO_2_ in Falcon^TM^ polystyrene 48 well plates (Thermo Fisher, 10059110). Each culture condition contained 0.2% DMSO with or without the indicated amounts of ILA and/or the AhR-inhibitor CH-223191. After 3 days of culture, supernatants were collected for ELISA and frozen down until further use. The ELISA to detect IL-22 was performed in technical duplicates using the ELISA MAX^TM^ Deluxe Set Human IL-22 kit (Biolegend, 434504) following the supplied manufacturer’s protocol. In short, a Nunc MaxiSorb^TM^ flat bottom 96 well plate (Thermo Fischer, 44-2404-21) was coated for 12h at 4°C with IL-22 coating antibody followed by 4 rounds of washing with PBS+0.05% Tween-20. The washed plate was blocked with supplied assay diluent A buffer for 1h at room temperature and 400rpm, washed 4 more times with PBS+0.05% Tween-20 and incubated with cell culture supernatants for 2h at room temperature and 400rpm. Serially diluted standard controls and a blank control were included as a reference. To detect bound IL-22, the plate was washed 4 times with PBS+0.05% Tween-20 and incubated with IL-22 detection antibody for 1h at room temperature and 400rpm. After 4 further washing steps with PBS+0.05% Tween-20, Avidin-HRP was added for 30min at room temperature and 400rpm. To detect HRP activity, the plate was washed 5 times with PBS+0.05% Tween-20 followed by an incubation with Solution F substrate solution in the dark at room temperature. HRP activity was stopped after 20min using 1M H2SO4 and the optical density recorded (absorption at 450nm) using a PowerWave HT Microplate Spectrophotometer (BioTek Instruments). Values below limit of detection (16 pg/ml) of the kit were set to LOD/2. Sources and identifiers of all reagents used are given in **Supplementary Table 9**.

### Isolation, cell culture and stimulation of monocytes

PBMCs were isolated by Ficoll-Paque (GE Healthcare) density centrifugation. Monocytes were isolated to > 92% purity using the Pan Monocyte Isolation kit (Miltenyi Biotec). Cells were stained with CD14-PE-Cy7 (eBiosciences) and CD16-FITC (Biolegend) to determine the purity of monocytes by flow cytometry (BD FACS Canto II). Monocytes were cultured in culture medium (RPMI 1640 (Lonza) containing 2 mM L-glutamine (Lonza), 10% heat-inactivated fetal bovine serum (Lonza), 1% penicillin/streptomycin (Lonza), 50 μM 2-ME (Sigma-Aldrich)) in a humidified 37°C, 5% CO_2_ incubator. Cells were added different compounds, with the final concentrations of ILA (Sigma I5508) at 5, 50 or 200 μM, dissolved in a maximum of 0.1% DMSO. Controls were added 0.1% DMSO in culture medium. All cells were then added LPS (TLR4 ligand, *E. coli* O26:B6, Sigma L2654) at 100 ng/mL (final conc.) and IFN-gamma (RD285-IF-100) at 10 ng/mL (final conc.) and stimulated for 18 hours. To determine the contribution of AhR, monocytes were pre-treated 1 hour before addition of above compounds with the AhR antagonist CH-223191 (Sigma C8124) at 10 μM. For HCAR3 silencing, where no specific antagonist is available, we performed reverse transfection using Lipofectamine RNAiMAX in Opti-MEM (Life technologies) added 10 nM of scramble siRNA (ThermoFisher 4390846) or HCAR3-specific siRNA (ThermoFisher 4427037), reaching a maximum knockdown of 79% based on qPCR validation (HCAR3 primer (ThermoFisher 4448892) vs GAPDH) as compared to scramble siRNA targeted cells. Monocytes were pre-treated with the siRNA constructs for 24 hours before stimulation as above. Viability of cells was >97 % as analysed by flow cytometry (BD FACS Canto II) using SYTOX AAD. Supernatants of stimulated cells were harvested after 18 hrs and kept at −80°C until analysis. IL-12p70 was quantified using Meso Scale Discovery kits as previously detailed^101^.

#### Statistical analyses

Statistical analyses were performed using QIIME v1.9^78^, R v3.1^102^ and GraphPad Prism v8.1 (GraphPad Software, Inc. CA). PCoA, ADONIS and PERMDISP tests (permutations = 999) of OTU distance/dissimilarity matrices were performed in QIIME, and PCoA plots were illustrated in R using the *ggplot2* package^103^. PCA was performed in R using the *ggbiplot* package^104^. Spearman’s Rank correlations were performed in GraphPad Prism, whereas partial Spearman’s Rank correlation analyses with adjustments for age and repeated measures correlation analyses were performed in R using the *ppcor*^30^ and *rmcorr* packages^29^, respectively. Heatmaps and hierarchically clustering of correlation coefficient were generated in R using the *gplots* package^105^ and visualised in GraphPad Prism. Longitudinal metabolite and taxonomic abundance were modelled using LOESS regression, and implemented and plotted with 95% confidence intervals in R using the *ggplot2* package^103^. Longitudinal AhR activity was modelled used coarse LOWESS curve fits within the GraphPad Prism software. Two-tailed paired or unpaired Student’s *t* test (if normally distributed, evaluated by D’Agostino-Pearson test) or two-tailed non-parametric Mann-Whitney *U* test (if not normally distributed) were performed when comparing two groups. For comparison of more than two groups, statistical significance was evaluated by one-way ANOVA (if normally distributed) or the non-parametric Kruskal-Wallis test (if not normally distributed). P-values < 0.05 were considered statistically significant. When applicable p-values were corrected for multiple testing by the Benjamini–Hochberg false discovery rate (FDR)^106^ using a cutoff of 0.1.

## ACKNOWLEDGMENTS

We thank the children and families participating in the SKOT I study, which was supported by the Danish Directorate for Food, Fisheries and Agribusiness (Grant no. 3304-FSE-06-0503), as well as the children and parents participating in the CIG cohort. Furthermore, we thank Aarstiderne A/S for providing a small gift for the CIG participants. We thank Marlene Danner Dalgaard at the Technical University of Denmark in-house facility (DTU Multi-Assay Core, DMAC) for performing the 16S rRNA gene sequencing, MS-Omics (Hørsholm, Denmark) for performing the lactate analyses, Glycom A/S (Hørsholm, Denmark) for kindly donating the human milk oligosaccharides, and Anette Schnipper for her efforts in supporting the work. We also thank Satoru Fukiya and Atsushi Yokota for providing *Bifidobacterium* gene manipulation tools and for technical suggestions, Yuta Sugiyama for helpful discussions on HPLC analysis, Shingo Maeda for technical support on enzyme purification, technician Lisbeth Buus Rosholm for performing the human monocyte experiments, and technician Birgitte Møller Plesning for running the AhR and HCAR-3 assays. This work was supported by Augustinus Fonden (grant no. 17-2003 to H.M.R.), Hørslev Fonden (grant no. 203866 to H.M.R.), Beckett Fonden (grant no. 17-2-0551 to H.M.R.), Aase og Ejnar Danielsens Fond (grant no. 10-002019 to H.M.R.), the Innovation Fund Denmark (grant no. 11-116163/0603-00487B; Center for Gut, Grain and Greens to T.R.L.), JSPS-KAKENHI (18K14379 to M.S., 19K22277 to T.K.), JSPS Overseas Research Fellowships (201860637 to M.S.) and the Institute for Fermentation, Osaka (to M.S. and T.K.), “Diet-induced Arrangement of the gut Microbiome for improvement of Cardiometabolic health” (DINAMIC) under the Joint Programming Initiative, “A Healthy Diet for a Healthy Life” (JPI-HDHL), supported by the Innovation Fund Denmark (grant no. 5195-00001B to L.O.D.) and “Biomarkers for Infant Fat Mass Development and Nutrition” under the ERA-HDHL joint transnational program “Biomarkers for Nutrition and Health”, supported by the Innovation Fund Denmark (grant no. 4203-00005B to S.B.).

## AUTHOR CONTRIBUTIONS

H.M.R. and M.F.L. conceived and designed the study. M.F.L. prepared the samples for sequencing/qPCR and analysed the sequencing/qPCR data. H.M.R. prepared the samples for faecal and *in vitro* metabolome analyses and performed together with H.L.F. the targeted and untargeted metabolomics experiments. C.T.P. and L.O.D. performed the urine metabolomics. M.F.L. and M.S. performed the *in vitro* growth and mutant construction experiments. M.S. and T.K. performed enzyme kinetics. M.F.L. and H.M.R. performed *in vivo* experiments. K.F.M. and C.M. designed the SKOT I study and M.V.L. managed the data. H.M.R. and M.F.L. designed the CIG cohort, recruited the study participants and managed the data. A.M.V. performed the AhR and HCAR3 reporter assays. J.M.M. performed bioinformatics analysis of strain variation and MAGs. U.M. and S.B. performed *ex vivo* immune experiments. N.B., D.A., and A.R. contributed to the interpretation of the results. S.B., W.A., T.K., M.I.B. and T.R.L. contributed with expert supervision. H.M.R. and M.F.L. led the work, undertook the integrative data analyses and drafted the manuscript. All authors contributed to and approved the final manuscript.

## Data statement

Partial (V3 region) 16S rRNA gene amplicon sequencing data is deposited in the Sequence Read Archive (SRA) under the BioProjects PRJNA273694 (SKOT) and PRJNA554596 (CIG).

## Competing Financial Interests statement

The authors declare no competing financial interests.

## Corresponding authors

\# Henrik Munch Roager

\# Tine Rask Licht

## Notes

### Competing Interest Statement

The authors have declared no competing interest.

