## Supplementary Materials for "Breastmilk-promoted bifidobacteria produce aromatic amino acids in the infant gut"

**Contains:**

**Supplementary Table 1-9**

**Supplementary Figures 1-20**

### Supplementary tables

**Supplementary Table 1**. Aromatic amino acids and derivatives and their retention time, mass-to-charge ratio (*m/z*) and internal standard.

| **Pathway**[1] | **Metabolite** | **Retention time [min]** | ***m/z*** | **ESI$^+$ ion** | **Internal standard** |
|---|---|---|---|---|---|
| Phenylalanine | Phenylalanine | 2.6 | 166.0863 | $[M+H]^+$ | L-Phenylalanine (ring-d5) |
| Phenylalanine | Phenyllactic acid | 4.7 | 189.0522 | $[M+Na]^+$ | L-Phenylalanine (ring-d5) |
| Phenylalanine | Phenylacetic acid | 5.3 | 137.0597 | $[M+H]^+$ | L-Phenylalanine (ring-d5) |
| Phenylalanine | Phenylpropionic acid | 5.9 | 151.0754 | $[M+H]^+$ | L-Phenylalanine (ring-d5) |
| Tryptophan | Kynurenine | 2.5 | 209.0921 | $[M+H]^+$ | L-Tryptophan (indole-d5) |
| Tryptophan | Tryptophan | 3.4 | 205.0972 | $[M+H]^+$ | L-Tryptophan (indole-d5) |
| Tryptophan | Xanthurenic acid | 3.6 | 206.0448 | $[M+H]^+$ | L-Tryptophan (indole-d5) |
| Tryptophan | Tryptamine | 3.6 | 161.1073 | $[M+H]^+$ | L-Tryptophan (indole-d5) |
| Tryptophan | Kynurenic acid | 3.9 | 190.0499 | $[M+H]^+$ | L-Tryptophan (indole-d5) |
| Tryptophan | Indolelactic acid | 5.0 | 206.0812 | $[M+H]^+$ | Indoleacetic acid (2,2-d2) |
| Tryptophan | Indolealdehyde | 5.2 | 146.0600 | $[M+H]^+$ | Indoleacetic acid (2,2-d2) |
| Tryptophan | Indoleacetic acid | 5.3 | 176.0706 | $[M+H]^+$ | Indoleacetic acid (2,2-d2) |
| Tryptophan | Indoleethanol | 5.4 | 162.0913 | $[M+H]^+$ | Indoleacetic acid (2,2-d2) |
| Tryptophan | Indolepropionic acid | 5.9 | 190.0863 | $[M+H]^+$ | Indoleacetic acid (2,2-d2) |
| Tyrosine | Tyrosine | 1.3 | 182.0812 | $[M+H]^+$ | L-Tyrosine (ring-d4) |
| Tyrosine | Tyramine | 1.3 | 138.0913 | $[M+H]^+$ | L-Tyrosine (ring-d4) |
| Tyrosine | 4-Hydroxyphenyllactic acid | 3.5 | 205.0471 | $[M+Na]^+$ | L-Tyrosine (ring-d4) |
| Tyrosine | 4-Hydroxyphenylacetic acid | 3.9 | 153.0546 | $[M+H]^+$ | L-Tyrosine (ring-d4) |
| Tyrosine | 4-Hydroxyphenylpropionic acid | 4.6 | 189.0522 | $[M+Na]^+$ | L-Tyrosine (ring-d4) |
| *IS* | L-Phenylalanine (ring-d5) | 2.6 | 171.1176 | $[M+H]^+$ | - |
| *IS* | L-Tryptophan (indole-d5) | 3.4 | 210.1285 | $[M+H]^+$ | - |
| *IS* | L-Tyrosine (ring-d4) | 1.3 | 186.1063 | $[M+H]^+$ | - |
| *IS* | Indoleacetic acid (2,2-d2) | 5.3 | 178.0832 | $[M+H]^+$ | - |

[1]IS, internal standard

**Supplementary Table 2.** Identification of type 4 *ldh* in whole genome sequenced human gut derived *Bifidobacterium* species.

| **Species** | **Number of strains** | **Number of strains with type 4 *ldh*** | **Percent of strains with type 4 *ldh*** |
|---|---|---|---|
| *B. breve* | 44 | 44 | 100 % |
| *B. longum* | 27 | 27 | 100 % |
| *B. animalis* | 24 | 0 | 0 % |
| *B. bifidum* | 9 | 9 | 100 % |
| *B. adolescentis* | 8 | 0 | 0 % |
| *B. kashiwanohense* | 3 | 0 | 0 % |
| *B. pseudolongum* | 3 | 0 | 0 % |
| *B. angulatum* | 2 | 2 | 100 % |
| *B. catenulatum* | 2 | 0 | 0 % |
| *B. dentium* | 2 | 0 | 0 % |
| *B. pseudocatenulatum* | 2 | 0 | 0 % |
| *B. scardovii* | 1 | 1 | 100 % |

**Supplementary Table 3.** Faecal concentration of aromatic amino acid metabolites in the Copenhagen Infant Gut cohort (n=267).

| Metabolite | Prevalence (%) | Concentration (µmol/mg faeces) | | | |
|---|---|---|---|---|---|
| | | Median [IQR] | Mean [±SD] | Min | Max |
| *Tyrosine metabolites* | | | | | |
| *Tyrosine* | 100.0 | 307.4 [194.9-457.9] | 358.1 [±231.8] | 26.6 | 1776.0 |
| *Tyramine* | 63.7 | 11.0 [0.0-194] | 185.1 [±363.7] | 0.0 | 2428.0 |
| *4-hydroxyphenyllactic acid* | 93.3 | 40.6 [10.9-76.6] | 53.5 [±54.3] | 0.0 | 292.1 |
| *4-hydroxyphenylacetic acid* | 44.9 | 0.0 [0.0-41.9] | 51.7 [±125.7] | 0.0 | 995.3 |
| *4-hydroxyphenyl-propionic acid* | 16.1 | 0.0 [0.0-0.0] | 13.5 [±64.6] | 0.0 | 733.2 |
| *Phenylalanine metabolites* | | | | | |
| *Phenylalanine* | 100.0 | 381.1 [267-497.4] | 419.9 [±250.0] | 37.1 | 2159.0 |
| *Phenyllactic acid* | 97.8 | 38 [17.6-73.4] | 54.9 [±56.7] | 0.0 | 477.4 |
| *Phenylacetic acid* | 29.2 | 0.0 [0.0-9.6] | 24.5 [±76.6] | 0.0 | 718.2 |
| *Phenylpropionic acid* | 10.1 | 0.0 [0.0-0.0] | 10.9 [±49.1] | 0.0 | 601.6 |
| *Tryptophan metabolites* | | | | | |
| *Tryptophan* | 100.0 | 128.3 [84.6-197.1] | 155.7 [±104.0] | 10.6 | 647.8 |
| *Tryptamine* | 25.8 | 0.0 [0.0-0.1] | 4.0 [±13.2] | 0.0 | 104.2 |
| *Kynurenine* | 82.4 | 0.3 [0.1-0.5] | 0.4 [±0.4] | 0.0 | 2.3 |
| *Kynurenic acid* | 100.0 | 5.9 [3.4-9.0] | 8.1 [±8.7] | 0.9 | 90.8 |
| *Xanthurenic acid* | 98.5 | 0.7 [0.5-1.2] | 1.0 [±0.9] | 0.0 | 6.3 |
| *Indolelactic acid* | 98.1 | 47.7 [15.0-103.4] | 69.0 [±70.2] | 0.0 | 332.2 |
| *Indoleacetic acid* | 47.2 | 0.0 [0.0-1.0] | 2.1 [±9.9] | 0.0 | 127.2 |
| *Indolepropionic acid* | 7.1 | 0.0 [0.0-0.0] | 0.5 [±3.2] | 0.0 | 39.0 |
| *Indolealdehyde* | 85.4 | 0.9 [0.5-2.3] | 2.1 [±3.4] | 0.0 | 23.0 |
| *Indoleethanol* | 4.5 | 0.0 [0.0-0.0] | 0.1 [±0.3] | 0.0 | 3.2 |

**Supplementary Table 4.** Primers used for quantitative PCR.

| Primer target | Primer name | Sequence (5'-3') | Final conc. (μM) | Annealing temperature (°C) | Reference |
|---|---|---|---|---|---|
| **Universal bacteria** | PBU | CCTACGGGAGGCAGCAG | 0.2 | 60 | Tulstrup *et al.*, 2015[1] |
| | PBR | ATTACCGCGGCTGCTGG | 0.2 | | |
| ***B. longum* subsp. *infantis*** | Blon0915F | CGTATTGGCTTTGTACGCATTT | 0.75 | 50 | Frese *et al.*, 2017[2] |
| | Blon0915R | ATCGTGCCGGTGAGATTTAC | 0.75 | | |
| ***B. longum* subsp. *longum*** | lon_0274_F | GAGGCGATGGTCTGGAAGTT | 0.75 | 50 | Lawley *et al.*, 2017[3] |
| | lon_0274_R | CCACATCGCCGAGAAGATTC | 0.75 | | |

**Supplementary Table 5.** *Bifidobacterium* strains used in this study.

| **Strain** | **Culture Collection**[1] |
|---|---|
| *B. adolescentis* E194a | DSM $20083^T$ |
| *B. animalis* subsp. *lactis* UR1 | DSM $10140^T$ |
| *B. animalis* subsp. *animalis* R101-8 | DSM $20104^T$ |
| *B. bifidum* Ti | DSM $20456^T$ |
| *B. breve* S1 | DSM $20213^T$ |
| *B. dentium* B764 | DSM $20436^T$ |
| *B. longum* subsp. *longum* E194b | DSM $20219^T$ |
| *B. longum* subsp. *longum* 105-A | JCM 31944 |
| *B. longum* subsp. *longum* 105-A type 4 *ldh*::pMSK127 | - |
| *B. longum* subsp. *longum* 105-A type 4 *ldh*::pMSK127 / pMSK128 (P*xfp*-type4 *ldh*) | - |
| *B. longum* subsp. *infantis* S12 | DSM $20088^T$ |
| *B. pseudolongum* subsp. *pseudolongum* PNC-2-9G | DSM $20099^T$ |
| *B. scardovii* CCUG 13008 A | DSM $13734^T$ |
| *B. catenulatum* B669 | DSM $16992^T$ |
| *B. pseudocatenulatum* B1279 | DSM $20438^T$ |

[1]Superscript “T” indicates type strain

**Supplementary Table 6.** Primers used for generation of type 4 *ldh* insertional mutant and type 4 *ldh* complemented strain.

| Primer name | Primer sequence (5′-nucleotide sequence-3′)[1] |
|---|---|
| Pr-24 | agatttcattggcttctaaattttt |
| Pr-174 | gaacgacctacaccgaactgag |
| Pr-543 | TATATATGAGTACTGaggtgcagcgcgtgctcctt |
| Pr-546 | CAGGCATGCAAGCTTtggcctacgaacctgtctgg |
| Pr-579 | CGGTACCCGGGGATCtacgatgccggccacggtg |
| Pr-580 | CCAGCTCAAGGGATCgcccgcgacctcgatgatg |
| Pr-581 | CGGTACCCGGGGATCgtggacacggaggcgaaac |
| Pr-598 | GGAAGATCACTTCGCttcgaacatgggaaatcgaa |
| Pr-599 | GTTCATAGTGACCATaagcactccttgggggccgc |
| Pr-600 | atggtcactatgaaccgcaacaaag |
| Pr-601 | CGTGGGATCCGTCGAtcacagcagcccctcgcagt |

[1]Uppercase letter indicates sequence for In-Fusion cloning

**Supplementary Table 7**. Human milk oligosaccharides (HMOs), their retention time and mass-to-charge ratio (*m/z*).

| **Abbreviation** | **HMO** | **Retention time [min]** | **[*m/z*]** | **ESI$^+$ ion** |
|---|---|---|---|---|
| 2’FL / 3FL | 2'-*O*-Fucosyllactose / 3-*O*-Fucosyllactose | 0.6 | 489.1814 | $[M+H]^+$ |
| LNT / LN*n*T | Lacto-*N*-tetraose / Lacto-*N*-*neo*tetraose | 0.7 | 708.2557 | $[M+H]^+$ |
| 3’SL / 6’SL | 3'-*O*-Sialyllactose / 6'-*O*-Sialyllactose | 0.7 | 634.2189 | $[M+H]^+$ |

**Supplementary Table 8.** Optimized parameters used for pre-processing the urine metabolome data in mzMine2.8

| Batch step | | Parameters |
|---|---|---|
| Negative Mode | Mass detection | Noise level: 1.0E1 |
| | Chromatogram builder | Min time span (min): 0.01<br>Min height:3.0E1 ; *m/z* tolerance: 0.06 mz or 25 ppm |
| | Chromatogram deconvolution | Chromatographic threshold: 90%; Search minimum in RT range (min): 0.01; Minimum relative height: 5%; Minimum absolute height: 3.0E1; Min ratio of peak/top edge: 1.0; Peak duration range (min): 0.01-0.2 |
| | Isotopic pattern | *m/z* tolerance: 0.06 or 30 ppm; Retention time tolerance: 0.02; Monotonic shape; maximum charge: 1 |
| | Join aligner | *m/z* tolerance: 0.03 or 20 ppm; Absolute retention time tolerance: 0.2; Weight for both *m/z* tolerance and retention time tolerance: 10 |
| | Peak finder (gap filling) | Intensity tolerance: 50%; *m/z* tolerance: 0.03or 20 ppm; Absolute retention time tolerance: 0.03 |

**m/z*: mass to charge ratio; RT: retention time; Intensity: Peak area

**Supplementary Table 9.** Reagents used for human T-cell cultures.

| Reagent | Source | Cat no. / RRID |
|---|---|---|
| Triton X-100 | Sigma-Aldrich | T8787 |
| Paraformaldehyde (PFA) | Sigma-Aldrich | P6148 |
| Low melting point agarose | Invitrogen | 16520-100 |
| Fetal Bovine Serum (FBS) | Gibco | ADD |
| 4´,6´-diamidino-2-phenylindole (DAPI) | Thermo Fisher | D1306 |
| Tween-20 | Sigma-Aldrich | P1379 |
| Gibco™ DPBS | Thermo Fisher | 14-190-144 |
| ß-mercopatoethanol | | |
| Hepes | Gibco | |
| L-glutamine | Gibco | |
| Penicillin / Streptomycin | Thermo Fisher | 15140122 |
| $H_2SO_4$ | Sigma-Aldrich | 339741 |
| IL-6 | R&D | 206-IL-050 |
| TGFβ | R&D | 240-B-010 |
| IL-23 | R&D | 1290-IL-010 |
| IL-1β | R&D | 201-LB-010 |
| ImmunoCult™ Human CD3/CD28 T Cell Activator | Stemcell | 10971 |
| Lymphoprep | Axis Shield Poc As | 11508545 |

### Supplementary figures

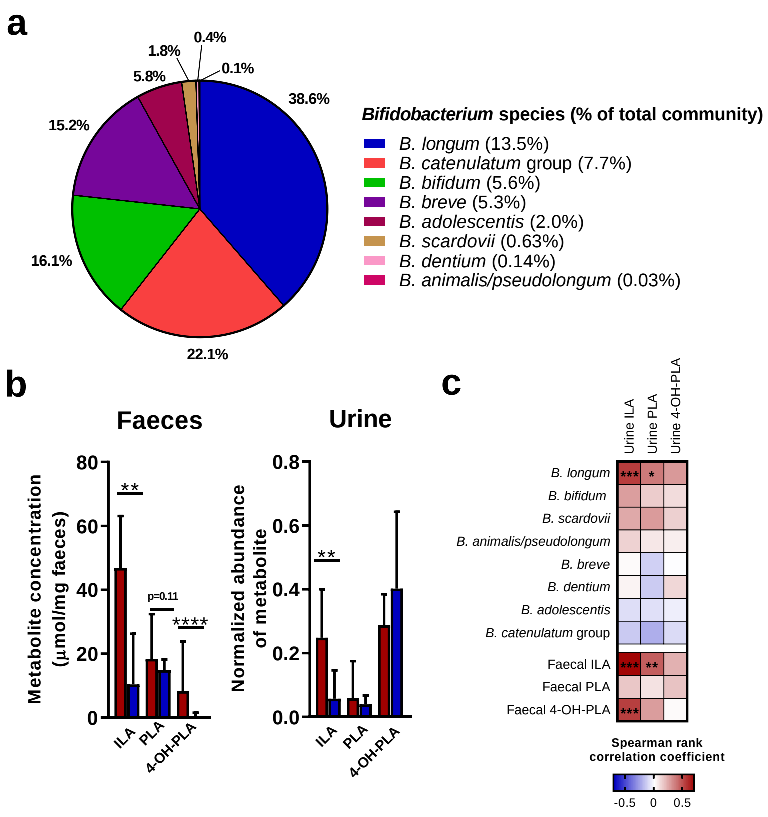


**Supplementary Figure 1. *Bifidobacterium* species composition and aromatic lactic acids in partially breastfed and weaned infants aged 9 months from the SKOT cohort.**

**a**, Pie chart of the average *Bifidobacterium* species composition given as percent of total *Bifidobacterium* abundance (n=59 infants). Legend includes the average percent of each species compared to the total gut microbiota community (See also **Supplementary Data 1f**). **b**, Left panel: Barplots with median+95%CI of faecal levels of ILA, PLA and 4-OH-PLA in partially breastfed (n = 24, red) and weaned (n = 35, blue) infants. Right panel: Barplots with median+95%CI of urine levels of ILA, PLA and 4-OH-PLA in partially breastfed infants (n = 19, red) and weaned (n = 30, blue). Statistical significance was evaluated by Mann-Whitney *U* test. **c**, Heatmap of Spearman's Rank correlation coefficients between the relative abundances of indolelactic acid (ILA), phenyllactic acid (PLA) and 4-hydroxyphenyllactic acid (4-OH-PLA) measured in urine and faecal relative abundance of *Bifidobacterium* species or faecal concentrations of ILA, PLA and 4-OH-PLA of the same infants (n=49 infants), *p<0.05, **p<0.01, ***p<0.001 and ****p < 0.0001.

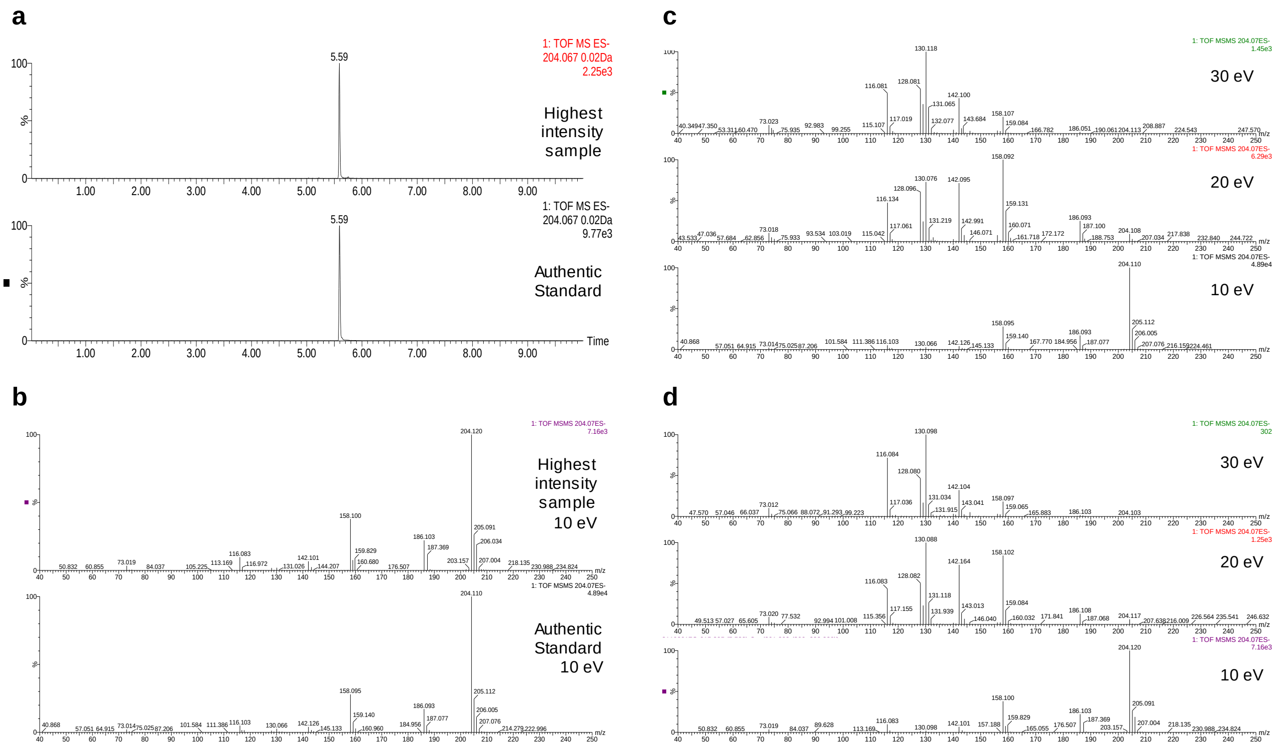


**Supplementary Figure 2. Identification of indolelactic acid in urine by extracted ion chromatograms and fragmentation profiles of indolelactic acid and sample with the highest intensity.**

**a**, Feature defined with 204.067 *m/z* and 5.59 RT in negative ionization mode confirmed with authentic standard of indole lactic acid. **b**, MS/MS fragmentation profile of authentic standard and the highest intensity sample fragmented at 10 eV collision dissociation energy. **c**, MS/MS fragmentation profile of authentic standard of indole lactic acid at 10, 20 and 30 eV collision dissociation energies. **d**, MS/MS fragmentation profile of sample with highest intensity at 10, 20 and 30 eV collision dissociation energies.

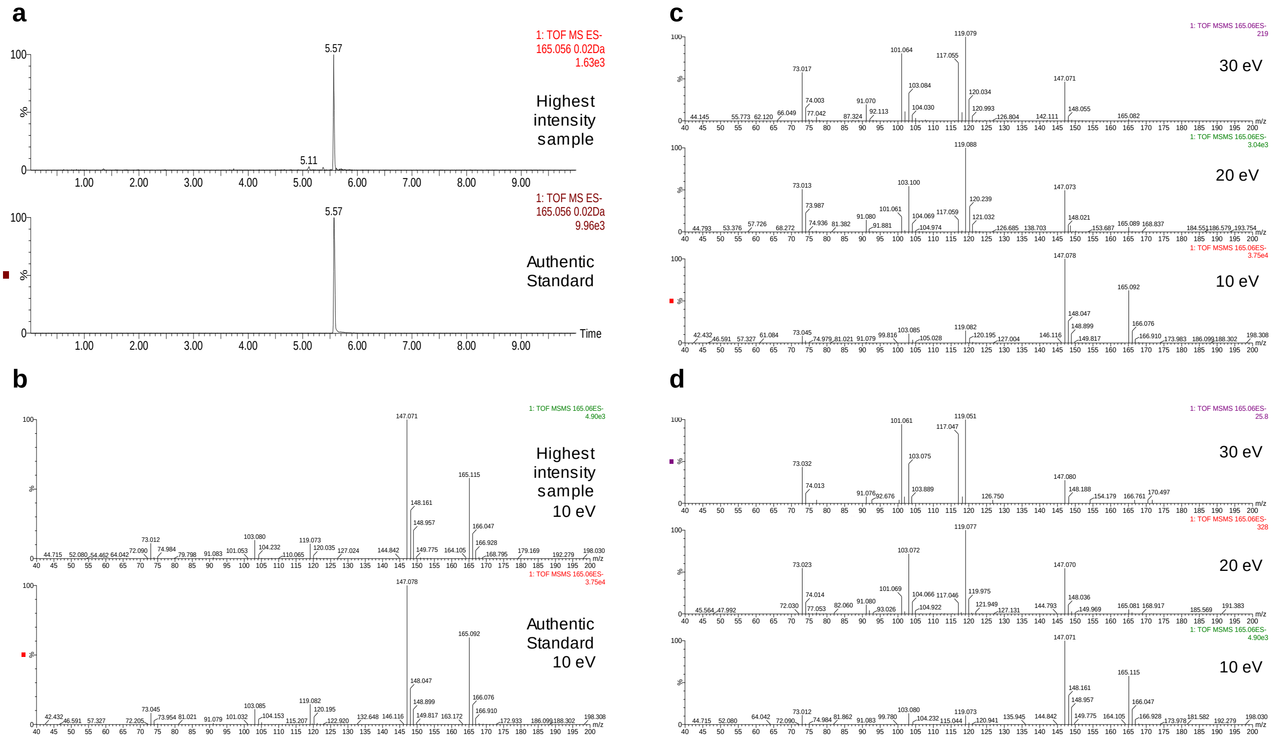


**Supplementary Figure 3. Identification of phenyllactic acid in urine by extracted ion chromatograms and fragmentation profiles of phenyllactic acid and sample with the highest intensity.**

**a**, Feature defined with 165.056 *m/z* and 5.57 RT in negative ionization mode confirmed with authentic standard of phenyl lactic acid. **b**, MS/MS fragmentation profile of authentic standard and the highest intensity sample fragmented at 10 eV collision dissociation energy. **c,** MS/MS fragmentation profile of authentic standard of phenyl lactic acid at 10, 20 and 30 eV collision dissociation energies. **d**, MS/MS fragmentation profile of sample with highest intensity at 10, 20 and 30 eV collision dissociation energies.

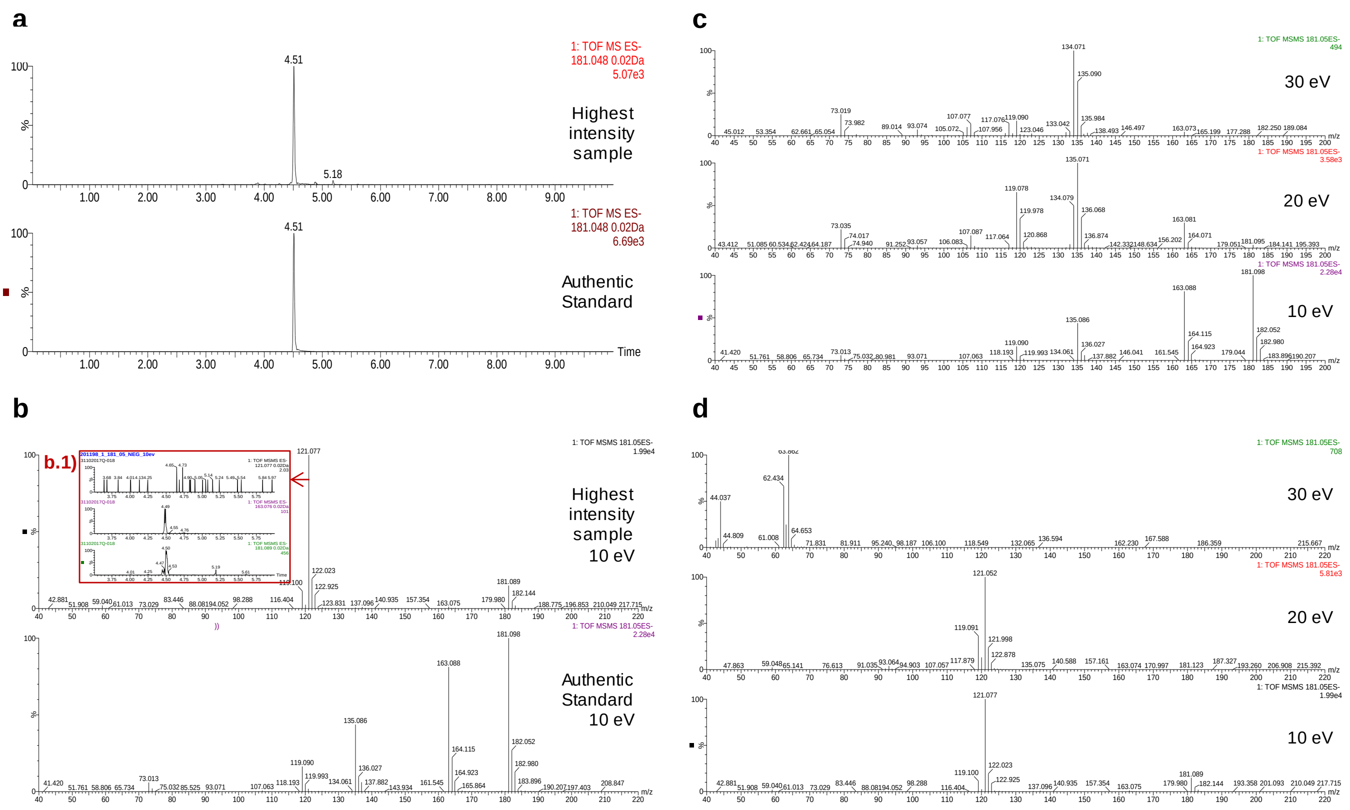


**Supplementary Figure 4. Identification of 4-hydroxyphenyllactic acid in urine by extracted ion chromatograms (EICs) and fragmentation profiles of 4-hydroxyphenyllactic acid and sample with the highest intensity.**

**a**, Feature defined with 181.048 *m/z* and 4.51 RT in negative ionization mode confirmed with authentic standard of 4-hydroxyphenyllactic acid. **b**, MS/MS fragmentation profile of authentic standard and the highest intensity sample fragmented at 10 eV collision dissociation energy. b.1) EIC of 181.089 *m/z* (parent ion), 163.075 *m/z* (main fragment) and 121.077 *m/z* shown with arrow indicates that the feature with 121.077 *m/z* does not belong to the peak of 4-hydroxyphenyllactic acid. **c**, MS/MS fragmentation profile of authentic standard of 4-hydroxyphenyllactic acid at 10, 20 and 30 eV collision dissociation energies. **d**, MS/MS fragmentation profile of sample with highest intensity at 10, 20 and 30 eV collision dissociation energies.

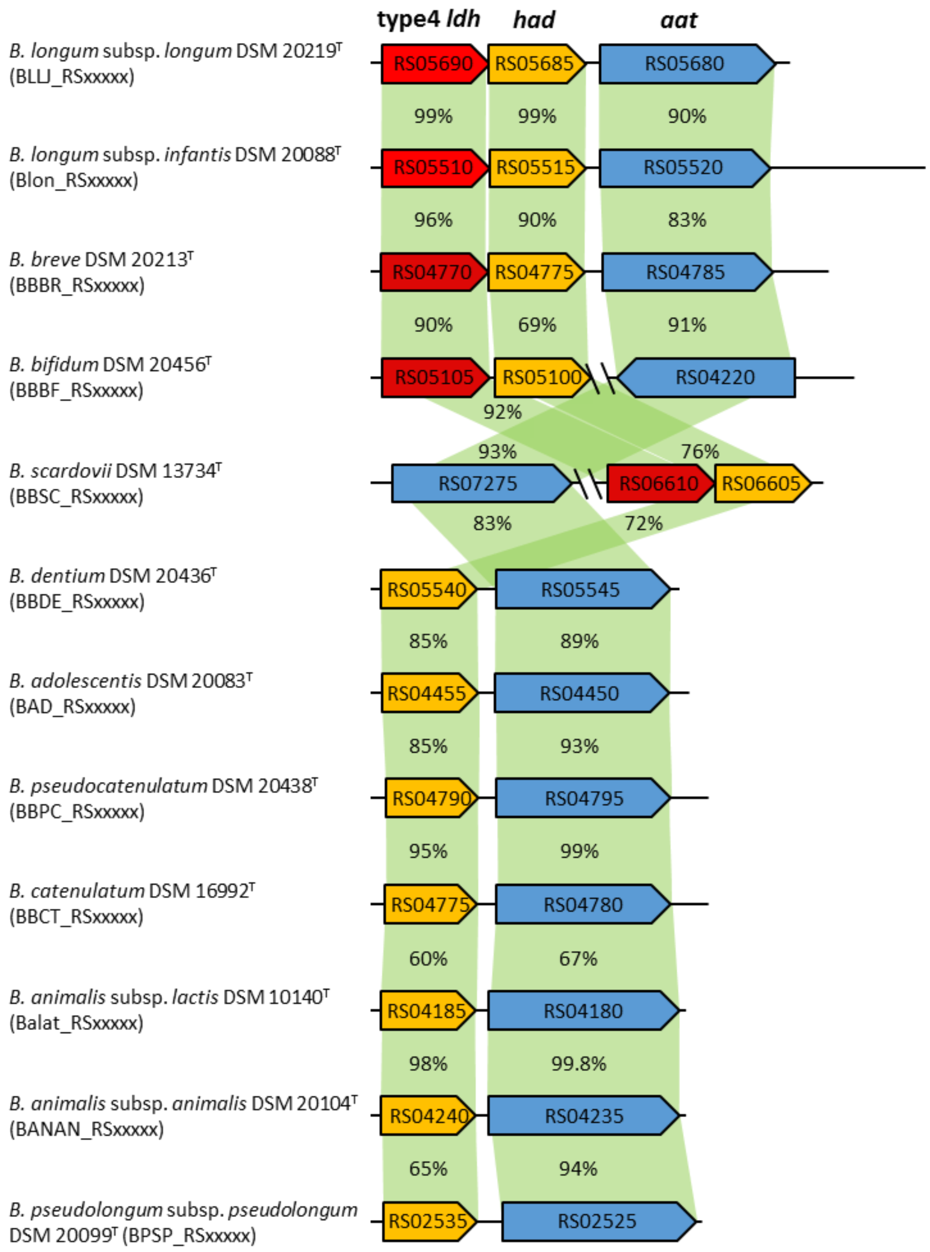


**Supplementary Figure 5. Comparison of 'type 4' *ldh* gene cluster in human-gut-associated *Bifidobacterium* type strains.** The 'type 4' *ldh*, *had*, and *aat* genes are represented by red, yellow, and blue arrows, respectively. The amino acid sequence identity was determined by conducting pairwise alignments in MBGD (Microbial Genome Database for Comparative Analysis; http://mbgd.genome.ad.jp/). *ldh*: lactate dehydrogenase gene, *had*: haloacid dehalogenase gene, *aat*: amino acid transaminase gene. The gene organization was drawn by drawGeneArrows3 (http://www.ige.tohoku.ac.jp/joho/labhome/tool.html).

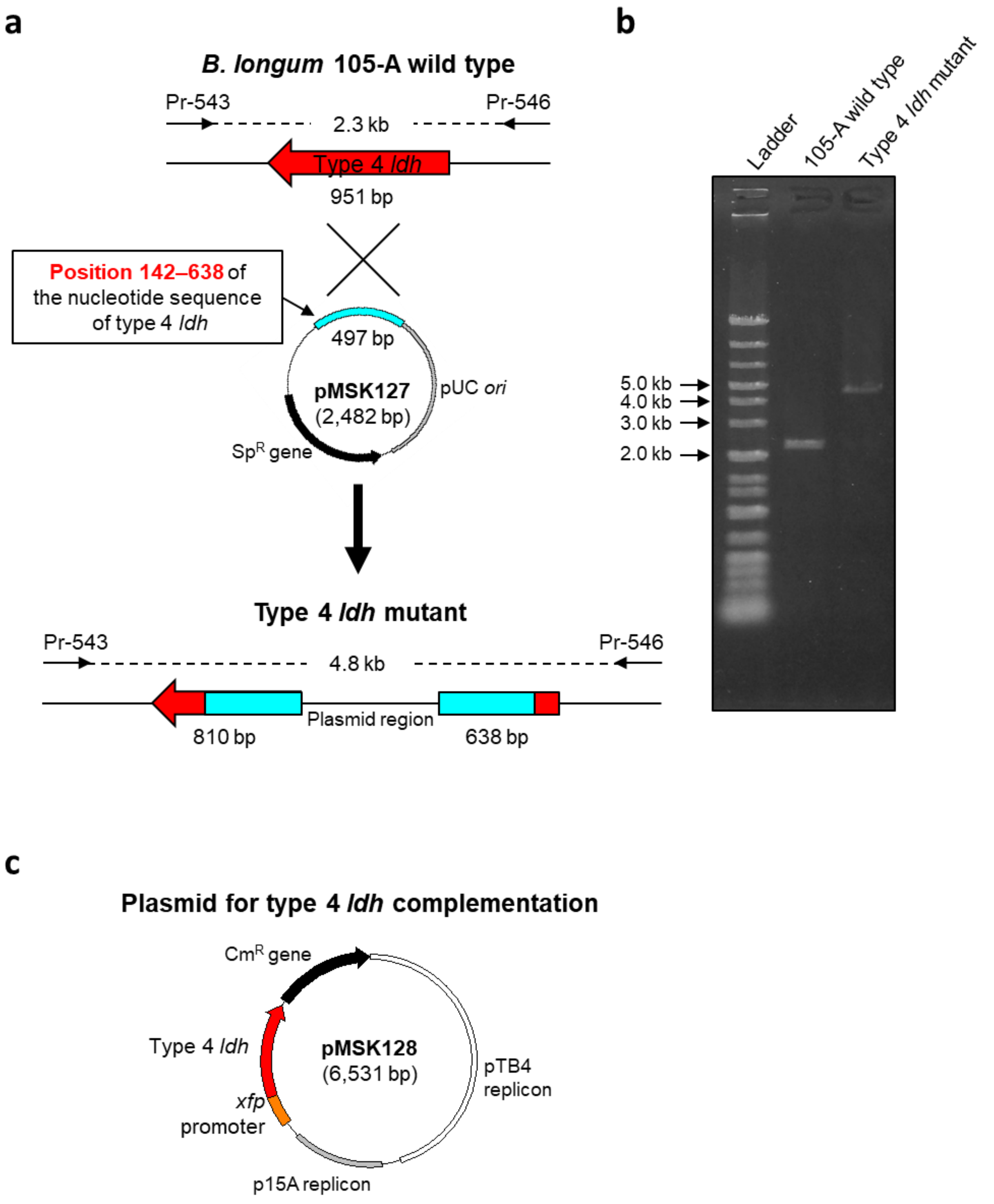


**Supplementary Figure 6. Generation of the type 4 *ldh* insertional mutant and type 4 *ldh* complemented strain. a,** Generation of type 4 *ldh* insertional mutant by single cross-over recombination with pMSK127 containing the partial *ldh* gene from *B. longum* subsp. *longum* 105-A (BL105A_0985). **b**, Insertion of pMSK127 was verified by PCR and gel electrophoresis using primiers (Pr-543/Pr-546) targeting regions flanking the *ldh* gene. **c**) pMSK128 containing the complete *ldh* gene (BL105A_0985), which was introduced into type 4 *ldh* mutant to generate complemented type 4 *ldh* strain.

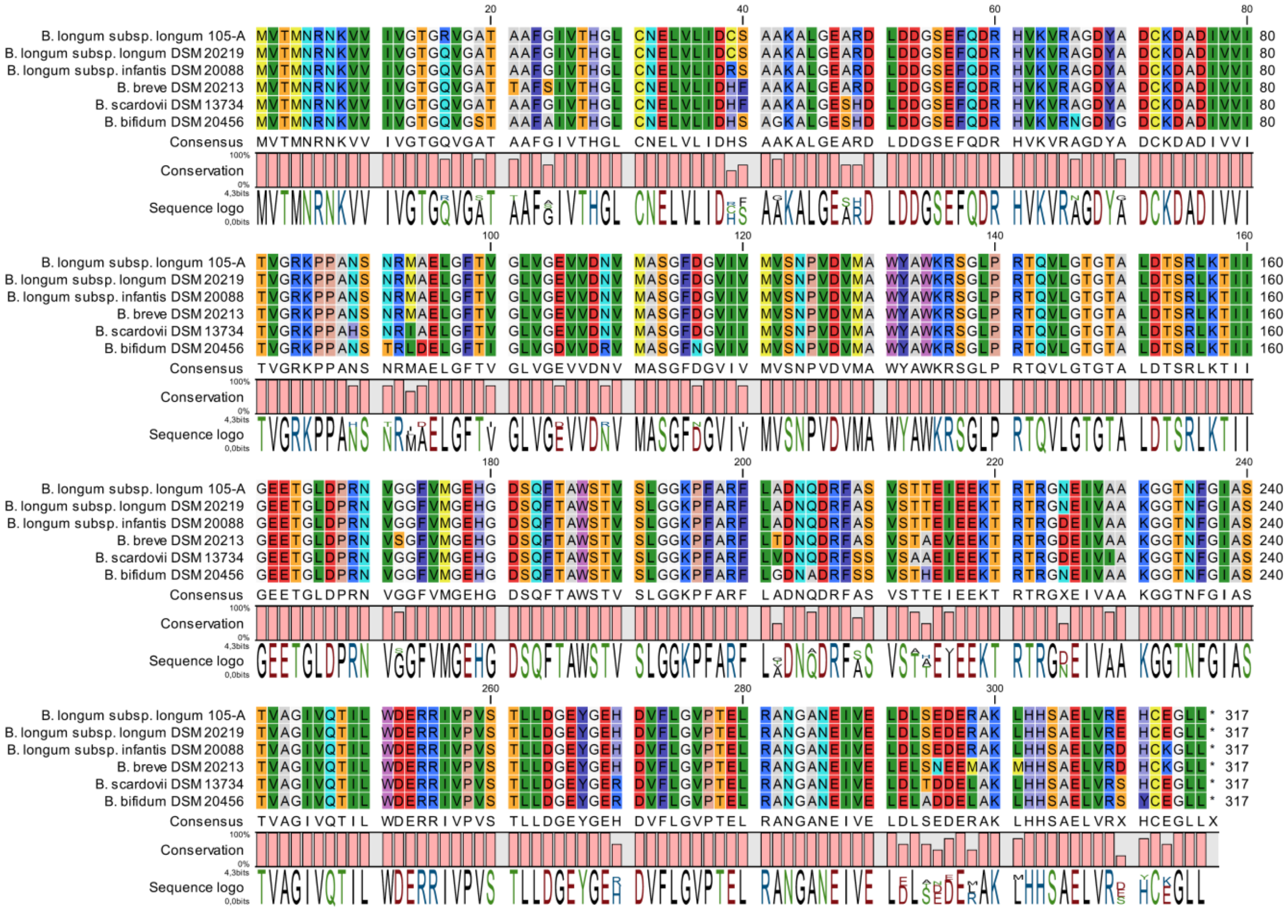

**Supplementary Figure 7. Alignment of the type 4 LDH amino acid sequences from *B. longum* subsp. *longum* 105-A and the type strains of *B. longum* subsp. *longum*, *B. longum* subsp. *infantis*, *B. scardovii*, *B. breve* and *B. bifidum*.**

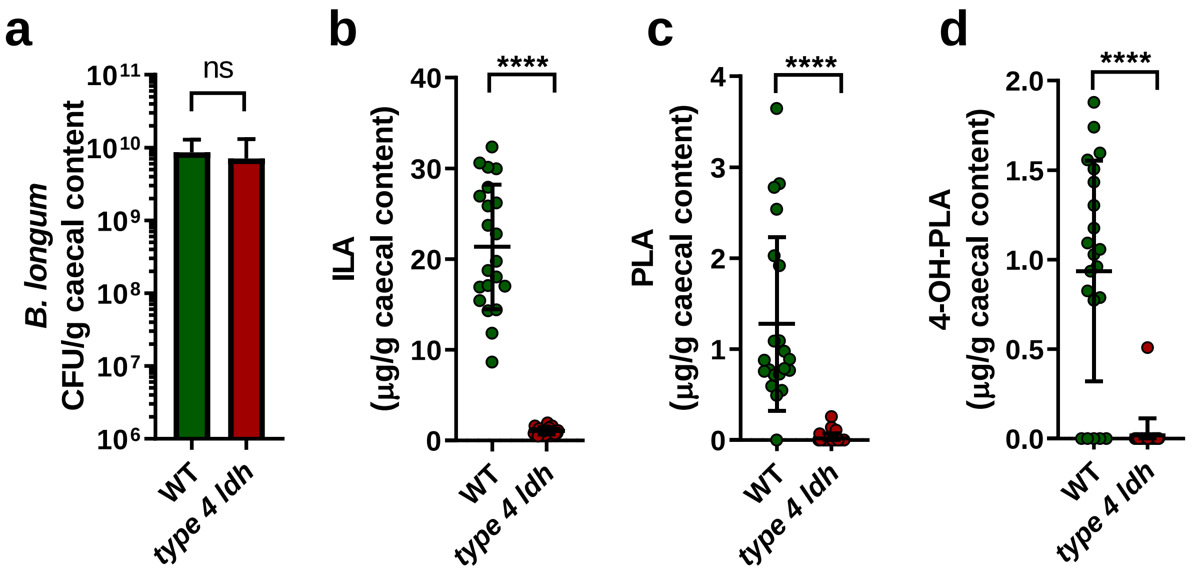


**Supplementary Figure 8.** *In vivo* production of aromatic lactic acid in previously germ free animals colonised with either *B. longum* 105-A WT or type 4 *ldh* mutant. Statistical significance was evaluated by Mann Whitney U tests, with **** $p < 0.0001$.

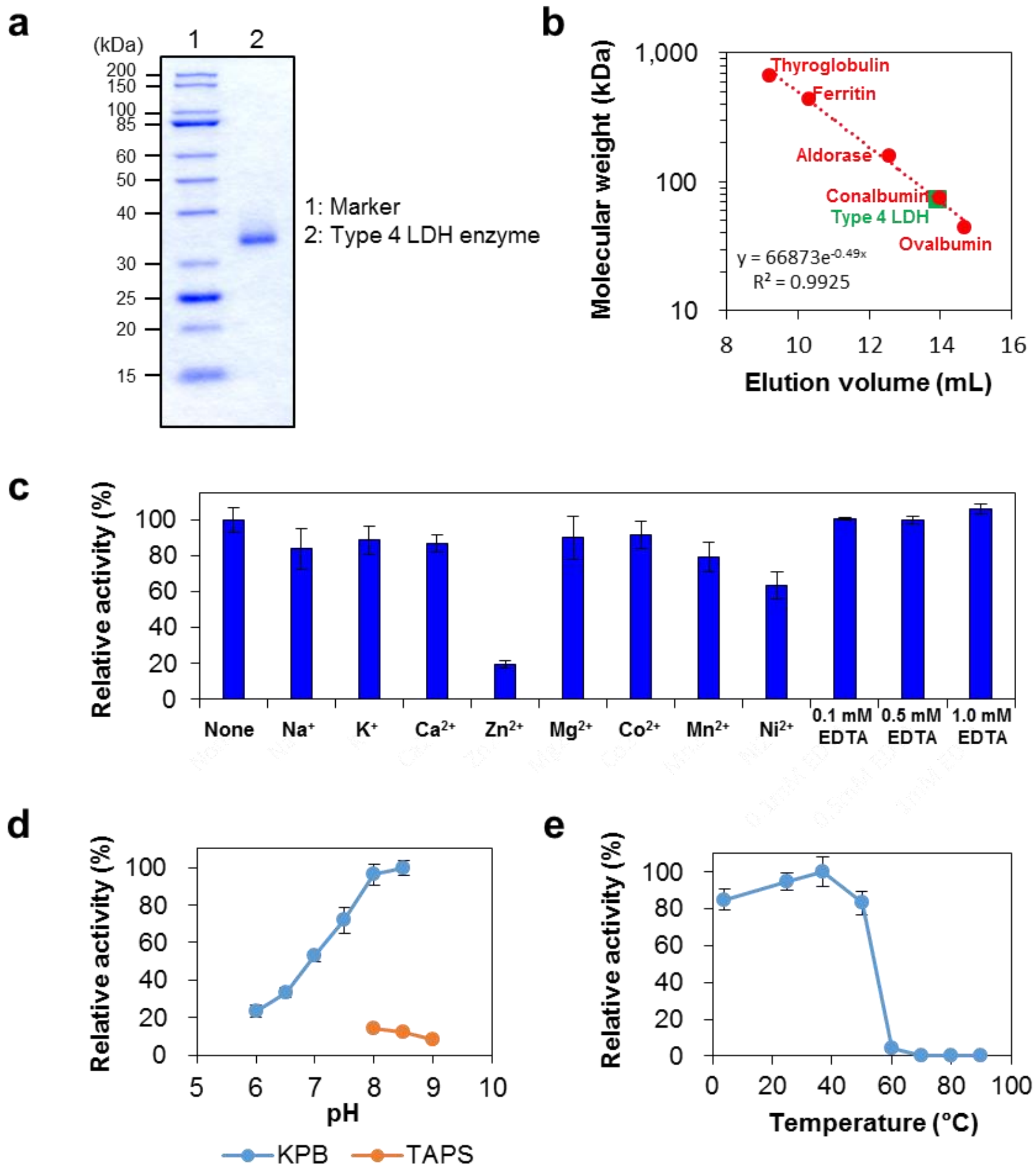


**Supplementary Figure 9. The physicochemical properties of type 4 LDH enzyme.**
**a**, Representative picture of two SDS-PAGE runs of type 4 LDH enzyme. Lane 1, Marker; Lane 2, purified type 4 LDH enzyme. **b**, The native molecular mass of the type 4 LDH enzyme was estimated by size exclusion chromatography. Thyroglobulin (669 kDa), ferritin (440 kDa), aldolase (158 kDa), conalbumin (75 kDa), ovalbumin (43 kDa) were used as molecular markers. Data is average of two replicates **c-d**,The effect of metal ions (**c**) and pH (**d**) on activity. Data are mean ± SD of two independent assays. **e**, The temperature stability of the enzyme. Data are mean ± SD of three independent assays.

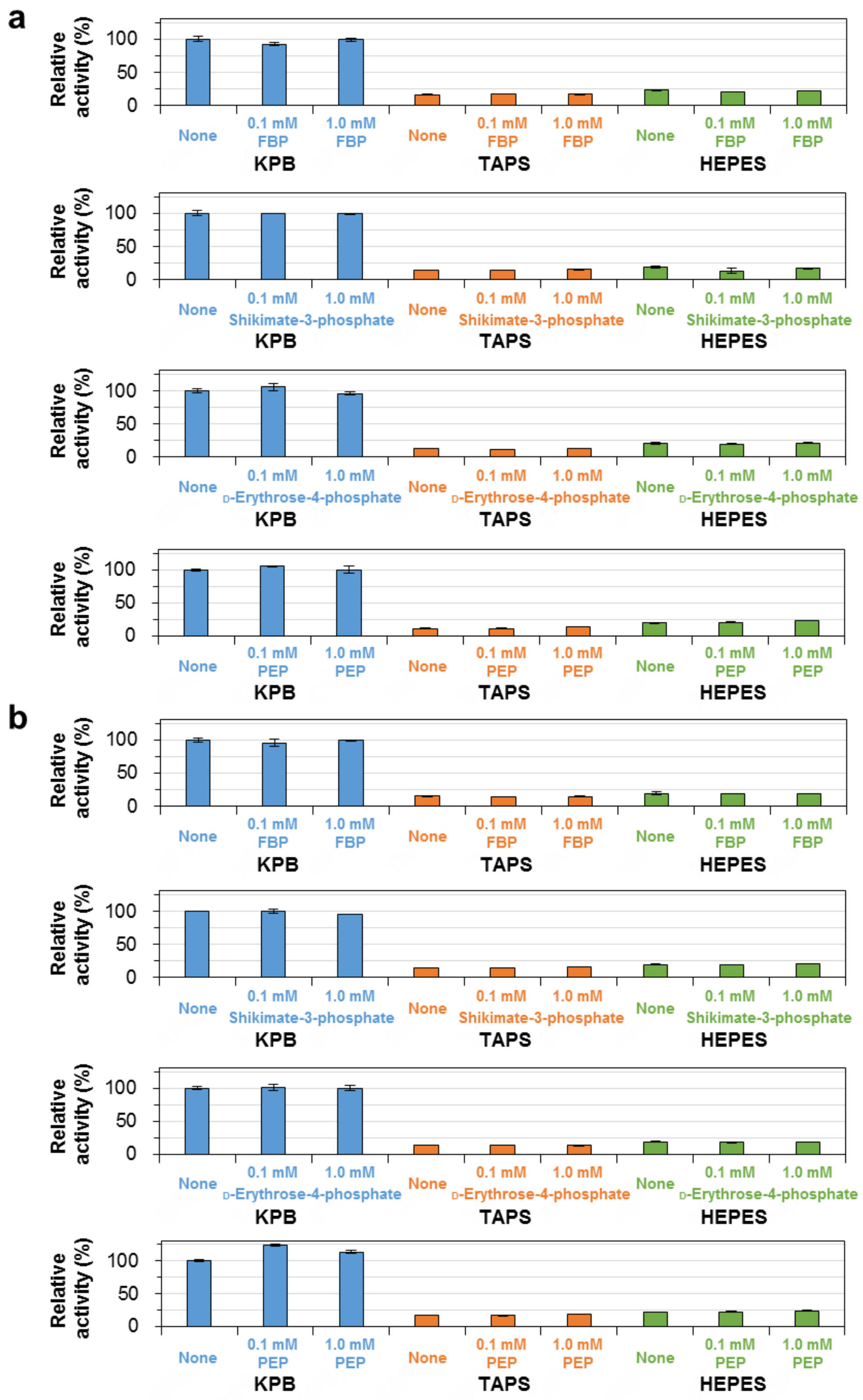


**Supplementary Figure 10. The effect of phosphate compounds on the activity of type 4 LDH enzyme.**
**a-b**, Phosphate compounds were added into the reaction mixture containing either 50 mM KPB, TAPS buffer, or HEPES buffer (pH 8.0) prior to the enzymatic assay. Substrate phenyl pyruvic acid was added to give a final concentration of 1 mM (**a**) and 4 mM (**b**). FBP, fructose-1,6-bisphosphate; PEP, phosphoenolpyruvate. All experiments were conducted in duplicates, and data are mean ± SD of two independent assays.

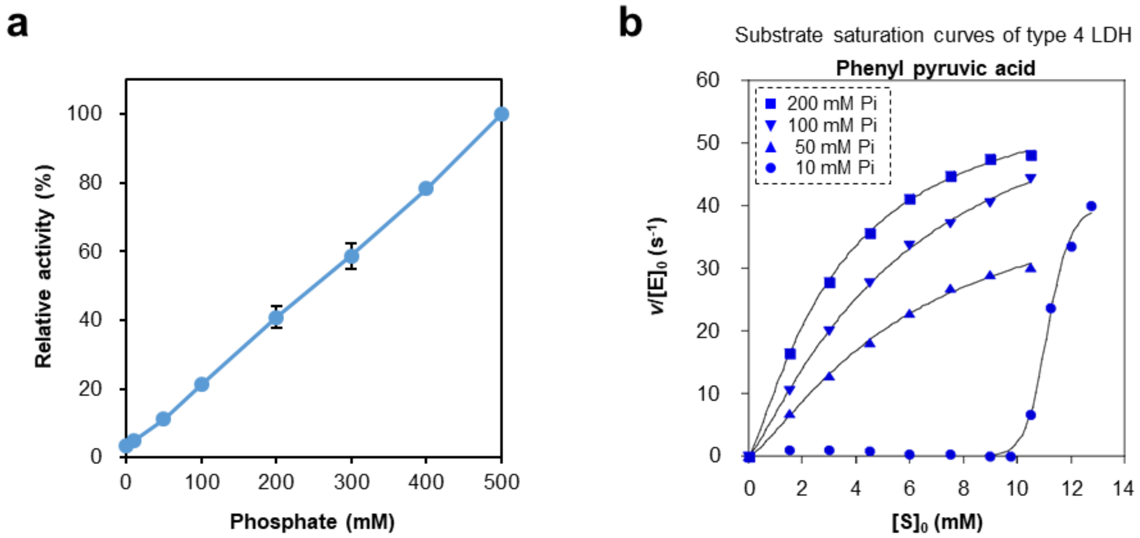


**c** Kinetic parameters (Hill equation) of type 4 LDH

| Substrate | Pi conc. (mM) | $K_{0.5}$ (mM) | $k_{cat}$ ($s^{-1}$) | $n_H$ |
|---|---|---|---|---|
| Phenyl pyruvic acid | 10 | 11.08 ± 0.08 | 39.62 ± 0.44 | 23.94 ± 3.69 |
| | 50 | 6.82 ± 1.18 | 51.08 ± 7.78 | 1.29 ± 0.03 |
| | 100 | 5.96 ± 0.06 | 64.22 ± 2.38 | 1.20 ± 0.01 |
| | 200 | 3.13 ± 0.29 | 55.45 ± 7.13 | 1.31 ± 0.07 |

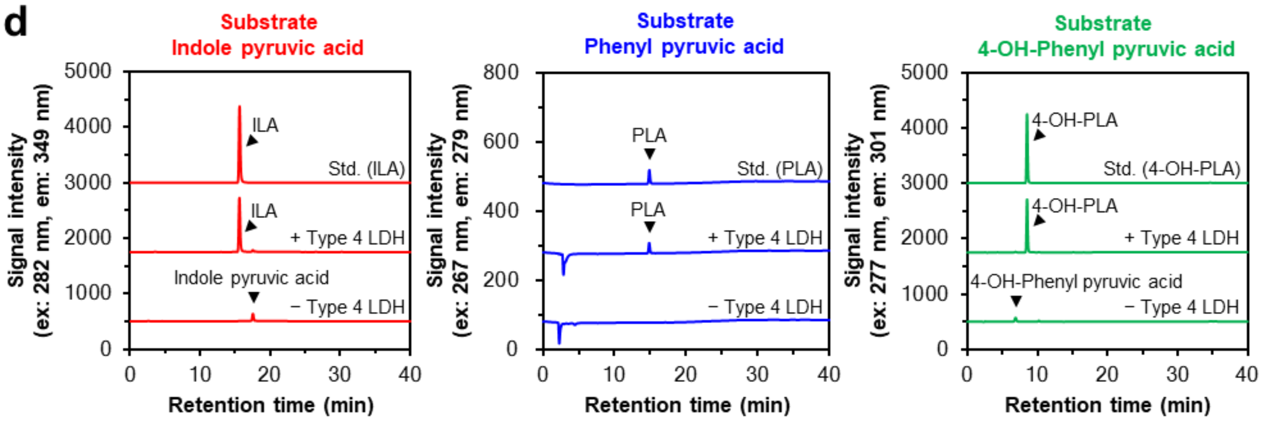


**Supplementary Figure 11. Effect of phosphate concentration on type 4 LDH activity and verification of aromatic lactic acid production.**

**a**, The effect of phosphate ion on enzyme activity. Data are mean ± SD of two independent assays. **b,c,** Substrate saturation curves (Hill equation) (**b**) and kinetic parameters (**c**) of type 4 LDH enzyme on the different concentrations of phosphate. The substrate saturation curves indicate the representative data of two independent assays. Kinetic parameters (Hill equation) are represented as mean ± SD of two independent assays. **d**, Verification of aromatic lactic acid production. The reaction mixture contained 100 mM KPB (pH 8.0), 1mM 2-ME, 1 mM β-NADH, 1.5 μM type 4 LDH enzyme, and 1 mM substrate, and the mixture was incubated at 37°C for 30 min. Production of ILA, PLA, and 4-OH-PLA was verified by HPLC. Only when indole pyruvic acid was used as a substrate, ten-fold diluted reaction mixture was applied to HPLC. Standard concentrations were 0.1 mM (ILA) or 1 mM (PLA and 4-OH-PLA). Note that phenyl pyruvic acid was not detected under the conditioned tested. Data are the representative of two independent assays.

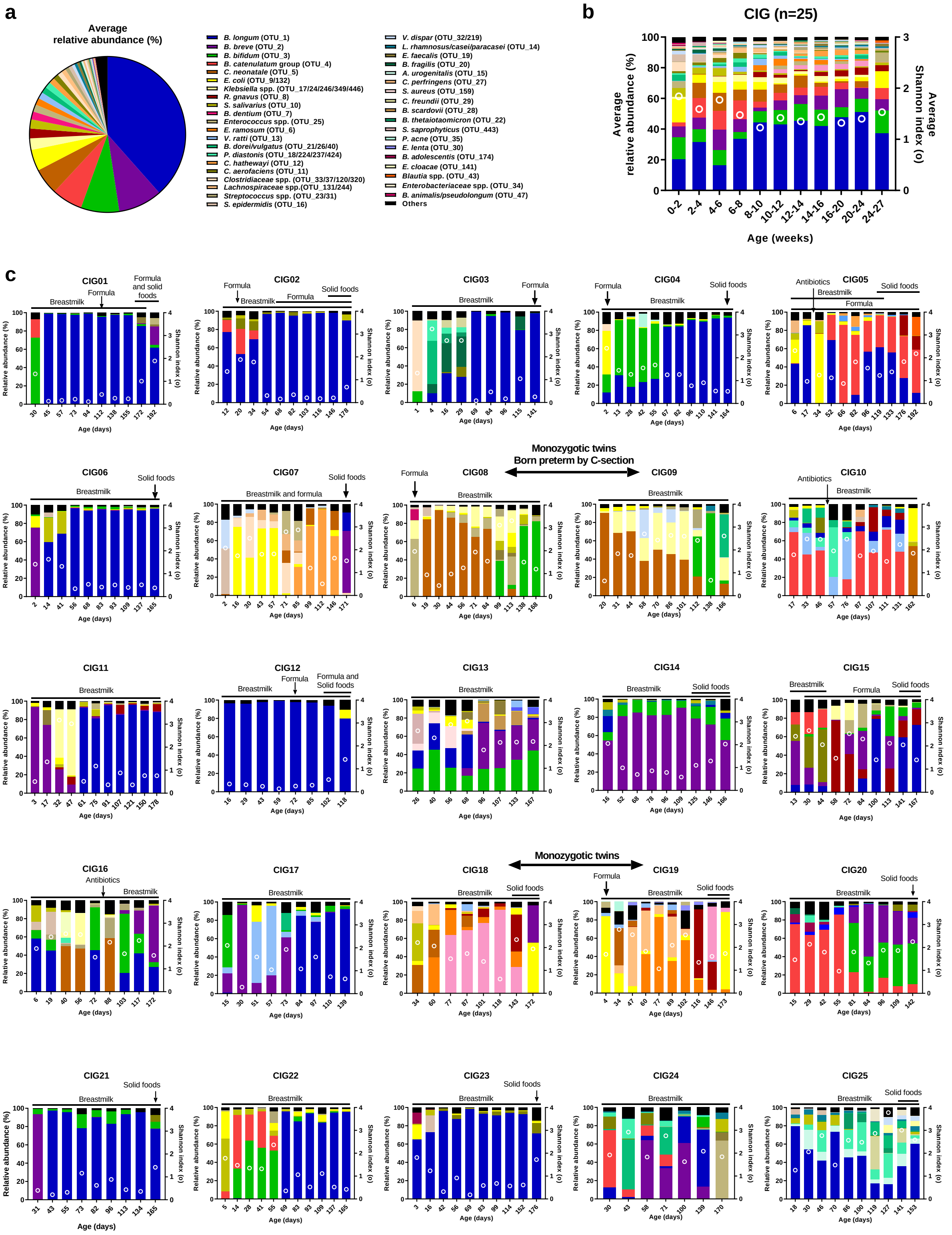


**Supplementary Figure 12. Gut microbiota composition of the CIG infants.**

**a**, Average relative abundance of the dominant gut microbial taxa (average relative abundance >0.1%) across all samples (comprising 97.5% of all microbial taxa detected). **b**, Temporal development of the average gut microbiota composition and Shannon diversity index (marked with circles) across all individuals. **c**, Intra-individual temporal development of gut microbiota composition and Shannon diversity index (marked with circles). Dietary patterns and consumption of antibiotics are indicated for each individual. If nothing else is indicated, infants were singletons, vaginally term born.

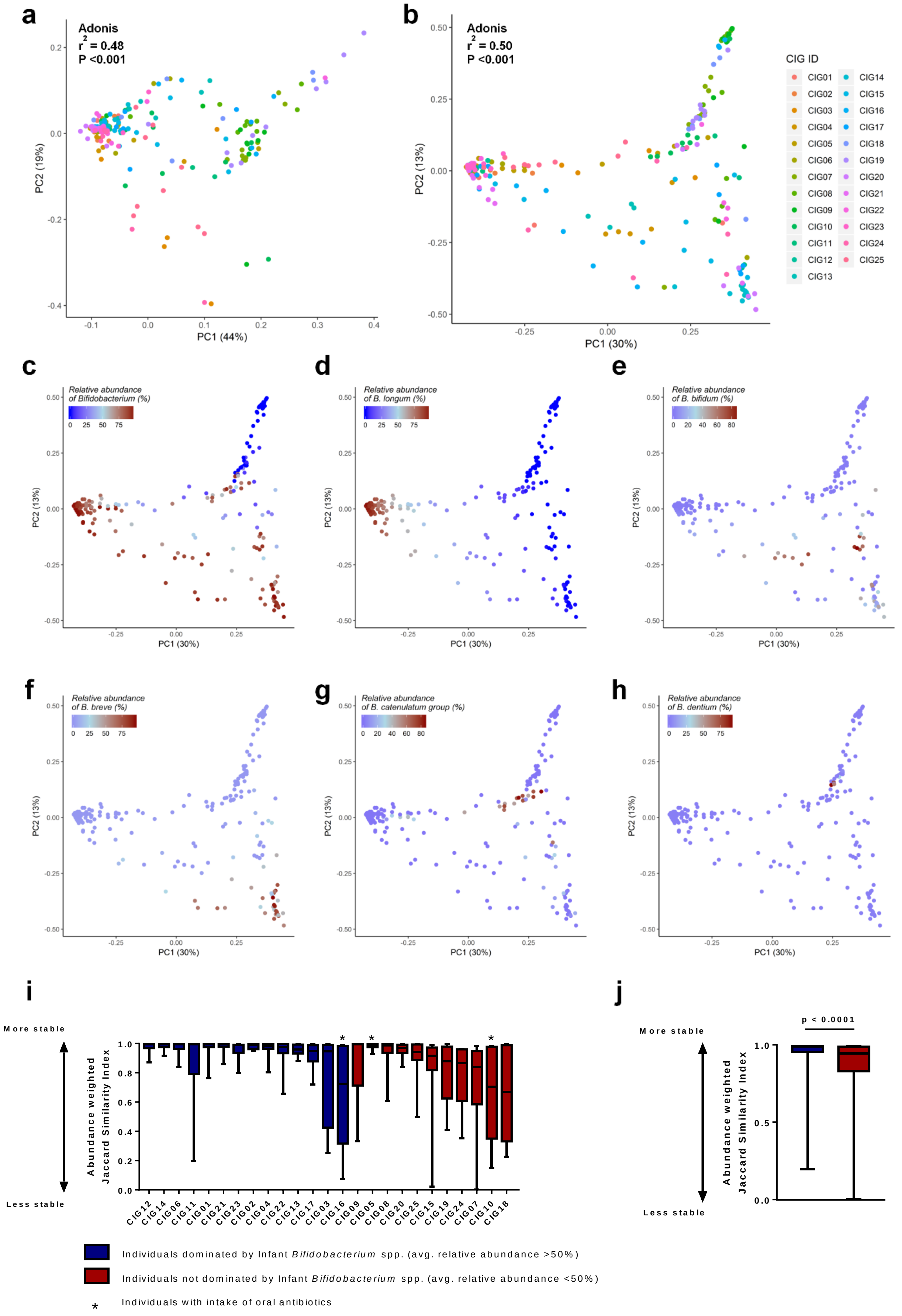
a
Adonis
r2 = 0.48
P <0.001
PC1 (44%)
PC2 (19%)
b
Adonis
r2 = 0.50
P <0.001
PC1 (30%)
PC2 (13%)
CIG ID
CIG01 CIG02 CIG03 CIG04 CIG05 CIG06 CIG07 CIG08 CIG09 CIG10 CIG11 CIG12 CIG13
CIG14 CIG15 CIG16 CIG17 CIG18 CIG19 CIG20 CIG21 CIG22 CIG23 CIG24 CIG25
c
Relative abundance of Bifidobacterium (%)
d
Relative abundance of B. longum (%)
e
Relative abundance of B. bifidum (%)
f
Relative abundance of B. breve (%)
g
Relative abundance of B. catenulatum group (%)
h
Relative abundance of B. dentium (%)
i
More stable
Less stable
Abundance weighted Jaccard Similarity Index
CIG12 CIG14 CIG06 CIG11 CIG01 CIG21 CIG23 CIG02 CIG04 CIG22 CIG13 CIG17 CIG03 CIG16 CIG09 CIG05 CIG08 CIG20 CIG25 CIG15 CIG19 CIG24 CIG07 CIG10 CIG18
j
More stable
Less stable
p < 0.0001
Abundance weighted Jaccard Similarity Index
Individuals dominated by Infant Bifidobacterium spp. (avg. relative abundance >50%)
Individuals not dominated by Infant Bifidobacterium spp. (avg. relative abundance <50%)
* Individuals with intake of oral antibiotics

**Supplementary Figure 13. Beta diversity in CIG cohort.**

**a-h**, Principal coordinates analysis (PCoA) plots of weighted UniFrac distances **(a)** or abundance Bray-Curtis dissimilarities **(b-h)**, based on all OTUs detected in CIG faecal samples (n=234), coloured according to **a-b,** CIG individual or relative abundances of **c,** *Bifidobacterium,* **d,** *Bifidobacterium longum,* **e**, *Bifidobacterium bifidum,* **f**, *Bifidobacterium breve,* **g**, *Bifidobacterium catenulatum* group or **h**, *Bifidobacterium dentium.* Statistical significance was evaluated by Adonis test. **i**, Boxplots of abundance weighted Jaccard similarity index (1- abundance weighted Jaccard distance). *Denotes that the individual at some point received oral antibiotics. **j**, Boxplots of abundance weighted Jaccard similarity index comparing CIG individuals dominated or not with infant type *Bifidobacterium* spp., excluding individuals receiving oral antibiotics. Statistical significance was evaluated by Mann-Whitney *U* test.

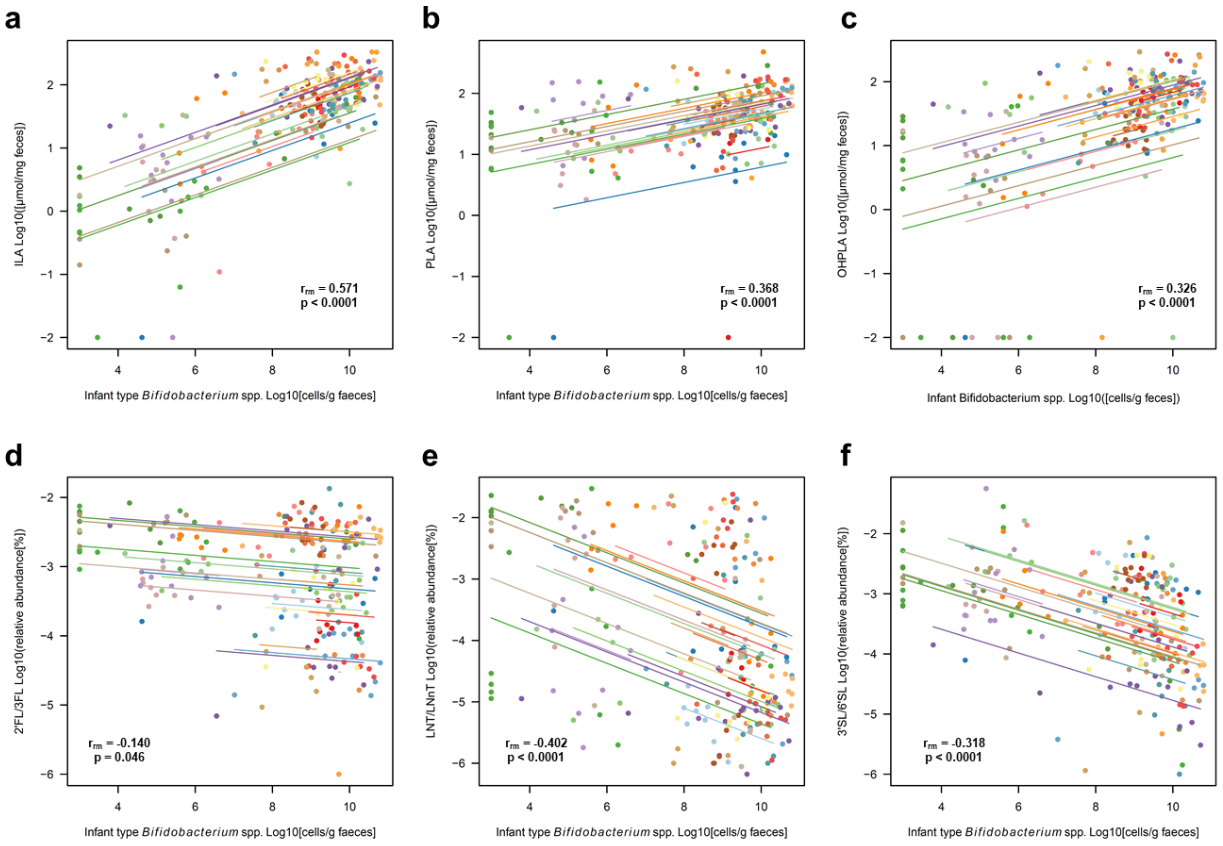


**Supplementary Figure 14. Associations between Infant type *Bifidobacterium* species and aromatic lactic acids and human milk oligosaccharides in faeces of infants from the CIG cohort.**

Repeated measures correlation plots of the relationship between faecal abundance of infant type *Bifidobacterium* spp, and faecal abundance of **a-c**, aromatic amino lactic acids (ILA, PLA and 4-OH-PLA) or **d-f**, HMOs (2´FL/3FL, LNT/LNnT and 3´SL/6'SL) in the CIG cohort. $r_{rm}$ is the repeated measures correlation coefficient. Each colour represent a CIG individual and parallel linear regression lines are shown for each individual.

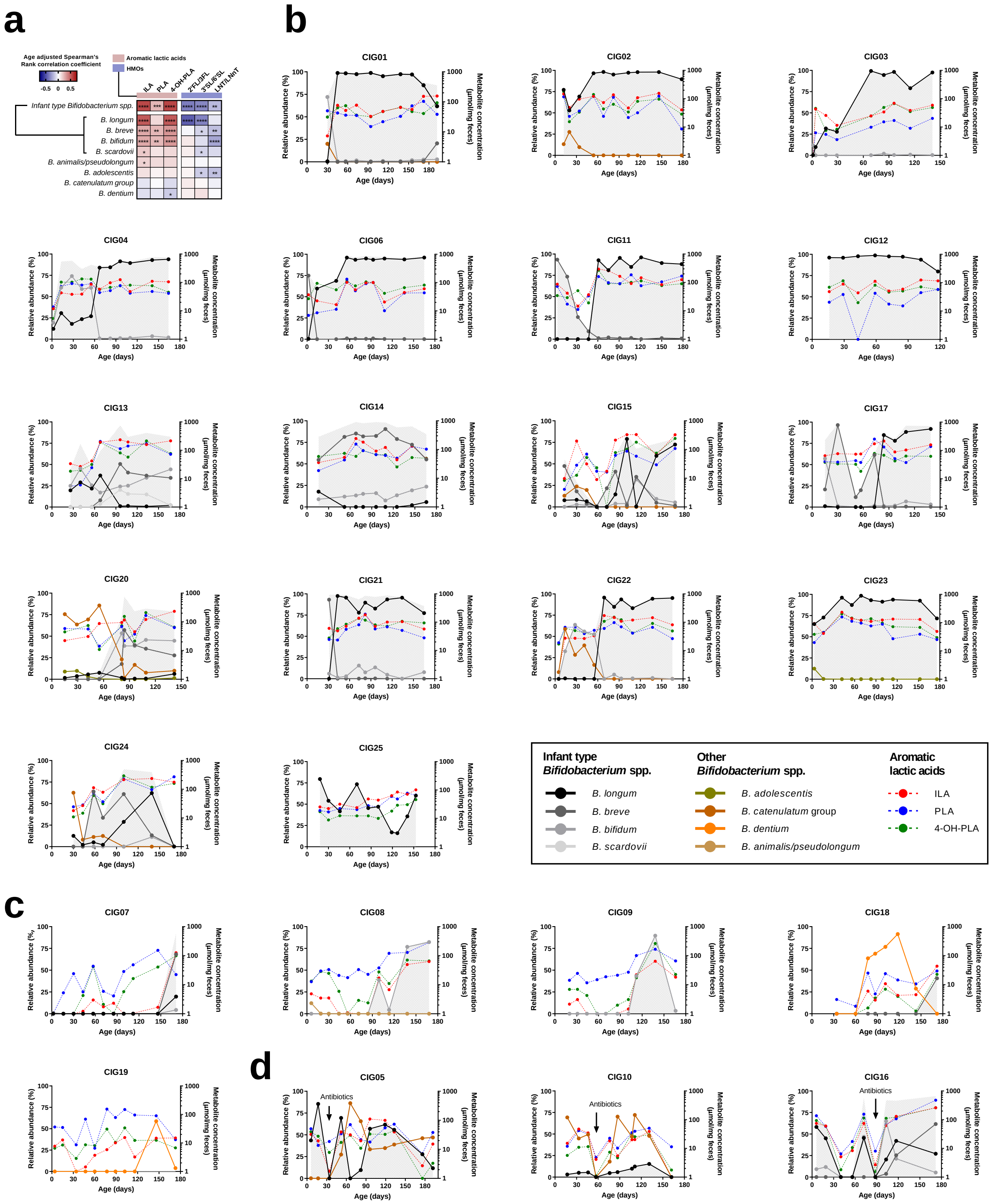


**Supplementary Figure 15. Relative abundances of *Bifidobacterium* species and concentrations of aromatic lactic acids in the CIG cohort.**

**a,** Heatmap illustrating partial Spearman's Rank correlation coefficients (adjusted for age) between the relative abundance of *Bifidobacterium* species and absolute faecal concentrations of aromatic lactic acids (n=240) or relative abundances of human milk oligosaccharides (n=228). Infant type *Bifidobacterium* spp. is the sum of the relative abundances of *B. longum*, *B. breve*, *B. bifidum* and *B. scardovii*. Statistical significance is indicated by asterisks with * p<0.05, **p<0.01, ***p<0.001 and ****p<0.0001. **b-d,** Relative abundance of *Bifidobacterium* spp. (average relative abundance >1% of total community) and concentrations of indolelactic acid (ILA), phenyllactic acid (PLA) and 4-hydroxyphenyllactic acid (4-OH-PLA) in all individuals from the Copenhagen Infant Gut (CIG) cohort. Values of metabolite concentrations below 1 µmol/mg feces are not shown. Summarized relative abundance of infant type *Bifidobactrium* spp. is indicated with grey background shading. **b**, Infants early and consistently colonised with infant type *Bifidobacterium* spp. (colonize within first month reaching average relative abundance >40% during first 6 months), **c,** Infants with late colonization of infant type *Bifidobacterium* spp. (not detectable or on average <0.5% of total community within the first 3 months of life) and **d**, Infants that received oral antibiotics during sampling.

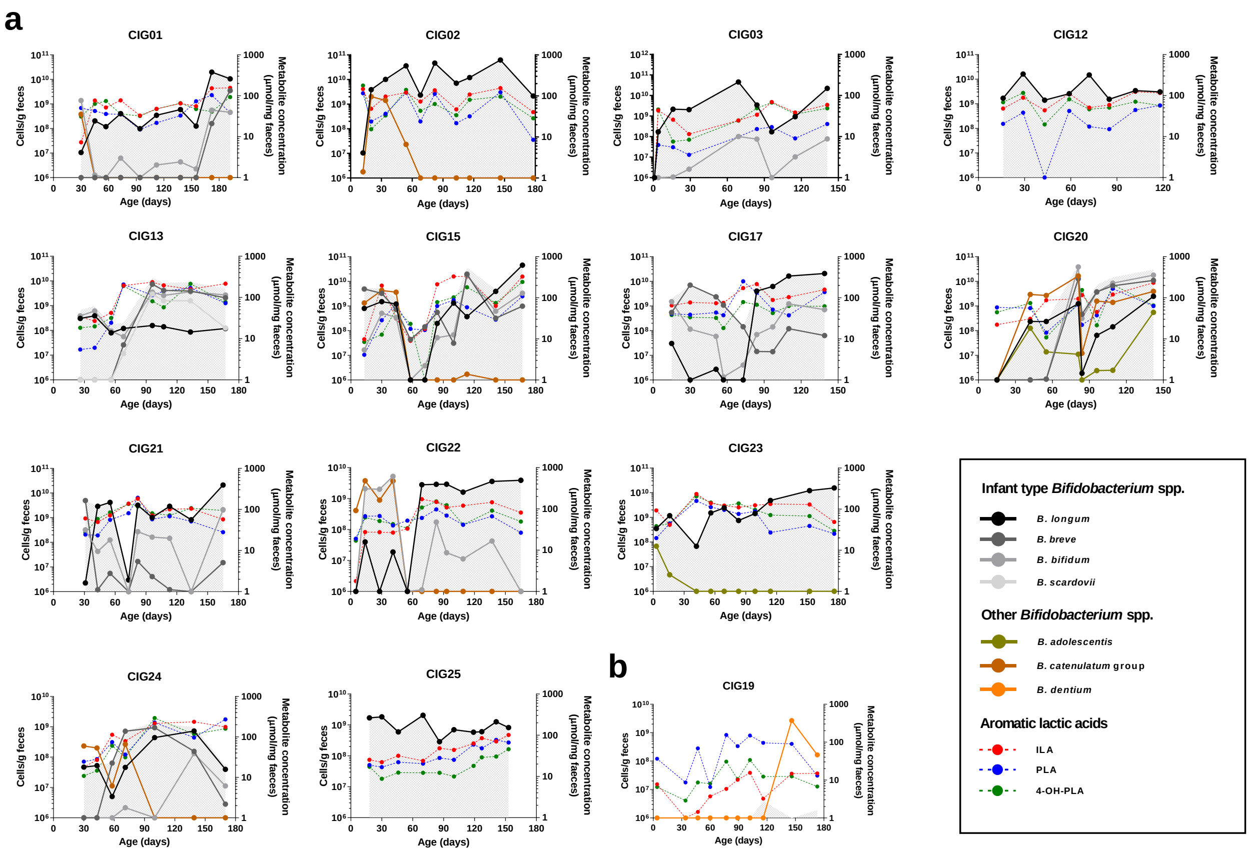


**Supplementary Figure 16. Absolute abundances of *Bifidobacterium* species and concentrations of aromatic lactic acids in the CIG cohort.**

**a**-**b**, Absolute abundance of *Bifidobacterium* spp. (average relative abundance >1% of total community) and concentrations of indolelactic acid (ILA), phenyllactic acid (PLA) and 4-hydroxyphenyllactic acid (4-OH-PLA) in individuals from the Copenhagen Infant Gut (CIG) cohort. Values of bacterial counts below $10^6$ cells/g feces and metabolite concentrations below 1 µmol/mg feces are not shown. Infant type *Bifidobacterium* spp. is the sum of the absolute abundances of *B. longum*, *B. breve*, *B. bifidum* and *B. scardovii* and is indicated with grey background shading. **a**, Infants early and consistently colonised with infant type *Bifidobacterium* spp. (colonize within first month reaching average relative abundance >40% during first 6 months), **b,** Infants with late colonization of infant type *Bifidobacterium* spp. (not detectable or on average <0.5% of total community within the first 3 months of life)

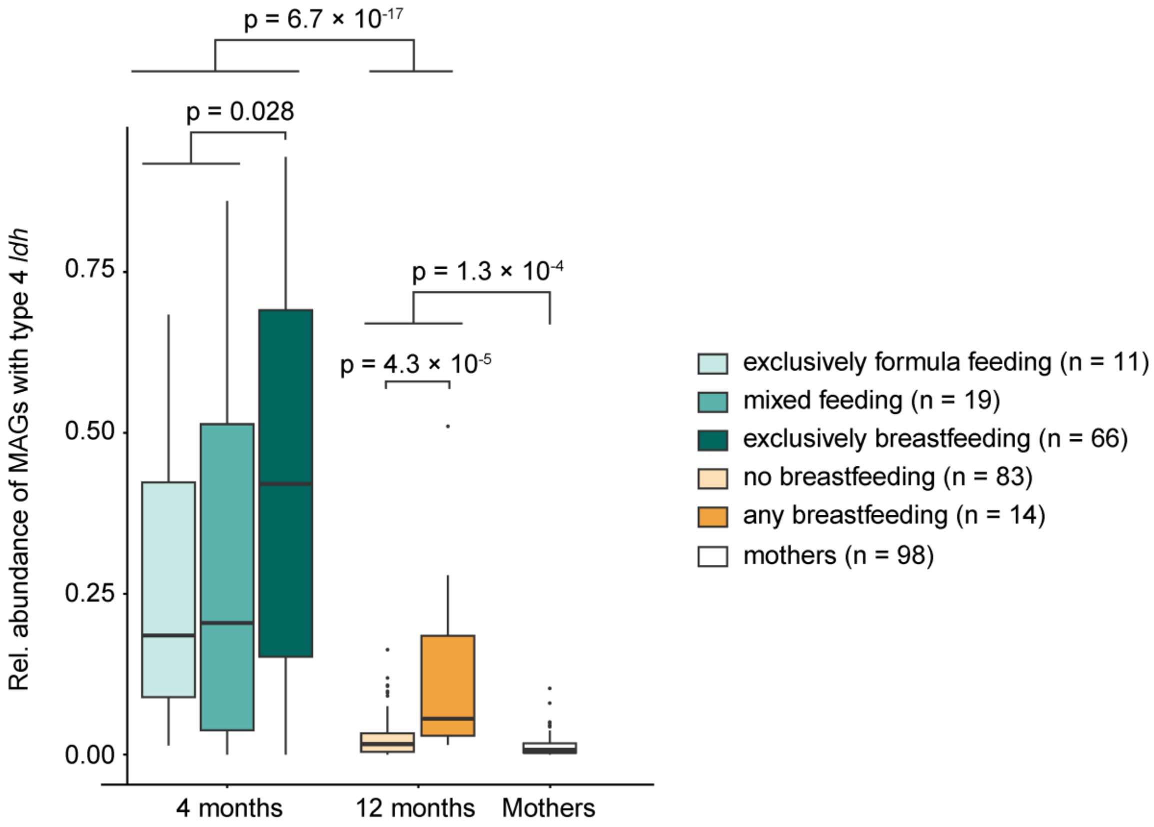


**Supplementary Figure 17. Relative abundance [fraction] of high quality metagenome-assembled genomes (MAGs) with *aldh* (type 4 *ldh*) in 4 months and 12 months old infants and mothers found by mining of publically available sequence data from Bäckhed *et al*[4].** Line, boxes and whiskers indicate median, IQR and ±1.5*IQR and statistical significance was evaluated by Mann Whitney U test.

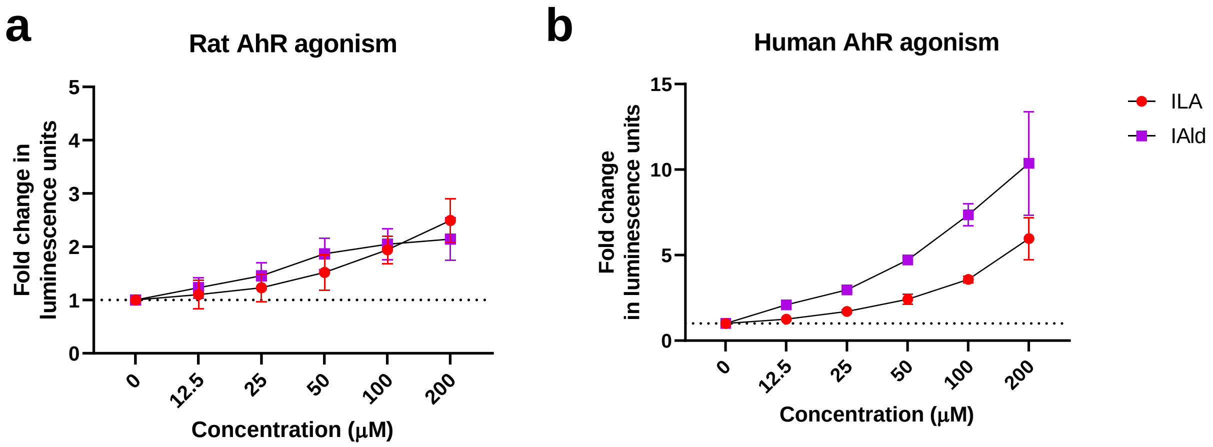


**Supplementary Figure 18. Indolelactic acid is an aryl hydrocarbon receptor (AhR) agonist.** Dose-dependent effects of indolelactic acid (ILA) and indolealdehyde (IAld) in **a,** rat and **b,** human AhR reporter assays. The AhR activity (Luminescence Units) of the indoles were reported as fold changes relative to vehicle signal (0 μM). Data are presented as mean+SD of two (**a**) to three (**b**) experiments including three technical replicates.

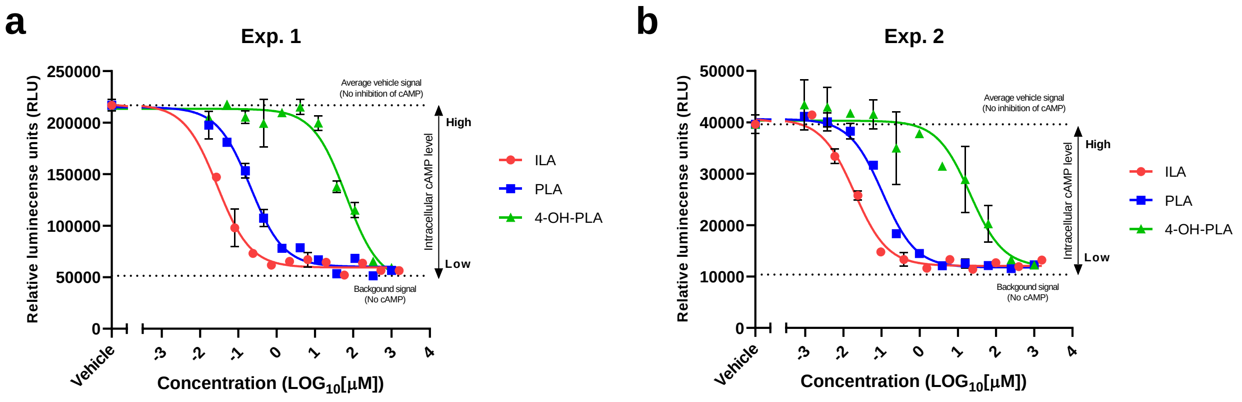


**Supplementary Figure 19. Aromatic lactic acids are hydroxycarboxylic acid 3 (HCAR3) receptor agonist.**

Dose-dependent effects of indolelactic acid (ILA), phenyllactic acid (PLA) and 4-hydroxy-phenyllactic acid (4-OH-PLA) in a human HCAR3 receptor assay (cAMP Hunter™ eXpress GPR109B CHO-K1 GPCR Assay). Activation of the G-protein coupled HCAR3 receptor results in activation of the Gα-inhibitory subunit, which inhibits intracellular adenylyl cyclase activity and reduces forskolin-induced cAMP levels. cAMP levels are detected by a competitive immunoassay based on β-galactosidase hydrolysis of a substrate producing a luminescence signal. The ability of the aromatic lactic acids to activate the HCAR3 receptor results in a decrease of forskolin-induced cAMP levels, resulting in inhibition of the luminescence signal (Relative Luminescence Units). Serial dilutions of the aromatic lactic acids were compared to average vehicle signal (0 µM) and average background signal (no forskolin induction of cAMP). Data are presented as two independent experiments (**a** and **b**) with mean±SD of two technical replicates each. Curves were fitted to the data points by non-linear regression.

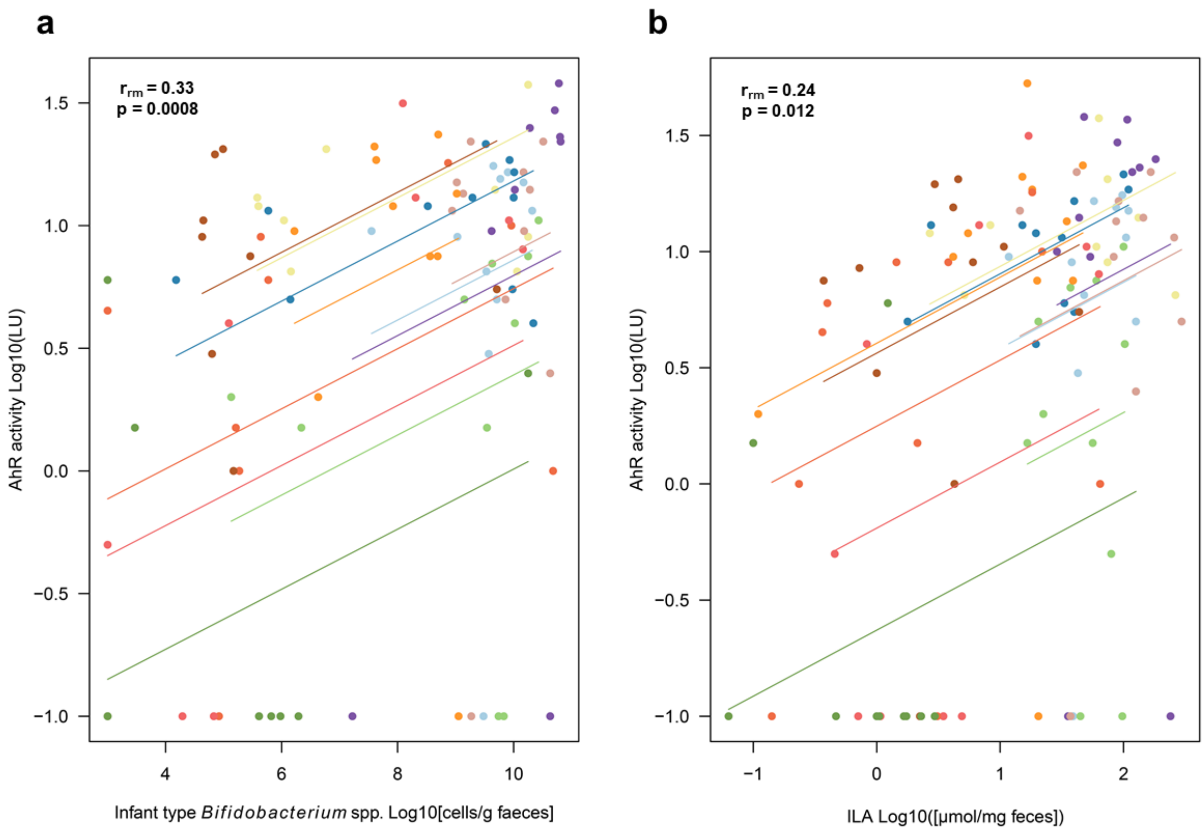


**Supplementary Figure 20. Associations between Infant type *Bifidobacterium* species/ILA and AhR activity in faecal water from the CIG infants.**

Repeated measures correlation plots of the relationship between faecal abundance of **a**, infant type *Bifidobacterium* spp or **b**, ILA and AhR activity (Luminescence Units, LU) measures in faecal water of the CIG infants. $r_{rm}$ is the repeated measures correlation coefficient. Each colour represent a CIG individual and parallel linear regression lines are shown for each individual.
