## Supplementary Data 1 for "Breastmilk-promoted bifidobacteria produce aromatic amino acids in the infant gut"

**Supplementary Data 1a. Metadata of the subset of SKOT cohort study participants included in this study (n=59)**

| Participant | Sample_ID | Birth mode# | Gender | Age (months) | Partial breastfeeding | Partial formulafeeding | Introduction to solid foods (month of age) | Medication within 14 days prior to sampling |
|---|---|---|---|---|---|---|---|---|
| 201008 | 1 | Vaginal | Female | 9.1 | No | No | 4 | No |
| 201010 | 2 | Vaginal | Male | 9.2 | Yes | Yes | 4 | No |
| 201011 | 3 | C-section | Female | 9.2 | No | Yes | 4 | No |
| 201012 | 4 | NA | Male | 9.4 | No | Yes | 4 | No |
| 201025 | 5 | Vaginal | Female | 8.8 | No | No | 6 | No |
| 201026 | 6 | Vaginal | Male | 8.7 | Yes | Yes | 4 | No |
| 201028 | 7 | Vaginal | Male | 8.8 | No | Yes | 4 | No |
| 201038 | 8 | Vaginal | Male | 9.4 | No | Yes | 4 | No |
| 201047 | 9 | Vaginal | Male | 9.0 | Yes | Yes | 4 | No |
| 201053 | 10 | Vaginal | Male | 9.2 | No | Yes | 4 | No |
| 201061 | 11 | Vaginal | Male | 9.3 | Yes | Yes | 4 | No |
| 201067 | 12 | Vaginal | Male | 8.9 | Yes | No | 5 | No |
| 201078 | 13 | Vaginal | Female | 9.2 | Yes | No | 4 | No |
| 201080 | 14 | C-section | Female | 9.4 | No | Yes | 4 | No |
| 201081 | 15 | Vaginal | Female | 9.4 | No | No | 6 | No |
| 201085 | 16 | Vaginal | Female | 9.1 | Yes | No | 5 | No |
| 201093 | 17 | Vaginal | Female | 9.1 | Yes | No | 5 | No |
| 201103 | 18 | Vaginal | Male | 9.3 | No | Yes | 4 | No |
| 201117 | 19 | Vaginal | Male | 8.9 | No | Yes | 5 | No |
| 201120 | 20 | Vaginal | Male | 8.8 | No | Yes | 4 | No |
| 201123 | 21 | Vaginal | Female | 8.6 | No | Yes | 4 | No |
| 201127 | 22 | Vaginal | Male | 8.6 | No | Yes | 6 | No |
| 201137 | 23 | Vaginal | Female | 9.3 | Yes | Yes | 5 | Oral penicillin ended one week prior to sample collection |
| 201139 | 24 | C-section | Female | 8.6 | No | Yes | 5 | No |
| 201149 | 25 | Vaginal | Female | 9.1 | No | Yes | 3 | No |
| 201157 | 26 | Vaginal | Male | 9.1 | No | Yes | 4 | No |
| 201158 | 27 | Vaginal | Male | 9.1 | Yes | No | 4 | Salbuvent/Salbutamol against asthma |
| 201159 | 28 | Vaginal | Female | 9.2 | No | Yes | 4 | No |
| 201164 | 29 | Vaginal | Female | 8.7 | Yes | Yes | 4 | No |
| 201168 | 30 | Vaginal | Female | 8.8 | Yes | Yes | 4 | No |
| 201174 | 31 | Vaginal | Female | 9.0 | No | Yes | 4 | No |
| 201183 | 32 | C-section | Male | 8.8 | No | Yes | 4 | No |
| 201193 | 33 | Vaginal | Female | 9.7 | No | Yes | 4 | No |
| 201203 | 34 | Vaginal | Female | 9.2 | Yes | No | 4 | No |
| 201211 | 35 | Vaginal | Female | 9.5 | No | No | 6 | No |
| 201213 | 36 | Vaginal | Male | 9.1 | No | Yes | 4 | No |
| 201222 | 37 | Vaginal | Male | 8.9 | Yes | No | 5 | No |
| 201225 | 38 | C-section | Female | 8.8 | No | Yes | 4 | No |
| 201227 | 39 | Vaginal | Female | 8.7 | No | Yes | 4 | No |
| 201233 | 40 | Vaginal | Male | 9.1 | No | Yes | 6 | No |
| 201236 | 41 | Vaginal | Female | 9.3 | Yes | Yes | 4 | No |
| 201248 | 42 | Vaginal | Male | 9.5 | No | Yes | 5 | No |
| 201251 | 43 | Vaginal | Male | 9.5 | Yes | No | 5 | No |
| 201257 | 44 | Vaginal | Male | 9.4 | Yes | Yes | 5 | No |
| 201261 | 45 | C-section | Male | 9.4 | No | Yes | 4 | No |
| 201276 | 46 | Vaginal | Male | 9.3 | Yes | No | 4 | No |
| 201278 | 47 | Vaginal | Male | 9.4 | No | Yes | 3 | No |
| 201285 | 48 | Vaginal | Female | 9.4 | No | Yes | 4 | No |
| 201286 | 49 | Vaginal | Female | 9.2 | Yes | Yes | 6 | No |
| 201291 | 50 | Vaginal | Female | 9.2 | No | Yes | 4 | No |
| 201297 | 51 | Vaginal | Male | 8.9 | Yes | Yes | 6 | No |
| 201311 | 52 | C-section | Female | 9.1 | No | Yes | 4 | No |
| 201312 | 53 | Vaginal | Female | 8.8 | Yes | Yes | 5 | No |
| 201313 | 54 | Vaginal | Female | 9.0 | No | Yes | 4 | No |
| 201316 | 55 | Vaginal | Male | 8.9 | Yes | No | 5 | No |
| 201318 | 56 | Vaginal | Female | 9.1 | Yes | Yes | 4 | No |
| 201319 | 57 | Vaginal | Male | 9.1 | No | Yes | 4 | No |
| 201321 | 58 | Vaginal | Male | 9.1 | Yes | Yes | 5 | No |
| 201322 | 59 | Vaginal | Male | 9.1 | No | No | 5 | No |

#Data on mode of delivery from one individual was missing

**Supplementary Data 1b. Cohort characteristics of the subset of SKOT cohort study participants included in this study (n=59)**

| Parameter | SKOT (n = 59) |
|---|---|
| Age (average months ± SD) | 9.1 ± 0.3 |
| Sex | |
| *Male* | 30 (50.8%) |
| *Female* | 29 (49.2%) |
| Mode of delivery[#] | |
| *Vaginal* | 51 (87.9%) |
| *C-section* | 7 (12.1%) |
| Diet | |
| *Duration of exclusive breastfeeding (average months ± SD)* | 3.8 ± 1.8 |
| *Introduction to solid food (average months ± SD)* | 4.4 ± 0.7 |
| *Total duration of breastfeeding (average months ± SD)* | 8.6 ± 3.9 |
| *Partially breastfed at 9 months of age* | 24 (40.7%) |

[#]Data on mode of delivery from one individual was missing

**Supplementary Data 1c. Adonis and permdisp test of beta diversity distance matrices according to metadata variables in the SKOT cohort study (n=59)**

| **Weighted UniFrac** | **adonis** | | | **permdisp[1]** | |
|---|---|---|---|---|---|
| **Parameter** | **F-test** | **$r^2$** | **permuted p-value** | **F-test** | **permuted p-value** |
| Mode of delivery | 2.217 | 0.038 | 0.076 | - | - |
| Age | 0.620 | 0.011 | 0.657 | - | - |
| Gender | 0.430 | 0.008 | 0.841 | - | - |
| Partial breastfeeding (vs. no breastfeeding) | 5.840 | 0.093 | **0.001** | 0.040 | 0.850 |
| Partial formulafeeding (vs. no formulafeeding) | 2.780 | 0.046 | **0.032** | 0.010 | 0.910 |
| Age at introduction to solid foods | 0.823 | 0.014 | 0.500 | - | - |
| **Unweighted UniFrac** | **adonis** | | | **permdisp[1]** | |
| **Parameter** | **F-test** | **$r^2$** | **permuted p-value** | **F-test** | **permuted p-value** |
| Mode of delivery | 1.918 | 0.033 | **0.021** | 0.101 | 0.762 |
| Age | 1.213 | 0.021 | 0.230 | - | - |
| Gender | 1.305 | 0.022 | 0.181 | - | - |
| Partial breastfeeding (vs. no breastfeeding) | 1.793 | 0.031 | **0.035** | 4.770 | **0.025** |
| Partial formulafeeding (vs. no formulafeeding) | 1.051 | 0.018 | 0.371 | - | - |
| Age at introduction to solid foods | 0.970 | 0.017 | 0.471 | - | - |
| **Abundance Bray Curtis** | **adonis** | | | **permdisp[1]** | |
| **Parameter** | **F-test** | **$r^2$** | **permuted p-value** | **F-test** | **permuted p-value** |
| Mode of delivery | 1.587 | 0.028 | 0.082 | - | - |
| Age | 0.782 | 0.014 | 0.693 | - | - |
| Gender | 1.158 | 0.020 | 0.312 | - | - |
| Partial breastfeeding (vs. no breastfeeding) | 2.650 | 0.044 | **0.008** | 0.087 | 0.780 |
| Partial formulafeeding (vs. no formulafeeding) | 2.037 | 0.035 | **0.021** | 3.415 | 0.081 |
| Age at introduction to solid foods | 0.895 | 0.015 | 0.564 | - | - |
| **Binary Bray Curtis** | **adonis** | | | **permdisp[1]** | |
| **Parameter** | **F-test** | **$r^2$** | **permuted p-value** | **F-test** | **permuted p-value** |
| Mode of delivery | 1.275 | 0.022 | 0.170 | - | - |
| Age | 1.270 | 0.022 | 0.210 | - | - |
| Gender | 1.181 | 0.020 | 0.242 | - | - |
| Partial breastfeeding (vs. no breastfeeding) | 2.440 | 0.041 | **0.001** | 3.032 | 0.072 |
| Partial formulafeeding (vs. no formulafeeding) | 1.348 | 0.023 | 0.138 | - | - |
| Age at introduction to solid foods | 0.980 | 0.017 | 0.469 | - | - |

[1]permdisp tests were only computed for variables that showed statistical significance (permuted p-value<0.05) in the adonis test

**Supplementary Data 1d. Effects of breastfeeding, formulafeeding and mode of delivery on genus abundances in the SKOT cohort study (n=59)**

| | Breastfeeding | | | | | Formula | | | | | Mode of delivery | | | | |
|---|---|---|---|---|---|---|---|---|---|---|---|---|---|---|---|
| **Genus** | **Test-Statistic[#]** | **p-value** | **FDR p-value[$]** | **Partially breastfed (Mean realtive abundance)** | **Weaned (Mean relative abundance)** | **Test-Statistic[#]** | **p-value** | **FDR p-value[$]** | **Partially formulafed (mean relative abundance)** | **No formula (mean realtive abundance)** | **Test-Statistic[#]** | **p-value** | **FDR p-value[$]** | **C-section (mean relative abundance)** | **Vaginal (mean relative abundance)** |
| g__Bifidobacterium | 204 | **0.001** | **0.044** | 49.578 | 25.018 | 213 | **0.043** | 0.603 | 31.231 | 46.087 | 95 | **0.048** | 0.476 | 19.574 | 37.318 |
| g__[Ruminococcus] | 251 | **0.009** | 0.215 | 0.040 | 0.081 | 280 | 0.389 | 0.934 | 0.075 | 0.035 | 105 | 0.081 | 0.678 | 0.166 | 0.051 |
| f__UnclassifiedLachnospiraceae | 265 | **0.017** | 0.215 | 20.238 | 30.889 | 275 | 0.343 | 0.934 | 29.027 | 19.308 | 127 | 0.223 | 0.903 | 40.412 | 24.982 |
| g__Lactobacillus | 280 | **0.024** | 0.215 | 1.040 | 0.346 | 236 | 0.087 | 0.603 | 0.325 | 1.517 | 156 | 0.582 | 0.903 | 0.190 | 0.701 |
| g__Pseudobutyrivibrio | 276 | **0.025** | 0.215 | 3.177 | 0.896 | 303 | 0.641 | 0.934 | 2.023 | 1.238 | 166 | 0.763 | 0.903 | 1.142 | 1.953 |
| g__Lachnospira | 283 | **0.026** | 0.215 | 0.619 | 1.208 | 297 | 0.543 | 0.934 | 1.003 | 0.867 | 175 | 0.940 | 0.959 | 0.497 | 1.052 |
| g__Moryella | 315 | **0.039** | 0.263 | 0.002 | 0.007 | 328 | 0.973 | 0.984 | 0.005 | 0.005 | 171 | 0.820 | 0.903 | 0.007 | 0.005 |
| g__Blautia | 292 | **0.047** | 0.263 | 5.086 | 12.613 | 219 | 0.052 | 0.603 | 11.642 | 3.418 | 156 | 0.598 | 0.903 | 12.021 | 9.400 |
| g__Roseburia | 296 | **0.049** | 0.263 | 0.146 | 0.835 | 324 | 0.921 | 0.984 | 0.266 | 1.401 | 144 | 0.404 | 0.903 | 0.430 | 0.583 |
| g__Megasphaera | 309 | 0.069 | 0.263 | 0.105 | 0.102 | 251 | 0.146 | 0.669 | 0.110 | 0.085 | 96 | **0.038** | 0.472 | 0.002 | 0.120 |
| g__Actinomyces | 303 | 0.072 | 0.263 | 0.062 | 0.028 | 265 | 0.261 | 0.869 | 0.035 | 0.063 | 174 | 0.924 | 0.959 | 0.034 | 0.044 |
| g__Coprococcus | 311 | 0.076 | 0.263 | 0.060 | 0.217 | 325 | 0.927 | 0.984 | 0.170 | 0.103 | 159 | 0.632 | 0.903 | 0.272 | 0.140 |
| g__Epulopiscium | 356 | 0.079 | 0.263 | 0.062 | 0.016 | 305 | 0.448 | 0.934 | 0.046 | 0.001 | 172 | 0.800 | 0.903 | 0.076 | 0.030 |
| g__Prevotella | 343 | 0.080 | 0.263 | 0.667 | 0.164 | 323 | 0.868 | 0.984 | 0.195 | 0.879 | 140 | 0.185 | 0.903 | 0.000 | 0.426 |
| g__Parabacteroides | 325 | 0.084 | 0.263 | 0.175 | 0.670 | 304 | 0.599 | 0.934 | 0.501 | 0.375 | 128 | 0.159 | 0.903 | 0.000 | 0.542 |
| g__[Eubacterium] | 311 | 0.084 | 0.263 | 0.060 | 0.275 | 251 | 0.161 | 0.669 | 0.211 | 0.120 | 161 | 0.676 | 0.903 | 0.133 | 0.188 |
| g__Enterococcus | 312 | 0.097 | 0.284 | 1.474 | 1.077 | 243 | 0.131 | 0.669 | 1.023 | 1.870 | 153 | 0.550 | 0.903 | 3.753 | 0.830 |
| g__Bacteroides | 318 | 0.114 | 0.317 | 2.615 | 7.180 | 327 | 0.958 | 0.984 | 5.707 | 4.198 | 153 | 0.549 | 0.903 | 1.994 | 4.776 |
| g__Oscillospira | 339 | 0.128 | 0.336 | 0.005 | 0.025 | 314 | 0.741 | 0.934 | 0.019 | 0.011 | 160 | 0.600 | 0.903 | 0.017 | 0.017 |
| g__Turicibacter | 350 | 0.161 | 0.380 | 0.002 | 0.011 | 312 | 0.690 | 0.934 | 0.007 | 0.008 | 109 | **0.027** | 0.452 | 0.027 | 0.005 |
| g__Klebsiella | 345 | 0.163 | 0.380 | 0.024 | 0.113 | 300 | 0.529 | 0.934 | 0.091 | 0.036 | 146 | 0.360 | 0.903 | 0.056 | 0.081 |
| g__Veillonella | 330 | 0.167 | 0.380 | 1.592 | 1.750 | 234 | 0.096 | 0.603 | 1.229 | 3.026 | 168 | 0.811 | 0.903 | 0.562 | 1.852 |
| g__Atopobium | 356 | 0.200 | 0.423 | 0.012 | 0.004 | 328 | 0.964 | 0.984 | 0.007 | 0.005 | 150 | 0.385 | 0.903 | 0.001 | 0.008 |
| g__Clostridium | 337 | 0.203 | 0.423 | 1.372 | 3.116 | 224 | 0.064 | 0.603 | 1.220 | 0.124 | 162 | 0.703 | 0.903 | 3.501 | 2.297 |
| g__Sedimentibacter | 403 | 0.241 | 0.476 | 0.006 | 0.000 | 323 | 0.586 | 0.934 | 0.004 | 0.000 | 175 | 0.751 | 0.903 | 0.000 | 0.003 |
| g__Slackia | 396 | 0.247 | 0.476 | 0.000 | 0.032 | 315 | 0.421 | 0.934 | 0.025 | 0.000 | 172 | 0.624 | 0.903 | 0.000 | 0.022 |
| g__Fusobacterium | 358 | 0.301 | 0.511 | 0.097 | 0.016 | 273 | 0.283 | 0.886 | 0.056 | 0.026 | 168 | 0.795 | 0.903 | 0.007 | 0.055 |
| g__SMB53 | 353 | 0.301 | 0.511 | 1.325 | 2.194 | 286 | 0.449 | 0.934 | 1.744 | 2.124 | 131 | 0.262 | 0.903 | 3.100 | 1.696 |
| g__Dorea | 355 | 0.306 | 0.511 | 0.324 | 1.077 | 225 | 0.064 | 0.603 | 0.985 | 0.142 | 78 | **0.015** | 0.426 | 3.882 | 0.359 |
| g__Butyricicoccus | 388 | 0.307 | 0.511 | 0.000 | 0.041 | 324 | 0.843 | 0.984 | 0.020 | 0.038 | 167 | 0.590 | 0.903 | 0.043 | 0.022 |
| Others | 356 | 0.327 | 0.523 | 1.310 | 1.178 | 284 | 0.428 | 0.934 | 1.156 | 1.454 | 170 | 0.849 | 0.903 | 0.718 | 1.323 |
| g__Streptococcus | 357 | 0.334 | 0.523 | 0.171 | 0.129 | 304 | 0.657 | 0.934 | 0.150 | 0.135 | 169 | 0.830 | 0.903 | 0.159 | 0.146 |
| g__Collinsella | 368 | 0.349 | 0.528 | 2.902 | 3.637 | 308 | 0.662 | 0.934 | 3.297 | 3.458 | 169 | 0.803 | 0.903 | 2.579 | 3.507 |
| g__Eggerthella | 372 | 0.377 | 0.554 | 0.012 | 0.011 | 274 | 0.244 | 0.869 | 0.011 | 0.013 | 152 | 0.449 | 0.903 | 0.015 | 0.011 |
| g__Escherichia | 370 | 0.431 | 0.616 | 0.117 | 0.065 | 326 | 0.943 | 0.984 | 0.071 | 0.131 | 167 | 0.789 | 0.903 | 0.080 | 0.086 |
| g__Dialister | 398 | 0.555 | 0.756 | 0.073 | 0.035 | 307 | 0.486 | 0.934 | 0.067 | 0.001 | 148 | 0.206 | 0.903 | 0.002 | 0.058 |
| g__Trabulsiella | 403 | 0.559 | 0.756 | 0.031 | 0.005 | 329 | 0.984 | 0.984 | 0.004 | 0.050 | 165 | 0.463 | 0.903 | 0.000 | 0.018 |
| g__Clostridium | 384 | 0.581 | 0.764 | 0.233 | 1.427 | 311 | 0.747 | 0.934 | 2.428 | 2.345 | 167 | 0.791 | 0.903 | 1.922 | 0.808 |
| g__Granulicatella | 386 | 0.598 | 0.767 | 0.035 | 0.026 | 209 | **0.034** | 0.603 | 0.036 | 0.011 | 79 | **0.017** | 0.426 | 0.057 | 0.026 |
| g__Citrobacter | 399 | 0.716 | 0.872 | 0.023 | 0.037 | 304 | 0.609 | 0.934 | 0.036 | 0.016 | 167 | 0.760 | 0.903 | 0.019 | 0.032 |
| g__Lactococcus | 402 | 0.730 | 0.872 | 0.013 | 0.006 | 314 | 0.730 | 0.934 | 0.011 | 0.002 | 177 | 0.964 | 0.964 | 0.008 | 0.009 |
| g__Haemophilus | 399 | 0.732 | 0.872 | 0.055 | 0.039 | 310 | 0.714 | 0.934 | 0.051 | 0.030 | 166 | 0.756 | 0.903 | 0.056 | 0.044 |
| g__Sutterella | 408 | 0.794 | 0.886 | 0.034 | 0.033 | 311 | 0.626 | 0.934 | 0.034 | 0.031 | 140 | 0.185 | 0.903 | 0.000 | 0.039 |
| g__Faecalibacterium | 406 | 0.801 | 0.886 | 4.523 | 2.459 | 260 | 0.158 | 0.669 | 2.872 | 4.550 | 155 | 0.523 | 0.903 | 0.579 | 3.737 |
| g__Dysgonomonas | 414 | 0.808 | 0.886 | 0.002 | 0.016 | 293 | 0.091 | 0.603 | 0.000 | 0.040 | 161 | 0.290 | 0.903 | 0.001 | 0.012 |
| g__Peptostreptococcus | 406 | 0.815 | 0.886 | 0.051 | 0.007 | 319 | 0.837 | 0.984 | 0.007 | 0.076 | 147 | 0.400 | 0.903 | 0.006 | 0.028 |
| g__Ruminococcus | 409 | 0.852 | 0.906 | 0.434 | 0.756 | 306 | 0.637 | 0.934 | 0.657 | 0.532 | 154 | 0.511 | 0.903 | 1.878 | 0.466 |
| g__Megamonas | 416 | 0.911 | 0.949 | 0.001 | 0.015 | 311 | 0.494 | 0.934 | 0.010 | 0.006 | 161 | 0.405 | 0.903 | 0.000 | 0.011 |
| g__Phascolarctobacterium | 417 | 0.936 | 0.955 | 0.004 | 0.017 | 293 | 0.182 | 0.702 | 0.016 | 0.000 | 161 | 0.405 | 0.903 | 0.000 | 0.014 |
| g__Enterobacter | 417 | 0.959 | 0.959 | 0.010 | 0.100 | 312 | 0.683 | 0.934 | 0.082 | 0.009 | 172 | 0.836 | 0.903 | 0.022 | 0.070 |

[#]Mann Whitney test

**Supplementary Data 1e. Correlations between relative abundance of gut microbiota genera and faecal concentrations of aromatic lactic acids and indole aldehyde in the SKOT cohort study (n=59)**

| GreenGenes Taxonomy | Spearmans rank correlation coefficient (rho) | | | | P-value 4-OH-PLA | P-value PLA | P-value ILA | P-value IAld | FDR$ P-value 4-OH-PLA | FDR$ P-value PLA | FDR$ P-value ILA | FDR$ P-value IAld |
|---|---|---|---|---|---|---|---|---|---|---|---|---|
| | 4-OH-PLA | PLA | ILA | IAld | | | | | | | | |
| g__Bifidobacterium | 0.442 | 0.284 | 0.548 | 0.416 | **0.0004498** | **0.0290239** | **0.0000070** | **0.0010635** | **0.008555** | 0.2495558 | **0.000175** | **0.0420525** |
| g__Actinomyces | 0.146 | 0.080 | 0.311 | 0.048 | 0.2704930 | 0.5458662 | **0.0165086** | 0.7157258 | 0.5201788 | 0.9289962 | 0.1544508 | 0.7779628 |
| g__Veillonella | 0.186 | 0.033 | 0.306 | -0.184 | 0.1576286 | 0.8020437 | **0.0185341** | 0.1634214 | 0.3753062 | 0.9312139 | 0.1544508 | 0.5865694 |
| g__Enterococcus | 0.274 | -0.148 | 0.261 | 0.152 | **0.0354055** | 0.2645806 | **0.0461982** | 0.2495612 | 0.2372769 | 0.8819353 | 0.230991 | 0.5865694 |
| g__Megasphaera | 0.222 | -0.007 | 0.249 | 0.050 | 0.0909266 | 0.9596590 | 0.0576953 | 0.7051705 | 0.3030887 | 0.959659 | 0.2403971 | 0.7779628 |
| g__Lactobacillus | 0.173 | 0.079 | 0.151 | 0.218 | 0.1900106 | 0.5521014 | 0.2551058 | 0.0966935 | 0.3958554 | 0.9289962 | 0.5545778 | 0.5865694 |
| g__Sedimentibacter | 0.033 | 0.062 | 0.147 | 0.170 | 0.8048974 | 0.6422936 | 0.2679576 | 0.1989926 | 0.8435579 | 0.9289962 | 0.5558316 | 0.5865694 |
| g__Ruminococcus | -0.083 | 0.169 | 0.125 | 0.320 | 0.5299559 | 0.2017811 | 0.3471119 | **0.0133736** | 0.7999229 | 0.8502565 | 0.6427998 | 0.2228933 |
| g__Atopobium | 0.191 | 0.149 | 0.119 | 0.079 | 0.1479732 | 0.2597555 | 0.3707286 | 0.5528294 | 0.369933 | 0.8819353 | 0.6602473 | 0.7725188 |
| g__SMB53 | -0.036 | 0.287 | 0.102 | -0.139 | 0.7840195 | **0.0274803** | 0.4400361 | 0.2932847 | 0.8435579 | 0.2495558 | 0.6602473 | 0.5865694 |
| g__Peptostreptococcus | -0.074 | -0.047 | 0.102 | 0.057 | 0.5782031 | 0.7236409 | 0.4438904 | 0.6704305 | 0.7999229 | 0.9290546 | 0.6602473 | 0.7779628 |
| g__Dialister | -0.169 | 0.023 | 0.101 | 0.063 | 0.1993905 | 0.8613919 | 0.4468355 | 0.6334654 | 0.398781 | 0.9312139 | 0.6602473 | 0.7725188 |
| g__Streptococcus | 0.080 | 0.046 | 0.099 | -0.026 | 0.5482163 | 0.7289691 | 0.4575813 | 0.8455403 | 0.7999229 | 0.9290546 | 0.6602473 | 0.8764227 |
| g__Epulopiscium | 0.066 | 0.037 | 0.098 | 0.126 | 0.6205657 | 0.7810028 | 0.4621731 | 0.3411654 | 0.7999229 | 0.9312139 | 0.6602473 | 0.6092239 |
| g__Trabulsiella | 0.065 | 0.017 | 0.085 | 0.129 | 0.6263253 | 0.8996419 | 0.5244414 | 0.3301630 | 0.7999229 | 0.9312139 | 0.7010209 | 0.6092239 |
| g__Dysgonomonas | -0.072 | 0.126 | 0.080 | -0.108 | 0.5860180 | 0.3402247 | 0.5453142 | 0.4175451 | 0.7999229 | 0.9289962 | 0.7010209 | 0.6734598 |
| g__Collinsella | -0.062 | 0.233 | 0.080 | 0.202 | 0.6403186 | 0.0755409 | 0.5467963 | 0.1248914 | 0.7999229 | 0.5395779 | 0.7010209 | 0.5865694 |
| g__Granulicatella | -0.032 | 0.030 | 0.071 | -0.178 | 0.8098156 | 0.8209402 | 0.5931955 | 0.1765097 | 0.8435579 | 0.9312139 | 0.7379984 | 0.5865694 |
| g__Citrobacter | -0.070 | -0.104 | 0.042 | -0.069 | 0.6002045 | 0.4326168 | 0.7517220 | 0.6031374 | 0.7999229 | 0.9289962 | 0.8584506 | 0.7725188 |
| g__Enterobacter | 0.048 | -0.080 | 0.034 | -0.064 | 0.7186160 | 0.5474336 | 0.7985486 | 0.6317140 | 0.8402048 | 0.9289962 | 0.8679876 | 0.7725188 |
| g__Clostridium | -0.034 | -0.067 | 0.006 | -0.068 | 0.7971909 | 0.6136399 | 0.9661666 | 0.6108373 | 0.8435579 | 0.9289962 | 0.9858843 | 0.7725188 |
| g__Escherichia | -0.090 | -0.212 | 0.002 | -0.065 | 0.4983883 | 0.1074015 | 0.9877483 | 0.6227996 | 0.7999229 | 0.6712594 | 0.9877483 | 0.7725188 |
| g__Haemophilus | 0.073 | -0.116 | -0.016 | -0.400 | 0.5831396 | 0.3820922 | 0.9027633 | 0.0016821 | 0.7999229 | 0.9289962 | 0.9403784 | 0.0420525 |
| g__Klebsiella | -0.100 | -0.060 | -0.027 | -0.175 | 0.4512175 | 0.6511640 | 0.8411621 | 0.1843987 | 0.7779612 | 0.9289962 | 0.8948533 | 0.5865694 |
| g__Eggerthella | -0.116 | 0.181 | -0.037 | 0.163 | 0.3821136 | 0.1706121 | 0.7824833 | 0.2168062 | 0.6823457 | 0.8502565 | 0.8679876 | 0.5865694 |
| g__Prevotella | -0.011 | 0.051 | -0.041 | -0.134 | 0.9322092 | 0.7021724 | 0.7554365 | 0.3134177 | 0.9322092 | 0.9290546 | 0.8584506 | 0.6027263 |
| g__Dorea | -0.235 | 0.020 | -0.050 | 0.053 | 0.0730098 | 0.8809158 | 0.7072053 | 0.6913053 | 0.2926457 | 0.9312139 | 0.8419111 | 0.7779628 |
| g__Megamonas | -0.039 | 0.057 | -0.069 | 0.298 | 0.7668089 | 0.6688773 | 0.6051587 | **0.0219974** | 0.8435579 | 0.9289962 | 0.7379984 | 0.2749675 |
| g__Faecalibacterium | -0.023 | 0.095 | -0.082 | 0.051 | 0.8637074 | 0.4735066 | 0.5362044 | 0.7030070 | 0.8813341 | 0.9289962 | 0.7010209 | 0.7779628 |
| g__Phascolarctobacterium | -0.059 | 0.283 | -0.103 | 0.148 | 0.6559368 | **0.0299467** | 0.4357441 | 0.2631798 | 0.7999229 | 0.2495558 | 0.6602473 | 0.5865694 |
| g__Slackia | -0.176 | -0.023 | -0.105 | -0.042 | 0.1831024 | 0.8619344 | 0.4278507 | 0.7513892 | 0.3958554 | 0.9312139 | 0.6602473 | 0.7993502 |
| g__Parabacteroides | -0.061 | 0.291 | -0.127 | 0.069 | 0.6444202 | **0.0253406** | 0.3375840 | 0.6027207 | 0.7999229 | 0.2495558 | 0.6427998 | 0.7725188 |
| g__Turicibacter | -0.124 | -0.015 | -0.144 | -0.142 | 0.3487486 | 0.9125896 | 0.2779158 | 0.2821248 | 0.6458307 | 0.9312139 | 0.5558316 | 0.5865694 |
| g__[Eubacterium] | -0.356 | 0.030 | -0.161 | -0.122 | **0.0056341** | 0.8201183 | 0.2224356 | 0.3558119 | 0.07042625 | 0.9312139 | 0.5055355 | 0.6134688 |
| g__Bacteroides | -0.202 | 0.102 | -0.173 | -0.023 | 0.1253999 | 0.4417915 | 0.1908746 | 0.8619881 | 0.3350045 | 0.9289962 | 0.4544633 | 0.8764227 |
| Others | -0.201 | 0.136 | -0.174 | 0.169 | 0.1273017 | 0.3060550 | 0.1877955 | 0.1998178 | 0.3350045 | 0.9289962 | 0.4544633 | 0.5865694 |
| g__Lachnospira | -0.261 | 0.162 | -0.177 | -0.189 | **0.0457670** | 0.2210667 | 0.1797372 | 0.1509646 | 0.238024 | 0.8502565 | 0.4544633 | 0.5865694 |
| g__Butyricicoccus | -0.230 | -0.190 | -0.177 | -0.150 | 0.0796357 | 0.1497693 | 0.1791126 | 0.2561422 | 0.2926457 | 0.8320517 | 0.4544633 | 0.5865694 |
| g__Pseudobutyrivibrio | -0.228 | 0.063 | -0.183 | -0.097 | 0.0819408 | 0.6332932 | 0.1663882 | 0.4664453 | 0.2926457 | 0.9289962 | 0.4544633 | 0.7288208 |
| g__Fusobacterium | -0.177 | -0.093 | -0.208 | -0.275 | 0.1807677 | 0.4837680 | 0.1142200 | **0.0351840** | 0.3958554 | 0.9289962 | 0.3569375 | 0.35184 |
| g__Blautia | -0.259 | -0.072 | -0.232 | -0.078 | **0.0476048** | 0.5881036 | 0.0772404 | 0.5563131 | 0.238024 | 0.9289962 | 0.257468 | 0.7725188 |
| g__Coprococcus | -0.271 | 0.081 | -0.235 | 0.021 | **0.0379643** | 0.5424958 | 0.0727595 | 0.8764227 | 0.2372769 | 0.9289962 | 0.257468 | 0.8764227 |
| g__Sutterella | -0.207 | 0.076 | -0.240 | -0.165 | 0.1165408 | 0.5649447 | 0.0675736 | 0.2112566 | 0.3350045 | 0.9289962 | 0.257468 | 0.5865694 |
| g__Roseburia | -0.341 | 0.112 | -0.250 | -0.184 | **0.0081292** | 0.3997124 | 0.0563907 | 0.1640897 | 0.081292 | 0.9289962 | 0.2403971 | 0.5865694 |
| g__Moryella | -0.230 | -0.087 | -0.262 | 0.083 | 0.0801815 | 0.5114224 | **0.0453861** | 0.5337881 | 0.2926457 | 0.9289962 | 0.230991 | 0.7725188 |
| g__Oscillospira | -0.279 | 0.167 | -0.266 | 0.141 | 0.0326268 | 0.2070223 | **0.0414742** | 0.2874382 | 0.2372769 | 0.8502565 | 0.230991 | 0.5865694 |
| g__Clostridium | -0.047 | -0.132 | -0.267 | -0.153 | 0.7225761 | 0.3203041 | **0.0407209** | 0.2466373 | 0.8402048 | 0.9289962 | 0.230991 | 0.5865694 |
| g__Lactococcus | -0.216 | -0.044 | -0.362 | 0.150 | 0.1005498 | 0.7432437 | **0.0048363** | 0.2571469 | 0.3142181 | 0.9290546 | **0.06045375** | 0.5865694 |
| g__[Ruminococcus] | -0.438 | -0.340 | -0.463 | -0.111 | **0.0005133** | **0.0085073** | **0.0002183** | 0.4020124 | **0.008555** | 0.2126825 | **0.003638333** | 0.6700207 |
| f__UnclassifiedLachnospiraceae | -0.477 | -0.376 | -0.593 | -0.202 | **0.0001328** | **0.0033377** | **0.0000007** | 0.1257732 | **0.00664** | 0.166885 | **0.000035** | 0.5865694 |

$False Discovery Corrected p-values, according to Benjamini & Hochberg, 1995

**Supplementary Data 1f. BLAST output of OTUs annotated as *Bifidobacterium* in the SKOT cohort study (n=59)**

| OTU ID[#] | GreenGenes taxonomy [RDP classifier 0.5] | 1st BLAST HIT | | | | | 2nd BLAST HIT | | | | Assigned taxonomy |
|---|---|---|---|---|---|---|---|---|---|---|---|
| | | % Identity | E-value | Query coverage | Gaps | Mismatch | % Identity | Query coverage | Gaps | Mismatch | |
| OTU_3 | Bifidobacterium;s__ | 99% *B. catenulatum* | 3.00E-68 | 100% | 1 | 0 | 97% *B. angulatum* | 100% | 1 | 3 | ***B. catenulatum* group** |
| | | 99% *B. pseudocatenulatum* | 3.00E-68 | 100% | 1 | 0 | | | | | |
| | | 99% *B. kashiwanohense* | 3.00E-68 | 100% | 1 | 0 | | | | | |
| OTU_4 | Bifidobacterium;s__longum | 99% *B. longum* subsp. *longum* | 2.00E-65 | 100% | 2 | 0 | 98% *B. longum* subsp. *infantis* | 100% | 2 | 1 | ***B. longum*** |
| | | | | | | | 98% *B. longum* subsp. *suillum* | 100% | 2 | 1 | |
| OTU_7 | Bifidobacterium;s__breve | 99% *B. breve* | 6.00E-70 | 100% | 1 | 0 | 94% *B. bifidum* | 100% | 4 | 4 | ***B. breve*** |
| OTU_23 | Bifidobacterium;s__ | 98% *B. scardovii* | 6.00E-65 | 100% | 2 | 1 | 97% *B. aerophillum* | 100% | 2 | 2 | ***B. scardovii*** |
| OTU_24 | Bifidobacterium;s__adolescentis | 97% *B. adolescentis* | 7.00E-64 | 100% | 4 | 0 | 96% *B. catenulatum* | 100% | 4 | 2 | ***B. adolescentis*** |
| | | 97% *B. faecale* | 7.00E-64 | 100% | 4 | 0 | 96% *B. pseudocatenulatum* | 100% | 4 | 2 | |
| | | 97% *B. stercoris* | 7.00E-64 | 100% | 4 | 0 | 96% *B. kashiwanohense* | 100% | 4 | 2 | |
| | | 97% *B. ruminantium* | 7.00E-64 | 100% | 4 | 0 | | | | | |
| OTU_56 | Bifidobacterium | 96% *B. breve* | 4.00E-61 | 100% | 4 | 2 | 92% *B. aerophillum* | 100% | 9 | 3 | ***B. breve*** |
| OTU_98 | Bifidobacterium;s__longum | 97% *B. longum* subsp. *infantis* | 1.00E-61 | 100% | 4 | 0 | 96% *B. longum* subsp. *longum* | 100% | 4 | 1 | ***B. longum*** |
| | | 97% *B. longum* subsp. *sullium* | 1.00E-61 | 100% | 4 | 0 | 96% *B. longum* subsp. *suis* | 100% | 4 | 1 | |
| OTU_107 | Bifidobacterium;s__longum | 97% *B. longum* subsp. *longum* | 1.00E-61 | 100% | 4 | 0 | 96% *B. longum* subsp. *infantis* | 100% | 4 | 1 | ***B. longum*** |
| | | | | | | | 96% *B. longum* subsp. *suillum* | 100% | 4 | 1 | |
| OTU_116 | Bifidobacterium;s__adolescentis | 99% *B. adolescentis* | 3.00E-68 | 100% | 2 | 0 | 97% *B. catenulatum* | 100% | 2 | 2 | ***B. adolescentis*** |
| | | 99% *B. faecale* | 3.00E-68 | 100% | 2 | 0 | 97% *B. pseudocatenulatum* | 100% | 2 | 2 | |
| | | 99% *B. stercoris* | 3.00E-68 | 100% | 2 | 0 | 97% *B. kashiwanohense* | 100% | 2 | 2 | |
| | | 99% *B. ruminantium* | 3.00E-68 | 100% | 2 | 0 | | | | | |
| OTU_129 | Bifidobacterium;s__adolescentis | 97% *B. adolescentis* | 2.00E-64 | 100% | 4 | 0 | 96% *B. catenulatum* | 100% | 4 | 2 | ***B. adolescentis*** |
| | | 97% *B. faecale* | 2.00E-64 | 100% | 4 | 0 | 96% *B. pseudocatenulatum* | 100% | 4 | 2 | |
| | | 97% *B. stercoris* | 2.00E-64 | 100% | 4 | 0 | 96% *B. kashiwanohense* | 100% | 4 | 2 | |
| | | 97% *B. ruminantium* | 2.00E-64 | 100% | 4 | 0 | | | | | |
| OTU_162 | Bifidobacterium;s__breve | 97% *B. breve* | 4.00E-66 | 100% | 2 | 1 | 92% *B. stellenboschense* | 100% | 3 | 9 | ***B. breve*** |
| OTU_215 | Bifidobacterium | 97% *B. catenulatum* | 4.00E-61 | 100% | 5 | 0 | 94% *B. angulatum* | 100% | 5 | 3 | ***B. catenulatum* group** |
| | | 97% *B. pseudocatenulatum* | 4.00E-61 | 100% | 5 | 0 | | | | | |
| | | 97% *B. kashiwanohense* | 4.00E-61 | 100% | 5 | 0 | | | | | |
| OTU_217 | Bifidobacterium;s__ | 98% *B. bifidum* | 2.00E-65 | 100% | 0 | 3 | 97% *B. catenulatum* | 100% | 0 | 4 | ***B. bifidum*** |
| | | | | | | | 97% *B. pseudocatenulatum* | 100% | 0 | 4 | |
| | | | | | | | 97% *B. kashiwanohense* | 100% | 0 | 4 | |
| OTU_225 | Bifidobacterium;s__ | 99% *B. dentium* | 1.00E-67 | 100% | 2 | 0 | 97% *B. boum* | 100% | 2 | 2 | ***B. dentium*** |
| | | | | | | | 97% *B. thermophilum* | 100% | 2 | 2 | |
| OTU_246 | Bifidobacterium;s__bifidum | 98% *B. bifidum* | 6.00E-65 | 100% | 1 | 2 | 95% *B. boum* | 100% | 3 | 4 | ***B. bifidum*** |
| | | | | | | | 95% *B. thermophilum* | 100% | 3 | 4 | |
| OTU_255 | Bifidobacterium;s__ | 98% *B. catenulatum* | 6.00E-65 | 100% | 3 | 0 | 96% *B. angulatum* | 100% | 3 | 3 | ***B. catenulatum* group** |
| | | 98% *B. pseudocatenulatum* | 6.00E-65 | 100% | 3 | 0 | | | | | |
| | | 98% *B. kashiwanohense* | 6.00E-65 | 100% | 3 | 0 | | | | | |
| OTU_345 | Bifidobacterium;s__breve | 97% *B. breve* | 7.00E-64 | 100% | 3 | 2 | 91% *B. biavatii* | 100% | 4 | 9 | ***B. breve*** |
| OTU_371 | Bifidobacterium;s__pseudolongum | 98% *B. animalis* subsp. *lactis* | 2.00E-69 | 98% | 3 | 0 | 94% *B. choerinum* | 98% | 5 | 4 | ***B. animalis/pseudolongum*** |
| | | 98% *B. pseudolongum* | 2.00E-69 | 98% | 3 | 0 | | | | | |
| OTU_425 | Bifidobacterium;s__bifidum | 98% *B. bifidum* | 6.00E-65 | 100% | 3 | 0 | 94% *B. boum* | 100% | 3 | 5 | ***B. bifidum*** |
| | | | | | | | 94% *B.thermophilum* | 100% | 3 | 5 | |
| OTU_439 | Bifidobacterium;s__bifidum | 99% *B. bifidum* | 1.00E-66 | 100% | 2 | 0 | 96% *B. boum* | 100% | 4 | 2 | ***B. bifidum*** |
| | | | | | | | 96% *B. thermophilum* | 100% | 4 | 2 | |
| OTU_487 | Bifidobacterium;s__bifidum | 99% *B. bifidum* | 1.00E-66 | 100% | 2 | 0 | 95% *B. boum* | 100% | 4 | 3 | ***B. bifidum*** |
| | | | | | | | 95% *B.thermophilum* | 100% | 4 | 3 | |
| OTU_495 | Bifidobacterium;s__longum | 96% B. *longum* subsp. *infantis* | 2.00E-60 | 100% | 5 | 0 | 96% *B. longum* subsp. *longum* | 100% | 5 | 1 | ***B. longum*** |
| | | 96% B. *longum* subsp. *sullium* | 2.00E-60 | 100% | 5 | 0 | 96% *B. longum* subsp. *suis* | 100% | 5 | 1 | |
| OTU_584 | Bifidobacterium;s__ | 96% *B. dentium* | 2.00E-60 | 100% | 3 | 2 | 95% *B. boum* | 100% | 3 | 3 | ***B. dentium*** |
| | | | | | | | 95% *B.thermophilum* | 100% | 3 | 3 | |

#Only OTUs representing relative abundance >0.1% within the total relative abundance of *Bifidobacterium* genus were included
