## Supplementary Data 2 for "Breastmilk-promoted bifidobacteria produce aromatic amino acids in the infant gut"

**Supplementary Data 2a. Metadata of the Copenhagen Infant Gut cohort study**

| Participant | Sample_ID | Birth mode | Gender | Birth classification | Twin | Age_feces (days) | Age_feces (weeks) | Breast feeding | Formula feeding | Solid food introduced | Probiotics | Antibiotics |
|---|---|---|---|---|---|---|---|---|---|---|---|---|
| CIG01 | CIG01-1 | Vaginal | Female | Full-term | No | 30 | 4.3 | Yes | No | No | No | No |
| CIG01 | CIG01-2 | Vaginal | Female | Full-term | No | 45 | 6.4 | Yes | No | No | No | No |
| CIG01 | CIG01-3 | Vaginal | Female | Full-term | No | 57 | 8.1 | Yes | No | No | No | No |
| CIG01 | CIG01-4 | Vaginal | Female | Full-term | No | 73 | 10.4 | Yes | No | No | No | No |
| CIG01 | CIG01-5 | Vaginal | Female | Full-term | No | 94 | 13.4 | Yes | No | No | No | No |
| CIG01 | CIG01-6 | Vaginal | Female | Full-term | No | 112 | 16.0 | Yes | Yes | No | No | No |
| CIG01 | CIG01-7 | Vaginal | Female | Full-term | No | 138 | 19.7 | Yes | No | No | No | No |
| CIG01 | CIG01-8 | Vaginal | Female | Full-term | No | 155 | 22.1 | Yes | No | No | No | No |
| CIG01 | CIG01-9 | Vaginal | Female | Full-term | No | 172 | 24.6 | Yes | Yes | Yes | No | No |
| CIG01 | CIG01-10 | Vaginal | Female | Full-term | No | 192 | 27.4 | No | Yes | Yes | No | No |
| CIG02 | CIG02-1 | Vaginal | Male | Full-term | No | 12 | 1.7 | Yes | No | No | No | No |
| CIG02 | CIG02-2 | Vaginal | Male | Full-term | No | 20 | 2.9 | Yes | Yes | No | No | No |
| CIG02 | CIG02-3 | Vaginal | Male | Full-term | No | 34 | 4.9 | Yes | No | No | No | No |
| CIG02 | CIG02-4 | Vaginal | Male | Full-term | No | 45 | 6.4 | Yes | No | No | No | No |
| CIG02 | CIG02-5 | Vaginal | Male | Full-term | No | 54 | 7.7 | Yes | No | No | No | No |
| CIG02 | CIG02-6 | Vaginal | Male | Full-term | No | 68 | 9.7 | Yes | Yes | No | No | No |
| CIG02 | CIG02-7 | Vaginal | Male | Full-term | No | 82 | 11.7 | Yes | Yes | No | No | No |
| CIG02 | CIG02-8 | Vaginal | Male | Full-term | No | 103 | 14.7 | Yes | Yes | No | No | No |
| CIG02 | CIG02-9 | Vaginal | Male | Full-term | No | 116 | 16.6 | Yes | Yes | No | No | No |
| CIG02 | CIG02-10 | Vaginal | Male | Full-term | No | 146 | 20.9 | Yes | Yes | Yes | No | No |
| CIG02 | CIG02-11 | Vaginal | Male | Full-term | No | 178 | 25.4 | Yes | Yes | Yes | No | No |
| CIG03 | CIG03-1 | Vaginal | Male | Full-term | No | 1 | 0.1 | Yes | No | No | No | No |
| CIG03 | CIG03-2 | Vaginal | Male | Full-term | No | 4 | 0.6 | Yes | No | No | No | No |
| CIG03 | CIG03-3 | Vaginal | Male | Full-term | No | 16 | 2.3 | Yes | No | No | No | No |
| CIG03 | CIG03-4 | Vaginal | Male | Full-term | No | 29 | 4.1 | Yes | No | No | No | No |
| CIG03 | CIG03-5 | Vaginal | Male | Full-term | No | 58 | 8.3 | Yes | No | No | No | No |
| CIG03 | CIG03-6 | Vaginal | Male | Full-term | No | 69 | 9.9 | Yes | No | No | No | No |
| CIG03 | CIG03-7 | Vaginal | Male | Full-term | No | 84 | 12.0 | Yes | No | No | No | No |
| CIG03 | CIG03-8 | Vaginal | Male | Full-term | No | 96 | 13.7 | Yes | No | No | No | No |
| CIG03 | CIG03-9 | Vaginal | Male | Full-term | No | 115 | 16.4 | Yes | No | No | No | No |
| CIG03 | CIG03-10 | Vaginal | Male | Full-term | No | 141 | 20.1 | Yes | Yes | No | No | No |
| CIG04 | CIG04-1 | Vaginal | Male | Full-term | No | 2 | 0.3 | Yes | Yes | No | No | No |
| CIG04 | CIG04-2 | Vaginal | Male | Full-term | No | 13 | 1.9 | Yes | No | No | No | No |
| CIG04 | CIG04-3 | Vaginal | Male | Full-term | No | 28 | 4.0 | Yes | No | No | No | No |
| CIG04 | CIG04-4 | Vaginal | Male | Full-term | No | 42 | 6.0 | Yes | No | No | No | No |
| CIG04 | CIG04-5 | Vaginal | Male | Full-term | No | 55 | 7.9 | Yes | No | No | No | No |
| CIG04 | CIG04-6 | Vaginal | Male | Full-term | No | 67 | 9.6 | Yes | No | No | No | No |
| CIG04 | CIG04-7 | Vaginal | Male | Full-term | No | 82 | 11.7 | Yes | No | No | No | No |
| CIG04 | CIG04-8 | Vaginal | Male | Full-term | No | 96 | 13.7 | Yes | No | No | No | No |
| CIG04 | CIG04-9 | Vaginal | Male | Full-term | No | 110 | 15.7 | Yes | No | No | No | No |
| CIG04 | CIG04-10 | Vaginal | Male | Full-term | No | 141 | 20.1 | Yes | No | No | No | No |
| CIG04 | CIG04-11 | Vaginal | Male | Full-term | No | 164 | 23.4 | Yes | No | Yes | No | No |
| CIG05 | CIG05-1 | Vaginal | Male | Full-term | No | 6 | 0.9 | Yes | Yes | No | No | No |
| CIG05 | CIG05-2 | Vaginal | Male | Full-term | No | 17 | 2.4 | Yes | Yes | No | No | No |
| CIG05 | CIG05-3 | Vaginal | Male | Full-term | No | 34 | 4.9 | Yes | Yes | No | No | Yes |
| CIG05 | CIG05-4 | Vaginal | Male | Full-term | No | 52 | 7.4 | Yes | Yes | No | No | No |
| CIG05 | CIG05-5 | Vaginal | Male | Full-term | No | 66 | 9.4 | Yes | Yes | No | No | No |
| CIG05 | CIG05-6 | Vaginal | Male | Full-term | No | 82 | 11.7 | Yes | Yes | No | No | No |
| CIG05 | CIG05-7 | Vaginal | Male | Full-term | No | 96 | 13.7 | Yes | Yes | No | No | No |
| CIG05 | CIG05-8 | Vaginal | Male | Full-term | No | 119 | 17.0 | Yes | Yes | Yes | No | No |
| CIG05 | CIG05-9 | Vaginal | Male | Full-term | No | 133 | 19.0 | No | Yes | Yes | No | No |
| CIG05 | CIG05-10 | Vaginal | Male | Full-term | No | 176 | 25.1 | No | Yes | Yes | No | No |
| CIG05 | CIG05-11 | Vaginal | Male | Full-term | No | 192 | 27.4 | No | Yes | Yes | No | No |
| CIG06 | CIG06-1 | Vaginal | Female | Full-term | No | 2 | 0.3 | Yes | No | No | No | No |
| CIG06 | CIG06-2 | Vaginal | Female | Full-term | No | 14 | 2.0 | Yes | No | No | No | No |
| CIG06 | CIG06-3 | Vaginal | Female | Full-term | No | 29 | 4.1 | Yes | No | No | Yes | No |
| CIG06 | CIG06-4 | Vaginal | Female | Full-term | No | 41 | 5.9 | Yes | No | No | Yes | No |
| CIG06 | CIG06-5 | Vaginal | Female | Full-term | No | 56 | 8.0 | Yes | No | No | No | No |
| CIG06 | CIG06-6 | Vaginal | Female | Full-term | No | 68 | 9.7 | Yes | No | No | No | No |
| CIG06 | CIG06-7 | Vaginal | Female | Full-term | No | 83 | 11.9 | Yes | No | No | No | No |
| CIG06 | CIG06-8 | Vaginal | Female | Full-term | No | 93 | 13.3 | Yes | No | No | No | No |
| CIG06 | CIG06-9 | Vaginal | Female | Full-term | No | 109 | 15.6 | Yes | No | No | No | No |
| CIG06 | CIG06-10 | Vaginal | Female | Full-term | No | 137 | 19.6 | Yes | No | No | No | No |
| CIG06 | CIG06-11 | Vaginal | Female | Full-term | No | 165 | 23.6 | Yes | No | Yes | No | No |
| CIG07 | CIG07-1 | Vaginal | Female | Full-term | No | 2 | 0.3 | Yes | Yes | No | No | No |
| CIG07 | CIG07-2 | Vaginal | Female | Full-term | No | 16 | 2.3 | Yes | Yes | No | No | No |
| CIG07 | CIG07-3 | Vaginal | Female | Full-term | No | 30 | 4.3 | Yes | Yes | No | No | No |

| | | | | | | | | | | | | |
|---|---|---|---|---|---|---|---|---|---|---|---|---|
| CIG07 | CIG07-4 | Vaginal | Female | Full-term | No | 43 | 6.1 | Yes | Yes | No | No | No |
| CIG07 | CIG07-5 | Vaginal | Female | Full-term | No | 57 | 8.1 | Yes | Yes | No | No | No |
| CIG07 | CIG07-6 | Vaginal | Female | Full-term | No | 71 | 10.1 | Yes | Yes | No | No | No |
| CIG07 | CIG07-7 | Vaginal | Female | Full-term | No | 85 | 12.1 | Yes | Yes | No | No | No |
| CIG07 | CIG07-8 | Vaginal | Female | Full-term | No | 99 | 14.1 | Yes | Yes | No | No | No |
| CIG07 | CIG07-9 | Vaginal | Female | Full-term | No | 112 | 16.0 | Yes | Yes | No | No | No |
| CIG07 | CIG07-10 | Vaginal | Female | Full-term | No | 146 | 20.9 | Yes | Yes | No | No | No |
| CIG07 | CIG07-11 | Vaginal | Female | Full-term | No | 171 | 24.4 | Yes | Yes | Yes | No | No |
| CIG08 | CIG08-1 | C-section | Female | Preterm | Yes monozygotic with CIG09 | 6 | 0.9 | Yes | Yes | No | No | No |
| CIG08 | CIG08-2 | C-section | Female | Preterm | Yes monozygotic with CIG09 | 19 | 2.7 | Yes | No | No | No | No |
| CIG08 | CIG08-3 | C-section | Female | Preterm | Yes monozygotic with CIG09 | 30 | 4.3 | Yes | No | No | No | No |
| CIG08 | CIG08-4 | C-section | Female | Preterm | Yes monozygotic with CIG09 | 44 | 6.3 | Yes | No | No | No | No |
| CIG08 | CIG08-5 | C-section | Female | Preterm | Yes monozygotic with CIG09 | 56 | 8.0 | Yes | No | No | No | No |
| CIG08 | CIG08-6 | C-section | Female | Preterm | Yes monozygotic with CIG09 | 71 | 10.1 | Yes | No | No | No | No |
| CIG08 | CIG08-7 | C-section | Female | Preterm | Yes monozygotic with CIG09 | 84 | 12.0 | Yes | No | No | No | No |
| CIG08 | CIG08-8 | C-section | Female | Preterm | Yes monozygotic with CIG09 | 99 | 14.1 | Yes | No | No | No | No |
| CIG08 | CIG08-9 | C-section | Female | Preterm | Yes monozygotic with CIG09 | 113 | 16.1 | Yes | No | No | No | No |
| CIG08 | CIG08-10 | C-section | Female | Preterm | Yes monozygotic with CIG09 | 138 | 19.7 | Yes | No | No | No | No |
| CIG08 | CIG08-11 | C-section | Female | Preterm | Yes monozygotic with CIG09 | 168 | 24.0 | Yes | No | No | No | No |
| CIG09 | CIG09-1 | C-section | Female | Preterm | Yes monozygotic with CIG08 | 6 | 0.9 | Yes | Yes | No | No | No |
| CIG09 | CIG09-2 | C-section | Female | Preterm | Yes monozygotic with CIG08 | 20 | 2.9 | Yes | No | No | No | No |
| CIG09 | CIG09-3 | C-section | Female | Preterm | Yes monozygotic with CIG08 | 31 | 4.4 | Yes | No | No | No | No |
| CIG09 | CIG09-4 | C-section | Female | Preterm | Yes monozygotic with CIG08 | 44 | 6.3 | Yes | No | No | No | No |

| | | | | | | | | | | | | |
|---|---|---|---|---|---|---|---|---|---|---|---|---|
| CIG09 | CIG09-5 | C-section | Female | Preterm | Yes monozygotic with CIG08 | 58 | 8.3 | Yes | No | No | No | No |
| CIG09 | CIG09-6 | C-section | Female | Preterm | Yes monozygotic with CIG08 | 70 | 10.0 | Yes | No | No | No | No |
| CIG09 | CIG09-7 | C-section | Female | Preterm | Yes monozygotic with CIG08 | 86 | 12.3 | Yes | No | No | No | No |
| CIG09 | CIG09-8 | C-section | Female | Preterm | Yes monozygotic with CIG08 | 101 | 14.4 | Yes | No | No | No | No |
| CIG09 | CIG09-9 | C-section | Female | Preterm | Yes monozygotic with CIG08 | 112 | 16.0 | Yes | No | No | No | No |
| CIG09 | CIG09-10 | C-section | Female | Preterm | Yes monozygotic with CIG08 | 138 | 19.7 | Yes | No | No | No | No |
| CIG09 | CIG09-11 | C-section | Female | Preterm | Yes monozygotic with CIG08 | 166 | 23.7 | Yes | No | No | No | No |
| CIG10 | CIG10-1 | Vaginal | Female | Full-term | No | 6 | 0.9 | Yes | No | No | No | No |
| CIG10 | CIG10-2 | Vaginal | Female | Full-term | No | 17 | 2.4 | Yes | No | No | No | No |
| CIG10 | CIG10-3 | Vaginal | Female | Full-term | No | 33 | 4.7 | Yes | No | No | No | No |
| CIG10 | CIG10-4 | Vaginal | Female | Full-term | No | 46 | 6.6 | Yes | No | No | No | No |
| CIG10 | CIG10-5 | Vaginal | Female | Full-term | No | 57 | 8.1 | Yes | No | No | No | Yes |
| CIG10 | CIG10-6 | Vaginal | Female | Full-term | No | 76 | 10.9 | Yes | No | No | No | No |
| CIG10 | CIG10-7 | Vaginal | Female | Full-term | No | 87 | 12.4 | Yes | No | No | No | No |
| CIG10 | CIG10-8 | Vaginal | Female | Full-term | No | 107 | 15.3 | Yes | No | No | No | No |
| CIG10 | CIG10-9 | Vaginal | Female | Full-term | No | 111 | 15.9 | Yes | No | No | No | No |
| CIG10 | CIG10-10 | Vaginal | Female | Full-term | No | 131 | 18.7 | Yes | No | No | No | No |
| CIG10 | CIG10-11 | Vaginal | Female | Full-term | No | 162 | 23.1 | Yes | No | No | No | No |
| CIG11 | CIG11-1 | Vaginal | Male | Full-term | No | 3 | 0.4 | Yes | No | No | No | No |
| CIG11 | CIG11-2 | Vaginal | Male | Full-term | No | 17 | 2.4 | Yes | No | No | No | No |
| CIG11 | CIG11-3 | Vaginal | Male | Full-term | No | 32 | 4.6 | Yes | No | No | No | No |
| CIG11 | CIG11-4 | Vaginal | Male | Full-term | No | 47 | 6.7 | Yes | No | No | No | No |
| CIG11 | CIG11-5 | Vaginal | Male | Full-term | No | 61 | 8.7 | Yes | No | No | No | No |
| CIG11 | CIG11-6 | Vaginal | Male | Full-term | No | 75 | 10.7 | Yes | No | No | No | No |
| CIG11 | CIG11-7 | Vaginal | Male | Full-term | No | 91 | 13.0 | Yes | No | No | No | No |
| CIG11 | CIG11-8 | Vaginal | Male | Full-term | No | 107 | 15.3 | Yes | No | No | No | No |
| CIG11 | CIG11-9 | Vaginal | Male | Full-term | No | 121 | 17.3 | Yes | No | No | No | No |
| CIG11 | CIG11-10 | Vaginal | Male | Full-term | No | 150 | 21.4 | Yes | No | No | No | No |
| CIG11 | CIG11-11 | Vaginal | Male | Full-term | No | 178 | 25.4 | Yes | No | No | No | No |
| CIG12 | CIG12-1 | Vaginal | Male | Full-term | No | 3 | 0.4 | Yes | No | No | No | No |
| CIG12 | CIG12-2 | Vaginal | Male | Full-term | No | 16 | 2.3 | Yes | No | No | No | No |
| CIG12 | CIG12-3 | Vaginal | Male | Full-term | No | 29 | 4.1 | Yes | No | No | No | No |
| CIG12 | CIG12-4 | Vaginal | Male | Full-term | No | 43 | 6.1 | Yes | No | No | No | No |
| CIG12 | CIG12-5 | Vaginal | Male | Full-term | No | 59 | 8.4 | Yes | No | No | No | No |
| CIG12 | CIG12-6 | Vaginal | Male | Full-term | No | 72 | 10.3 | Yes | Yes | No | No | No |
| CIG12 | CIG12-7 | Vaginal | Male | Full-term | No | 85 | 12.1 | Yes | No | No | No | No |
| CIG12 | CIG12-8 | Vaginal | Male | Full-term | No | 102 | 14.6 | Yes | No | No | No | No |
| CIG12 | CIG12-9 | Vaginal | Male | Full-term | No | 118 | 16.9 | Yes | No | No | No | No |
| CIG12 | CIG12-10 | Vaginal | Male | Full-term | No | 139 | 19.9 | Yes | Yes | Yes | No | No |
| CIG12 | CIG12-11 | Vaginal | Male | Full-term | No | 170 | 24.3 | Yes | Yes | Yes | No | No |
| CIG13 | CIG13-1 | Vaginal | Male | Full-term | No | 2 | 0.3 | Yes | No | No | No | No |
| CIG13 | CIG13-2 | Vaginal | Male | Full-term | No | 12 | 1.7 | Yes | No | No | No | No |
| CIG13 | CIG13-3 | Vaginal | Male | Full-term | No | 26 | 3.7 | Yes | No | No | No | No |
| CIG13 | CIG13-4 | Vaginal | Male | Full-term | No | 40 | 5.7 | Yes | No | No | No | No |
| CIG13 | CIG13-5 | Vaginal | Male | Full-term | No | 56 | 8.0 | Yes | No | No | No | No |
| CIG13 | CIG13-6 | Vaginal | Male | Full-term | No | 68 | 9.7 | Yes | No | No | No | No |
| CIG13 | CIG13-7 | Vaginal | Male | Full-term | No | 88 | 12.6 | Yes | No | No | No | No |
| CIG13 | CIG13-8 | Vaginal | Male | Full-term | No | 96 | 13.7 | Yes | No | No | No | No |
| CIG13 | CIG13-9 | Vaginal | Male | Full-term | No | 107 | 15.3 | Yes | No | No | No | No |

| | | | | | | | | | | | | |
|---|---|---|---|---|---|---|---|---|---|---|---|---|
| CIG13 | CIG13-10 | Vaginal | Male | Full-term | No | 133 | 19.0 | Yes | No | No | No | No |
| CIG13 | CIG13-11 | Vaginal | Male | Full-term | No | 167 | 23.9 | Yes | No | No | No | No |
| CIG14 | CIG14-1 | Vaginal | Male | Full-term | No | 5 | 0.7 | Yes | No | No | No | No |
| CIG14 | CIG14-2 | Vaginal | Male | Full-term | No | 16 | 2.3 | Yes | No | No | No | No |
| CIG14 | CIG14-3 | Vaginal | Male | Full-term | No | 30 | 4.3 | Yes | No | No | No | No |
| CIG14 | CIG14-4 | Vaginal | Male | Full-term | No | 52 | 7.4 | Yes | No | No | No | No |
| CIG14 | CIG14-5 | Vaginal | Male | Full-term | No | 68 | 9.7 | Yes | No | No | No | No |
| CIG14 | CIG14-6 | Vaginal | Male | Full-term | No | 78 | 11.1 | Yes | No | No | No | No |
| CIG14 | CIG14-7 | Vaginal | Male | Full-term | No | 96 | 13.7 | Yes | No | No | No | No |
| CIG14 | CIG14-8 | Vaginal | Male | Full-term | No | 109 | 15.6 | Yes | No | No | No | No |
| CIG14 | CIG14-9 | Vaginal | Male | Full-term | No | 125 | 17.9 | Yes | No | Yes | No | No |
| CIG14 | CIG14-10 | Vaginal | Male | Full-term | No | 146 | 20.9 | Yes | No | Yes | No | No |
| CIG14 | CIG14-11 | Vaginal | Male | Full-term | No | 166 | 23.7 | Yes | No | Yes | No | No |
| CIG15 | CIG15-1 | Vaginal | Male | Full-term | No | 1 | 0.1 | Yes | No | No | No | No |
| CIG15 | CIG15-2 | Vaginal | Male | Full-term | No | 13 | 1.9 | Yes | Yes | No | No | No |
| CIG15 | CIG15-3 | Vaginal | Male | Full-term | No | 30 | 4.3 | Yes | Yes | No | No | No |
| CIG15 | CIG15-4 | Vaginal | Male | Full-term | No | 44 | 6.3 | No | Yes | No | No | No |
| CIG15 | CIG15-5 | Vaginal | Male | Full-term | No | 58 | 8.3 | No | Yes | No | No | No |
| CIG15 | CIG15-6 | Vaginal | Male | Full-term | No | 72 | 10.3 | No | Yes | No | No | No |
| CIG15 | CIG15-7 | Vaginal | Male | Full-term | No | 84 | 12.0 | No | Yes | No | No | No |
| CIG15 | CIG15-8 | Vaginal | Male | Full-term | No | 100 | 14.3 | No | Yes | No | No | No |
| CIG15 | CIG15-9 | Vaginal | Male | Full-term | No | 113 | 16.1 | No | Yes | No | No | No |
| CIG15 | CIG15-10 | Vaginal | Male | Full-term | No | 141 | 20.1 | No | Yes | Yes | No | No |
| CIG15 | CIG15-11 | Vaginal | Male | Full-term | No | 167 | 23.9 | No | Yes | Yes | No | No |
| CIG16 | CIG16-1 | Vaginal | Male | Full-term | No | 6 | 0.9 | Yes | No | No | Yes | No |
| CIG16 | CIG16-2 | Vaginal | Male | Full-term | No | 19 | 2.7 | Yes | No | No | Yes | No |
| CIG16 | CIG16-3 | Vaginal | Male | Full-term | No | 40 | 5.7 | Yes | No | No | Yes | No |
| CIG16 | CIG16-4 | Vaginal | Male | Full-term | No | 56 | 8.0 | Yes | No | No | Yes | No |
| CIG16 | CIG16-5 | Vaginal | Male | Full-term | No | 72 | 10.3 | Yes | No | No | Yes | No |
| CIG16 | CIG16-6 | Vaginal | Male | Full-term | No | 88 | 12.6 | Yes | No | No | Yes | Yes |
| CIG16 | CIG16-7 | Vaginal | Male | Full-term | No | 103 | 14.7 | Yes | No | No | Yes | No |
| CIG16 | CIG16-8 | Vaginal | Male | Full-term | No | 117 | 16.7 | Yes | No | No | Yes | No |
| CIG16 | CIG16-9 | Vaginal | Male | Full-term | No | 172 | 24.6 | Yes | No | No | Yes | No |
| CIG17 | CIG17-1 | Vaginal | Male | Full-term | No | 2 | 0.3 | Yes | No | No | No | No |
| CIG17 | CIG17-2 | Vaginal | Male | Full-term | No | 15 | 2.1 | Yes | No | No | No | No |
| CIG17 | CIG17-3 | Vaginal | Male | Full-term | No | 30 | 4.3 | Yes | No | No | No | No |
| CIG17 | CIG17-4 | Vaginal | Male | Full-term | No | 51 | 7.3 | Yes | No | No | No | No |
| CIG17 | CIG17-5 | Vaginal | Male | Full-term | No | 57 | 8.1 | Yes | No | No | No | No |
| CIG17 | CIG17-6 | Vaginal | Male | Full-term | No | 73 | 10.4 | Yes | No | No | No | No |
| CIG17 | CIG17-7 | Vaginal | Male | Full-term | No | 84 | 12.0 | Yes | No | No | No | No |
| CIG17 | CIG17-8 | Vaginal | Male | Full-term | No | 97 | 13.9 | Yes | No | No | No | No |
| CIG17 | CIG17-9 | Vaginal | Male | Full-term | No | 110 | 15.7 | Yes | No | No | No | No |
| CIG17 | CIG17-10 | Vaginal | Male | Full-term | No | 139 | 19.9 | Yes | No | No | No | No |
| CIG18 | CIG18-1 | Vaginal | Female | Full-term | Yes monozygotic with CIG19 | 4 | 0.6 | Yes | Yes | No | No | No |
| CIG18 | CIG18-2 | Vaginal | Female | Full-term | Yes monozygotic with CIG19 | 15 | 2.1 | Yes | No | No | No | No |
| CIG18 | CIG18-3 | Vaginal | Female | Full-term | Yes monozygotic with CIG19 | 34 | 4.9 | Yes | No | No | No | No |
| CIG18 | CIG18-4 | Vaginal | Female | Full-term | Yes monozygotic with CIG19 | 47 | 6.7 | Yes | No | No | No | No |
| CIG18 | CIG18-5 | Vaginal | Female | Full-term | Yes monozygotic with CIG19 | 60 | 8.6 | Yes | No | No | No | No |
| CIG18 | CIG18-6 | Vaginal | Female | Full-term | Yes monozygotic with CIG19 | 77 | 11.0 | Yes | No | No | No | No |

| | | | | | | | | | | | | |
|---|---|---|---|---|---|---|---|---|---|---|---|---|
| CIG18 | CIG18-7 | Vaginal | Female | Full-term | Yes monozygotic with CIG19 | 87 | 12.4 | Yes | No | No | No | No |
| CIG18 | CIG18-8 | Vaginal | Female | Full-term | Yes monozygotic with CIG19 | 101 | 14.4 | Yes | No | No | No | No |
| CIG18 | CIG18-9 | Vaginal | Female | Full-term | Yes monozygotic with CIG19 | 118 | 16.9 | Yes | No | No | No | No |
| CIG18 | CIG18-10 | Vaginal | Female | Full-term | Yes monozygotic with CIG19 | 143 | 20.4 | Yes | No | Yes | No | No |
| CIG18 | CIG18-11 | Vaginal | Female | Full-term | Yes monozygotic with CIG19 | 172 | 24.6 | Yes | No | Yes | No | No |
| CIG19 | CIG19-1 | Vaginal | Female | Full-term | Yes monozygotic with CIG18 | 4 | 0.6 | Yes | Yes | No | No | No |
| CIG19 | CIG19-2 | Vaginal | Female | Full-term | Yes monozygotic with CIG18 | 15 | 2.1 | Yes | No | No | No | No |
| CIG19 | CIG19-3 | Vaginal | Female | Full-term | Yes monozygotic with CIG18 | 34 | 4.9 | Yes | No | No | No | No |
| CIG19 | CIG19-4 | Vaginal | Female | Full-term | Yes monozygotic with CIG18 | 47 | 6.7 | Yes | No | No | No | No |
| CIG19 | CIG19-5 | Vaginal | Female | Full-term | Yes monozygotic with CIG18 | 60 | 8.6 | Yes | No | No | No | No |
| CIG19 | CIG19-6 | Vaginal | Female | Full-term | Yes monozygotic with CIG18 | 77 | 11.0 | Yes | No | No | No | No |
| CIG19 | CIG19-7 | Vaginal | Female | Full-term | Yes monozygotic with CIG18 | 89 | 12.7 | Yes | No | No | No | No |
| CIG19 | CIG19-8 | Vaginal | Female | Full-term | Yes monozygotic with CIG18 | 102 | 14.6 | Yes | No | No | No | No |
| CIG19 | CIG19-9 | Vaginal | Female | Full-term | Yes monozygotic with CIG18 | 116 | 16.6 | Yes | No | No | No | No |
| CIG19 | CIG19-10 | Vaginal | Female | Full-term | Yes monozygotic with CIG18 | 146 | 20.9 | Yes | No | Yes | No | No |
| CIG19 | CIG19-11 | Vaginal | Female | Full-term | Yes monozygotic with CIG18 | 173 | 24.7 | Yes | No | Yes | No | No |
| CIG20 | CIG20-1 | Vaginal | Male | Full-term | No | 1 | 0.1 | Yes | No | No | No | No |
| CIG20 | CIG20-2 | Vaginal | Male | Full-term | No | 15 | 2.1 | Yes | No | No | No | No |
| CIG20 | CIG20-3 | Vaginal | Male | Full-term | No | 29 | 4.1 | Yes | No | No | No | No |
| CIG20 | CIG20-4 | Vaginal | Male | Full-term | No | 42 | 6.0 | Yes | No | No | No | No |
| CIG20 | CIG20-5 | Vaginal | Male | Full-term | No | 55 | 7.9 | Yes | No | No | No | No |

| | | | | | | | | | | | | |
|---|---|---|---|---|---|---|---|---|---|---|---|---|
| CIG20 | CIG20-6 | Vaginal | Male | Full-term | No | 81 | 11.6 | Yes | No | No | No | No |
| CIG20 | CIG20-7 | Vaginal | Male | Full-term | No | 84 | 12.0 | Yes | No | No | No | No |
| CIG20 | CIG20-8 | Vaginal | Male | Full-term | No | 96 | 13.7 | Yes | No | No | No | No |
| CIG20 | CIG20-9 | Vaginal | Male | Full-term | No | 109 | 15.6 | Yes | No | No | No | No |
| CIG20 | CIG20-10 | Vaginal | Male | Full-term | No | 142 | 20.3 | Yes | No | Yes | No | No |
| CIG21 | CIG21-1 | Vaginal | Female | Full-term | No | 6 | 0.9 | Yes | No | No | No | No |
| CIG21 | CIG21-2 | Vaginal | Female | Full-term | No | 18 | 2.6 | Yes | No | No | No | No |
| CIG21 | CIG21-3 | Vaginal | Female | Full-term | No | 31 | 4.4 | Yes | No | No | No | No |
| CIG21 | CIG21-4 | Vaginal | Female | Full-term | No | 43 | 6.1 | Yes | No | No | No | No |
| CIG21 | CIG21-5 | Vaginal | Female | Full-term | No | 55 | 7.9 | Yes | No | No | No | No |
| CIG21 | CIG21-6 | Vaginal | Female | Full-term | No | 73 | 10.4 | Yes | No | No | No | No |
| CIG21 | CIG21-7 | Vaginal | Female | Full-term | No | 82 | 11.7 | Yes | No | No | No | No |
| CIG21 | CIG21-8 | Vaginal | Female | Full-term | No | 96 | 13.7 | Yes | No | No | No | No |
| CIG21 | CIG21-9 | Vaginal | Female | Full-term | No | 113 | 16.1 | Yes | No | No | Yes | No |
| CIG21 | CIG21-10 | Vaginal | Female | Full-term | No | 134 | 19.1 | Yes | No | No | No | No |
| CIG21 | CIG21-11 | Vaginal | Female | Full-term | No | 165 | 23.6 | Yes | No | Yes | No | No |
| CIG22 | CIG22-1 | Vaginal | Female | Full-term | No | 5 | 0.7 | Yes | No | No | No | No |
| CIG22 | CIG22-2 | Vaginal | Female | Full-term | No | 14 | 2.0 | Yes | No | No | No | No |
| CIG22 | CIG22-3 | Vaginal | Female | Full-term | No | 28 | 4.0 | Yes | No | No | No | No |
| CIG22 | CIG22-4 | Vaginal | Female | Full-term | No | 41 | 5.9 | Yes | No | No | No | No |
| CIG22 | CIG22-5 | Vaginal | Female | Full-term | No | 55 | 7.9 | Yes | No | No | No | No |
| CIG22 | CIG22-6 | Vaginal | Female | Full-term | No | 69 | 9.9 | Yes | No | No | No | No |
| CIG22 | CIG22-7 | Vaginal | Female | Full-term | No | 83 | 11.9 | Yes | No | No | No | No |
| CIG22 | CIG22-8 | Vaginal | Female | Full-term | No | 93 | 13.3 | Yes | No | No | No | No |
| CIG22 | CIG22-9 | Vaginal | Female | Full-term | No | 109 | 15.6 | Yes | No | No | No | No |
| CIG22 | CIG22-10 | Vaginal | Female | Full-term | No | 137 | 19.6 | Yes | No | No | No | No |
| CIG22 | CIG22-11 | Vaginal | Female | Full-term | No | 165 | 23.6 | Yes | No | No | No | No |
| CIG23 | CIG23-1 | Vaginal | Female | Full-term | No | 3 | 0.4 | Yes | No | No | No | No |
| CIG23 | CIG23-2 | Vaginal | Female | Full-term | No | 16 | 2.3 | Yes | No | No | No | No |
| CIG23 | CIG23-3 | Vaginal | Female | Full-term | No | 29 | 4.1 | Yes | No | No | No | No |
| CIG23 | CIG23-4 | Vaginal | Female | Full-term | No | 42 | 6.0 | Yes | No | No | No | No |
| CIG23 | CIG23-5 | Vaginal | Female | Full-term | No | 56 | 8.0 | Yes | No | No | No | No |
| CIG23 | CIG23-6 | Vaginal | Female | Full-term | No | 69 | 9.9 | Yes | No | No | No | No |
| CIG23 | CIG23-7 | Vaginal | Female | Full-term | No | 83 | 11.9 | Yes | No | No | No | No |
| CIG23 | CIG23-8 | Vaginal | Female | Full-term | No | 99 | 14.1 | Yes | No | No | No | No |
| CIG23 | CIG23-9 | Vaginal | Female | Full-term | No | 114 | 16.3 | Yes | No | No | No | No |
| CIG23 | CIG23-10 | Vaginal | Female | Full-term | No | 152 | 21.7 | Yes | No | No | No | No |
| CIG23 | CIG23-11 | Vaginal | Female | Full-term | No | 176 | 25.1 | Yes | No | Yes | No | No |
| CIG24 | CIG24-1 | Vaginal | Female | Full-term | No | 2 | 0.3 | Yes | No | No | No | No |
| CIG24 | CIG24-2 | Vaginal | Female | Full-term | No | 16 | 2.3 | Yes | No | No | No | No |
| CIG24 | CIG24-3 | Vaginal | Female | Full-term | No | 30 | 4.3 | Yes | No | No | No | No |
| CIG24 | CIG24-4 | Vaginal | Female | Full-term | No | 43 | 6.1 | Yes | No | No | No | No |
| CIG24 | CIG24-5 | Vaginal | Female | Full-term | No | 58 | 8.3 | Yes | No | No | No | No |
| CIG24 | CIG24-6 | Vaginal | Female | Full-term | No | 71 | 10.1 | Yes | No | No | No | No |
| CIG24 | CIG24-7 | Vaginal | Female | Full-term | No | 84 | 12.0 | Yes | No | No | No | No |
| CIG24 | CIG24-8 | Vaginal | Female | Full-term | No | 100 | 14.3 | Yes | No | No | No | No |
| CIG24 | CIG24-9 | Vaginal | Female | Full-term | No | 111 | 15.9 | Yes | No | No | No | No |
| CIG24 | CIG24-10 | Vaginal | Female | Full-term | No | 139 | 19.9 | Yes | No | No | Yes | No |
| CIG24 | CIG24-11 | Vaginal | Female | Full-term | No | 170 | 24.3 | Yes | No | No | No | No |
| CIG25 | CIG25-1 | Vaginal | Female | Full-term | No | 4 | 0.6 | Yes | No | No | No | No |
| CIG25 | CIG25-2 | Vaginal | Female | Full-term | No | 18 | 2.6 | Yes | No | No | No | No |
| CIG25 | CIG25-3 | Vaginal | Female | Full-term | No | 30 | 4.3 | Yes | No | No | No | No |
| CIG25 | CIG25-4 | Vaginal | Female | Full-term | No | 46 | 6.6 | Yes | No | No | No | No |
| CIG25 | CIG25-5 | Vaginal | Female | Full-term | No | 70 | 10.0 | Yes | No | No | No | No |
| CIG25 | CIG25-6 | Vaginal | Female | Full-term | No | 86 | 12.3 | Yes | No | No | Yes | No |
| CIG25 | CIG25-7 | Vaginal | Female | Full-term | No | 100 | 14.3 | Yes | No | No | Yes | No |
| CIG25 | CIG25-8 | Vaginal | Female | Full-term | No | 119 | 17.0 | Yes | No | No | Yes | No |
| CIG25 | CIG25-9 | Vaginal | Female | Full-term | No | 127 | 18.1 | Yes | No | No | Yes | No |
| CIG25 | CIG25-10 | Vaginal | Female | Full-term | No | 141 | 20.1 | Yes | No | Yes | Yes | No |
| CIG25 | CIG25-11 | Vaginal | Female | Full-term | No | 153 | 21.9 | Yes | No | Yes | Yes | No |

**Supplementary Data 2b. Bacterial taxa assignment in Copenhagen Infant Gut study**

| OTU Ids | Relative abundance (%) | Prevalence (%) | Top BLAST hit# [Identity %] | GreenGenes taxonomy | Assinged taxonomy |
|---|---|---|---|---|---|
| OTU_1 | 38.5 | 92 | Bifidobacterium longum 100% | g__Bifidobacterium; s__longum | *B. longum* |
| OTU_2 | 9.1 | 55 | Bifidobacterium breve 100% | g__Bifidobacterium; s__breve | *B. breve* |
| OTU_3 | 7.9 | 75 | Bifidobacterium bifidum 100% | g__Bifidobacterium; s_bifidum | *B. bifidum* |
| OTU_4 | 6.4 | 49 | Bifidobacterium catenulatum 100%; Bifidobacterium pseudocatnulatum 100%; Bifidobacterium kashiwanohense 100% | g__Bifidobacterium | *B. catenulatum* group |
| OTU_5 | 5.5 | 39 | Clostridium neonatale 100% | g__Clostridium; s__neonatale | *C. neonatale* |
| OTU_9/132 | 4.4 | 82 | Escherichia coli 100% / Escherichia coli 99% | g__Escherichia; s__coli/f__Enterobacteriaceae | *E. coli* |
| OTU_17/24/246/349/446 | 2.4 | 34 | Klebsiella pneumoniae 100% / Klebsiella variicola 99% / Klebsiella pneumoniae 99% / Klebsiella variicola 99% / Klebsiella pneumoniae 97% | g__Klebsiella; s__/g__Klebsiella; s__/g__Klebsiella; s__/g__Klebsiella; s__/g__Klebsiella; s__ | *Klebsiella* spp. |
| OTU_8 | 2.0 | 33 | Ruminococcus gnavus 100% | g__[Ruminococcus]; s__gnavus | *R. gnavus* |
| OTU_10 | 1.9 | 92 | Streptococcus salivarius 100% | g__Streptococcus; s__ | *S. salivarius* |
| OTU_7 | 1.6 | 24 | Bifidobacterium dentium 99% | g__Bifidobacterium; s__ | *B. dentium* |
| OTU_25 | 1.5 | 49 | multiple Enterococcus spp. 100% | g__Enterococcus; s__ | *Enterococcus* spp. |
| OTU_6 | 1.4 | 24 | Erysipeloclostridium ramosum 100% | f__Erysipelotrichaceae; g__; s__ | *E. ramosum* |
| OTU_13 | 1.4 | 15 | Veillonella ratti 100% | g__Veillonella; s__ | *V. ratti* |
| OTU_21/26/40 | 1.3 | 37 | Bacteroides dorei 99% / Bacteroides vulgatus 98% / Bacteroides vulgatus 99% | g__Bacteroides; s__/g__Bacteroides; s__/g__Bacteroides; s__ | *B. dorei/vulgatus* |
| OTU_18/224/237/424 | 1.3 | 34 | Parabacteroides diastonis 99% / Parabacteroides diastonis 98% / Parabacteroides diastonis 97% / Parabacteroides diastonis 97% | g__Parabacteroides; s__distasonis/g__Parabacteroides; s__distasonis/g__Parabacteroides; s__distasonis/g__Parabacteroides; s__distasonis | *P. diastonis* |
| OTU_12 | 1.2 | 22 | C. hathewayi 100% | f__Lachnospiraceae | *C. hathewayi* |
| OTU_11 | 1.1 | 27 | C. aerofaciens 100% | g__Collinsella; s__aerofaciens | *C. aerofaciens* |
| OTU_33/37/120/320 | 1.0 | 47 | multiple Clostridium spp. 100% / Clostridium paraputrificum 100% / multiple Clostridium spp. 99% / multiple Clostridium spp. 99% | f__Clostridiaceae; g__SMB53; s__/f__Clostridiaceae; g__Clostridium/f__Clostridiaceae/f__Clostridiaceae; g__Clostridium; s__botulinum | *Clostridium* spp. |
| OTU_131/244 | 0.9 | 25 | multiple Lachnospiraceae spp. 100% | f__Lachnospiraceae/f__Lachnospiraceae | *Lachnospiraceae* spp. |
| OTU_23/31 | 0.8 | 95 | multiple Streptococcus spp. 99% /multiple Streptococcus spp. 100% | g__Streptococcus; s__/g__Streptococcus; s__infantis | *Streptococcus* spp. |
| OTU_16 | 0.7 | 79 | Staphylococcus epidermidis 100% | g__Staphylococcus; s__epidermidis | *S. epidermidis* |
| OTU_32/219 | 0.6 | 71 | Veillonella dispar 99% / Veilonella parvula 99%; Veillonella dispar 99% | g__Veillonella; s__dispar/g__Veillonella; s__dispar | *V. dispar/parvula* |
| OTU_14 | 0.6 | 50 | Lactobacillus rhamnosus 100%; Lactobacillus casei 100%; Lactobacillus paracasei 100% | g__Lactobacillus; s__zeae | *L. rhamnosus/casei/paracasei* |
| OTU_19 | 0.5 | 53 | Enterococcus faecalis 100% | g__Enterococcus; s__ | *E. faecalis* |
| OTU_20 | 0.4 | 18 | Bacteroides fragilis 100% | g__Bacteroides; s__fragilis | *B. fragilis* |
| OTU_15 | 0.4 | 17 | Actinomyces urogenitalis 100% | g__Actinomyces; s__ | *A. urogenitalis* |
| OTU_27 | 0.4 | 29 | Clostridium perfringens 100% | g__Clostridium; s__perfringens | *C. perfringens* |
| OTU_159 | 0.4 | 63 | Staphylococcus aureus 100% | g__Staphylococcus; s__aureus | *S. aureus* |
| OTU_29 | 0.3 | 11 | Citrobacter freundii 100% | g__Citrobacter; s__ | *C. freundii* |
| OTU_28 | 0.2 | 5 | Bifidobacterium scardovii 99% | g__Bifidobacterium; s__ | *B. scardovii* |
| OTU_22 | 0.2 | 15 | Bacteroides thetaiotaomicron 100% | g__Bacteroides; s__ovatus | *B. thetaiotaomicron* |
| OTU_443 | 0.2 | 46 | Staphylococcus saprophyticus 100% | g__Staphylococcus; s__saprophyticus | *S. saprophyticus* |
| OTU_35 | 0.2 | 23 | Propionibacterium acne 100% | g__Propionibacterium; s__ | *P. acne* |
| OTU_30 | 0.2 | 42 | Eggertella lenta 100% | g__Eggerthella; s__lenta | *E. lenta* |
| OTU_174 | 0.2 | 8 | Bifidobacterium adolescentis 100% | g__Bifidobacterium; s__adolescentis | *B. adolescentis* |
| OTU_141 | 0.1 | 22 | Enterobacter cloacae 99% | g__Enterobacter; s__cloacae | *E. cloacae* |
| OTU_43 | 0.1 | 11 | Blautia obeum 100%; Blautia luti 100%; Blautia wexlerae 100% | g__Blautia; s__obeum | *Blautia obeum/luti/wexlerae* |
| OTU_34 | 0.1 | 7 | multiple Enterobacteriaceae spp. 100% | f__Enterobacteriaceae | *Enterobacteriaceae spp.* |
| OTU_47 | 0.1 | 6 | Bifidobacterium animalis 100%; Bifidobacterium pseudolongum 100% | g__Bifidobacterium; s__pseudolongum | *B. animalis/pseudolongum* |

#BLAST against the 16s rRNA Database at NCBI

**Supplementary Data 2c. Species-level taxonomic assignment of Bifidobacterium OTUs**

| OTU ID# | GreenGenes taxonomy [RDP classifier 0.5] | 1st BLAST hit | | | | | 2nd BLAST HIT | | | | Assigned taxonomy |
|---|---|---|---|---|---|---|---|---|---|---|---|
| | | % Identity | E-value | Query coverage | Gaps | Mismatch | % Identity | Query coverage | Gaps | Mismatch | |
| OTU_1 | Bifidobacterium longum | 100% B. longum subsp. infantis | 9.00E-72 | 100% | 0 | 0 | 99% B. longum subsp. suis | 100% | 0 | 1 | ***B. longum*** |
| | | 100% B. longum subsp. suillum | 9.00E-72 | 100% | 0 | 0 | 99% B. longum subsp. longum | 100% | 0 | 1 | |
| OTU_2 | Bifidobacterium breve | 100% B. breve | 1.00E-74 | 100% | 0 | 0 | 95% B. bifidum | 100% | 3 | 4 | ***B. breve*** |
| OTU_3 | Bifidobacterium bifidum | 100% B. bifidum | 7.00E-73 | 100% | 0 | 0 | 97% B. boum | 100% | 2 | 3 | ***B. bifidum*** |
| OTU_4 | Bifidobacterium; s__ | 100% B. catenulatum | 7.00E-73 | 100% | 0 | 0 | 98% B. angulatum | 100% | 0 | 3 | ***B. catenulatum group*** |
| | | 100% B. pseudocatenulatum | 7.00E-73 | 100% | 0 | 0 | | | | | |
| | | 100% B. kashiwanohense | 7.00E-73 | 100% | 0 | 0 | | | | | |
| OTU_7 | Bifidobacterium; s__ | 99% B. dentium | 2.00E-72 | 100% | 0 | 1 | 99% B. boum | 100% | 0 | 2 | ***B. dentium*** |
| OTU_28 | Bifidobacterium; s__ | 99% B. scardovii | 3.00E-71 | 100% | 0 | 1 | 99% B. aerophillum | 100% | 0 | 2 | ***B. scardovii*** |
| OTU_174 | Bifidobacterium; s__adolescentis | 99% B. adolescentis | 9.00E-72 | 100% | 1 | 0 | 97% B. boum | 100% | 1 | 4 | ***B. adolescentis*** |
| | | 99% B. faecale | 9.00E-72 | 100% | 1 | 0 | | | | | |
| | | 99% B. stercoris | 9.00E-72 | 100% | 1 | 0 | | | | | |
| | | 99% B. ruminantium | 9.00E-72 | 100% | 1 | 0 | | | | | |
| OTU_47 | Bifidobacterium; s__pseudolongum | 100% B. pseudolongum subsp. pseudolongum | 2.00E-78 | 100% | 0 | 0 | 99% B. animalis subsp. animalis | 100% | 1 | 1 | ***B. pseudolongum/animalis*** |
| | | 100% B. pseudolongum subsp. globosum | 2.00E-78 | 100% | 0 | 0 | | | | | |
| | | 100% B. animalis subsp. lactis | 2.00E-78 | 100% | 0 | 0 | | | | | |

#Only OTUs representing relative abundance >0.1% within the total relative abundance of *Bifidobacterium* genus were included
